# Improving the secretion of designed protein assemblies through negative design of cryptic transmembrane domains

**DOI:** 10.1101/2022.08.04.502842

**Authors:** Jing Yang (John) Wang, Alena Khmelinskaia, William Sheffler, Marcos C. Miranda, Aleksandar Antanasijevic, Andrew J. Borst, Susana Vazquez Torres, Chelsea Shu, Yang Hsia, Una Nattermann, Daniel Ellis, Carl Walkey, Maggie Ahlrichs, Sidney Chan, Alex Kang, Hannah Nguyen, Claire Sydeman, Banumathi Sankaran, Mengyu Wu, Asim K. Bera, Lauren Carter, Brooke Fiala, Michael Murphy, David Baker, Andrew B. Ward, Neil P. King

## Abstract

Computationally designed protein nanoparticles have recently emerged as a promising platform for the development of new vaccines and biologics. For many applications, secretion of designed nanoparticles from eukaryotic cells would be advantageous, but in practice they often secrete poorly. Here we show that designed hydrophobic interfaces that drive nanoparticle assembly are often predicted to form cryptic transmembrane domains, suggesting that interaction with the membrane insertion machinery could limit efficient secretion. We develop a general computational protocol, the Degreaser, to design away cryptic transmembrane domains without sacrificing protein stability. Retroactive application of the Degreaser to previously designed nanoparticle components and nanoparticles considerably improves secretion, and modular integration of the Degreaser into design pipelines results in new nanoparticles that secrete as robustly as naturally occurring protein assemblies. Both the Degreaser protocol and the novel nanoparticles we describe may be broadly useful in biotechnological applications.

## Introduction

Secreted proteins make up nearly 20% of the human proteome, and are the primary medium for intercellular communication in animals (Uhlén et al. 2019; Farhan and Rabouille 2011). Due to their potent and wide-ranging functions, many secreted proteins such as antibodies, hormones, cytokines, and growth factors are of great interest for therapeutic applications. Furthermore, secreted and membrane-anchored proteins from pathogens are common targets for prophylactic or therapeutic interventions in infectious disease. Secretion from eukaryotic cells is required for the recombinant production of many protein biologics, as they often feature secretory pathway-specific post-translational modifications such as proteolytic cleavage (Braun and Sauter 2019), glycosylation (Ohtsubo and Marth 2006), or disulfide bond formation (Wittrup 1995). Understanding and controlling the secretion of a protein of interest is thus mandatory for optimal development of secreted protein technologies.

Several strategies for increasing the yield of secreted proteins have been established. Most approaches have focused on engineering or adapting cell lines for secretion. Chinese hamster ovary (CHO) cells are by far the most widely used for the production of antibodies and other secreted biologics, accounting for 84% of approved monoclonal biopharmaceuticals from 2015-2018 (Walsh 2018). Genomic, transcriptomic, and proteomic profiling of this cell line has yielded insights into the adaptations that confer its ability to grow at high density and robustly secrete target proteins, and has also identified opportunities for intentional improvements through cell engineering (Baycin-Hizal et al. 2012; Rupp et al. 2018; Kuo et al. 2018; X. Xu et al. 2011). For example, overexpression of secretory chaperones or other secretory pathway factors can significantly increase the secreted yield of target proteins (Xiao, Shiloach, and Betenbaugh 2014; Le Fourn et al. 2014). Engineering other eukaryotic expression hosts, such as *S. cerevisiae* or *P. pastoris*, has provided further avenues for recombinant expression (Vieira Gomes et al. 2018). However, host-directed efforts like these are agnostic to the sequence of the target protein and therefore do not take into account sequence-specific factors that could affect secretion.

An alternative approach that is more customizable and amenable to downstream biological applications, in particular when the host cell cannot be modified, is engineering the sequences of protein biologics themselves. For example, computational or experimental evolution of signal peptides in both yeast and mammalian cells have yielded portable sequences that maximize secretion (Rakestraw et al. 2009; Güler-Gane et al. 2016; Kober, Zehe, and Bode 2013). The introduction (Sagt et al. 2000; Liu et al. 2009) or removal (Dalvie, Brady, et al. 2021) of potential N-linked glycosylation sites has also been shown to substantially affect protein secretion. Importantly, sequence optimization via rational design or evolutionary methods can maximize production of a protein while still preserving its biological function, such as immunogenicity or receptor binding (J. Jardine et al. 2013; Rojas et al. 2019)). Although powerful, these methods tend to focus on the maximization of secreted yield for single proteins and can be laborious, underscoring the need for general methods applicable to broad classes of secreted proteins. To devise such general methods, it is necessary to understand the effects of common sequence elements on protein secretion and how they can be modulated through design.

Protein translocation across the endoplasmic reticulum (ER) membrane is a key step in the traversal of proteins through the eukaryotic secretory pathway. Transmembrane, secreted, and ER/Golgi resident proteins typically contain an N-terminal signal peptide that is recognized by the signal recognition particle (SRP) (Akopian et al. 2013; von Heijne 1990; Egea, Stroud, and Walter 2005). The SRP-polypeptide-ribosome complex is then targeted to the ER membrane, across which the nascent polypeptide is translocated in a mostly-unfolded state through a protein channel, the Sec translocon. The translocon complex senses the hydrophobicity of segments within a polypeptide to determine its fate: highly hydrophobic segments are partitioned into the ER membrane, while segments of low hydrophobicity are translocated into the ER lumen (Rapoport 2007). To predict this biological hydrophobicity sensing, Hessa *et al*. empirically determined the position-specific amino acid contribution of each residue in a model polypeptide segment to transmembrane insertion potential (dG_ins_) and generated a theoretical model that can, using sequence alone, predict the dG_ins_ of any given polypeptide segment (Hessa et al. 2005, 2007)dG_ins,pred_; _(Hessa et al. 2005, 2007)_. Unsurprisingly, the model predicts that transmembrane proteins contain segments of low dG_ins,pred_ while secretory proteins are generally devoid of such segments. Interestingly, many proteins that are typically expressed in the prokaryotic or eukaryotic cytoplasm also contain segments of low dG_ins,pred_, as they have not evolved under selective pressure to avoid them. We hypothesize that, when expressed recombinantly, the Sec translocon may interpret segments of low dG_ins,pred_ within proteins targeted for secretion as transmembrane domains, leading to membrane insertion instead of efficient secretion.

Protein nanoparticles, which in nature evolved to function as containers, reaction chambers, and multivalent display platforms (Abrescia et al. 2012; Bobik, Lehman, and Yeates 2015; Goodsell and Olson 2000), have long been a promising biotechnological platform (S. Zhang 2003; Padilla, Colovos, and Yeates 2001; Douglas and Young 2006). Virus-like particles (VLPs) and other naturally occurring protein nanoparticles such as ferritin and lumazine synthase have been engineered or evolved to encapsulate a number of different cargoes, such as inorganic particles, polymers, micelles, enzymes, DNA, and small molecules (de Ruiter et al. 2019; Edwardson, Tetter, and Hilvert 2020; Tetter et al. 2021). Self-assembling protein complexes have also been used as scaffolds for nanoparticle vaccine design, an application that often requires secretion from eukaryotic cells to obtain natively folded antigens with appropriate post-translational modifications (Kanekiyo et al. 2013; J. Jardine et al. 2013; López-Sagaseta et al. 2016). In the last few years, new and accurate computational methods have made possible the design of novel self-assembling proteins with customized structures (King et al. 2012; Khmelinskaia, Wargacki, and King 2021; Ueda et al. 2020a; King et al. 2014; Hsia et al. 2016, 2021). These methods allow rational exploration of structural and functional space beyond the limited set of protein nanoparticle architectures sampled during evolution (Huang, Boyken, and Baker 2016). Additionally, because they are explicitly designed to adopt low-energy states, designed protein nanoparticles are often hyperstable, a property that can enhance the stability of proteins fused to them (Bruun et al. 2018; Marcandalli et al. 2019; He et al. 2021). The recent entry of multiple vaccines based on computationally designed nanoparticles into clinical trials, and the regulatory approval of one of them for COVID-19 (Song et al. 2022), highlights the potential technological impact of this promising class of proteins (Marcandalli et al. 2019; Walls et al. 2020; Boyoglu-Barnum et al. 2021).

Naturally occurring and designed protein nanoparticles alike use hydrophobic interactions both in the folding of their subunits as well as in the interactions between subunits that drive supramolecular assembly (Janin, Bahadur, and Chakrabarti 2008; Khmelinskaia, Wargacki, and King 2021). One approach to protein nanoparticle design computationally docks symmetric protein building blocks into a desired geometry and then designs novel protein-protein interfaces between them (King et al. 2012, 2014). This design strategy has been previously shown to generate robust, well-expressed, stable protein nanoparticles constructed from subunits or pairs of subunits that are most often expressed in the cytoplasm of bacterial cells (King et al. 2012, 2014; Bale et al. 2016; Ueda et al. 2020a; Hsia et al. 2016; Sahasrabuddhe et al. 2018). Nonetheless, only a small subset of the components of these designed nanoparticles have been robustly secreted from eukaryotic cells (Marcandalli et al. 2019; Walls et al. 2020; Boyoglu-Barnum et al. 2021).

Here, we develop a method to improve the secretion of computationally designed protein nanoparticles. First, we identify a correlation between the presence of cryptic transmembrane domains and low levels of secreted protein. We then develop a general computational protocol, named the Degreaser, that specifically designs away cryptic transmembrane domains without sacrificing overall structural stability. We demonstrate the ability of the Degreaser to not only retroactively improve the secreted yield of existing nanoparticles and nanoparticle components, but also to avoid the introduction of cryptic transmembrane domains during the design of a new set of robustly secreting designed protein nanoparticles.

## Results

### The Degreaser

During the course of our work designing novel self-assembling protein nanomaterials and using them as scaffolds for heterologous antigen display, we have observed that many (but not all (Boyoglu-Barnum et al. 2021; Walls et al. 2020)) designed nanoparticles and nanoparticle components secrete from eukaryotic cells at very low levels. We hypothesized that their inefficient secretion may derive from the introduction of long contiguous stretches of hydrophobic amino acids during nanoparticle interface design. These stretches could be interpreted by the Sec translocon as cryptic transmembrane domains, leading to inefficient translocation across the ER membrane and poor protein secretion. Using the model of Hessa et al. (Hessa et al. 2005, 2007) and a sliding window of 19 amino acids, we compared the distributions of lowest dG_ins,pred_ per sequence of 489 computationally designed two-component icosahedral nanoparticle scaffolds (Bale et al. 2016) to those of 309 transmembrane and 585 non-transmembrane proteins in the RCSB Protein Databank (Figure 1a) (Burley et al. 2021; UniProt Consortium 2021). We identified a dG_ins,pred_ threshold of +2.7 kcal/mol that maximally discriminated between natural transmembrane and non-transmembrane proteins (84.7% and 15.6% < +2.7 kcal/mol, respectively). 61.3% of the protein nanoparticle components contained segments below this threshold, indicating that the designed proteins differ substantially from naturally occurring non-transmembrane proteins in their likelihood to include hydrophobic segments. As a preliminary experimental test of whether cryptic transmembrane domains affect protein secretion, we analyzed the expression of a set of 480 *de novo* designed peptide-binding proteins by yeast surface display (Boder and Wittrup 1997). To reach the cell surface, these proteins must be translocated across the ER membrane and into the lumen. We found that the presence of segments of low dG_ins,pred_ strongly hindered surface display, while proteins lacking such cryptic transmembrane domains were generally efficiently exported to the cell surface (Supplementary Figure S1).

**Figure 1.**
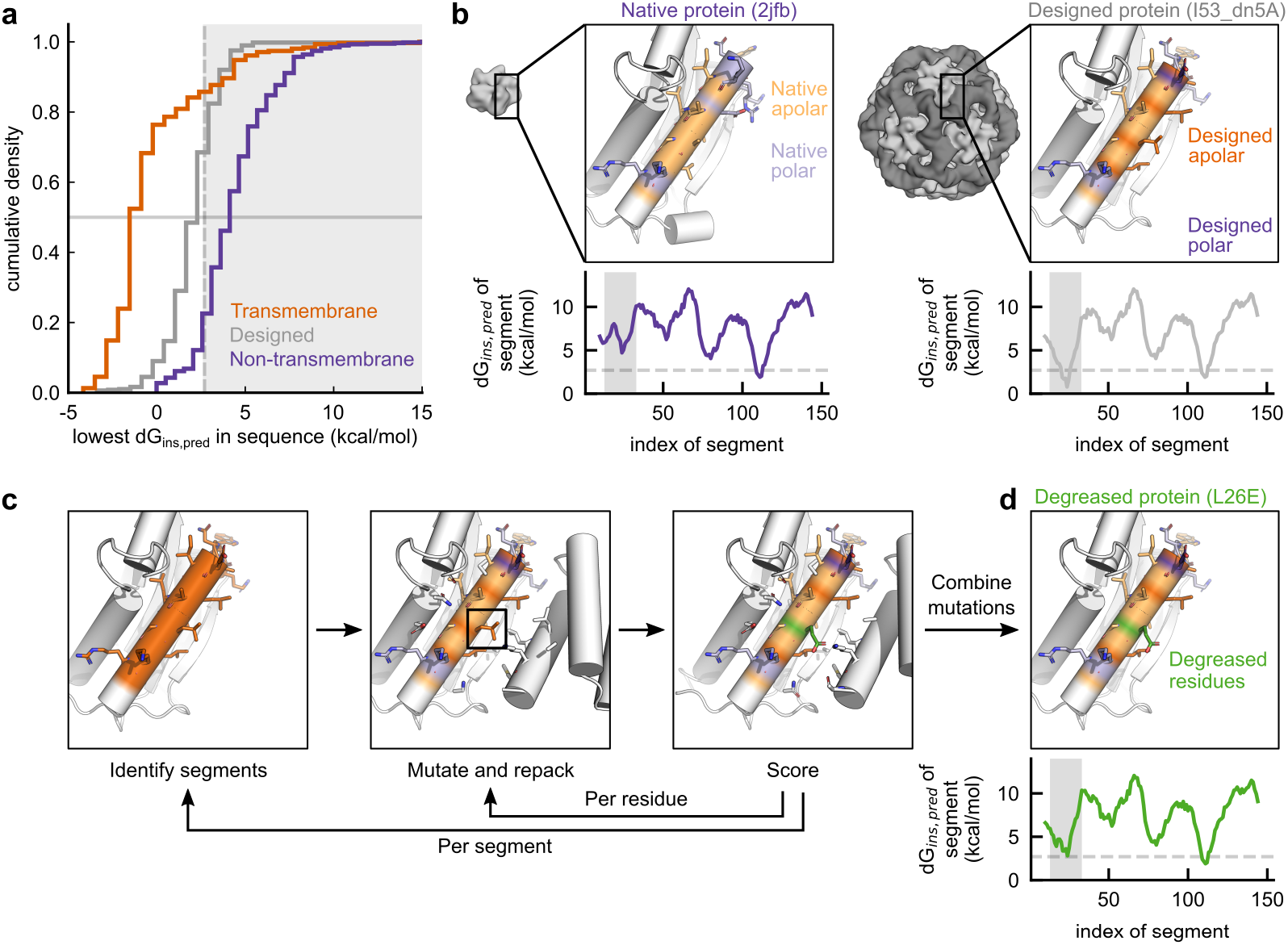
Graphical depiction of the Degreaser protocol. (a) Designed proteins have segments of lower dG_ins,pred_ than do non-transmembrane proteins, but higher than those of transmembrane proteins. The shaded region indicates dG_ins,pred_ > +2.7 kcal/mol. (b) *Top*, Protein building blocks have surface polar residues (light and dark purple) that are mutated to apolar residues (light and dark orange) during nanoparticle interface design. Alpha helices are depicted as cylinders, and side chains of residues within the identified segment or within 8 Å of the designed interface are shown as sticks. *Bottom,* These mutations often decrease the lowest dG_ins,pred_ segment in the sequence (lowest region of designed protein shaded gray, horizontal dashed line at +2.7 kcal/mol). (c) The Degreaser iteratively identifies hydrophobic segments, perturbs each residue within such segments through point mutations, and chooses the optimal variant after neighborhood repacking. (d) Degreased proteins feature one or more polar mutations (green) that preserve the originally designed interface while increasing dG_ins,pred_.

Inspection of individual designed nanoparticle components highlighted the impact that interface design can have on dG_ins,pred_. For example, docking naturally occurring pentameric (PDB 2jfb) and computationally designed trimeric (PDB 5hrz) building blocks and designing them to form an icosahedral nanoparticle (I53_dn5; (Ueda et al. 2020b)) resulted in two hydrophilic to hydrophobic substitutions in a single helix of the pentamer (Figure 1b). As a result, the dG_ins,pred_ at index 14 (residues 14-32) of the pentamer (I53_dn5A) was reduced from +4.71 kcal/mol to +0.779 kcal/mol, suggesting that this region of the protein may be interpreted as a cryptic transmembrane domain during translocation. Quantification of secreted protein by western blot revealed a nearly 100-fold reduction in secretion of I53_dn5A relative to its naturally occurring counterpart (Figure 2a and Supplementary Figure S2). Inspection of several other previously designed nanoparticle components that secrete with varying efficiency from eukaryotic cells supported the relationship between the presence of cryptic transmembrane domains and poor secretion (Supplementary Figure S2). In each poorly secreted case, cryptic transmembrane domains resulted from the introduction of a few surface-exposed hydrophobic residues at the designed interface that, combined with existing hydrophobic residues involved in packing within the protein itself, form segments of low transmembrane insertion potential. Together, these observations suggest that the secretion of designed proteins from eukaryotic cells, a prerequisite for several biotechnological applications, may be limited by the presence of cryptic transmembrane domains introduced during design.

**Figure 2.**
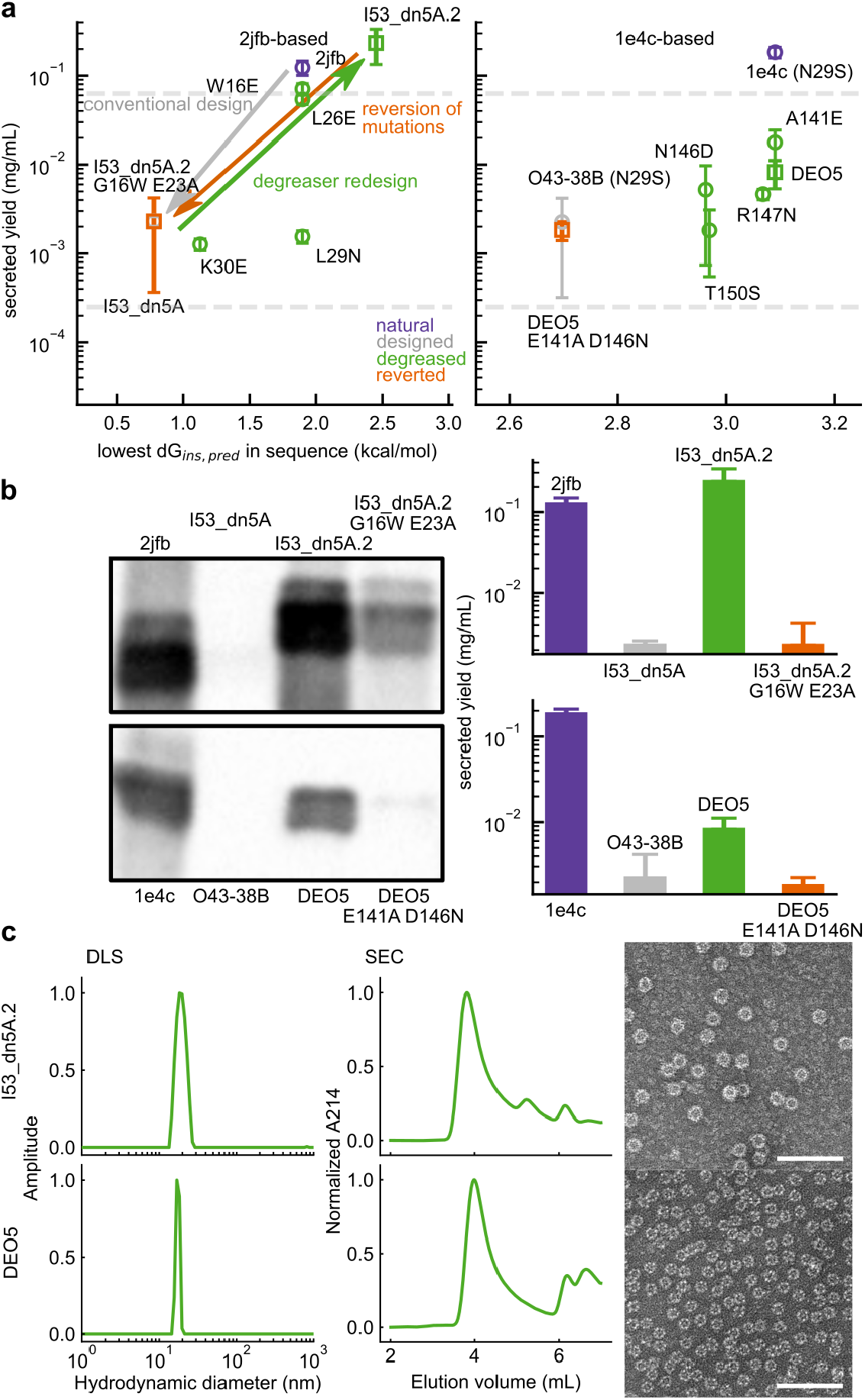
Retroactive Degreasing of protein nanoparticle components improves secretion yield. (a) To assess secreted yield of previously designed proteins and their natural counterparts, material from transiently transfected cell cultures was fractionated and analyzed by Western blot. Horizontal dashed lines represent the upper and lower limits of the linear range of a standard curve generated by densitometry. Square markers represent variants that incorporate mutations from parallel redesign for improvement of other phenotypes. (b) Representative Western blots of cell culture supernatants of two design trajectories (naturally occurring scaffold, purple; original design, gray; Degreased re-design, green; reversion of Degreaser mutations, orange). (c) Characterization of *in vitro* assemblies of secreted I53_dn5A.2 (*top row*) and DEO5 (*bottom row*) with their respective second components I53_dn5B and O43-38A, purified from bacterial cells, reveal nanoparticle sizes and morphologies indistinguishable to those of bacterially-produced I53_dn5 (Ueda et al. 2020b) and O43-38 (Supplementary Figure S3), respectively. Scale bar, 100 nm.

To address this issue, we developed the Degreaser, a new design protocol in the Rosetta macromolecular modeling software suite (Leaver-Fay et al. 2011; Leman et al. 2020) that detects and eliminates cryptic transmembrane domains without disrupting protein structure or stability. We had three primary goals for the protocol: (1) accurate identification of regions of low dG_ins,pred_ and their improvement through minimal sequence perturbations, (2) preservation of the overall integrity of the protein’s structure, and (3) compatibility with large-scale protein design protocols. The Degreaser uses an internal implementation of the model of Hessa et al. (Hessa et al. 2005, 2007) to identify 19-residue segments of input protein structures that have dG_ins,pred_ < 3.5 kcal/mol. Each residue in these segments is individually mutated to polar amino acids selected from a user-specified set, the mutated residue and nearby residues in three-dimensional space are repacked to attempt to accommodate the mutation, and the overall Rosetta score and dG_ins,pred_ are re-evaluated (Figure 1c). All chains in the input structure are modeled throughout the process, including symmetry-related subunits when appropriate, to capture the potential energetic effects of each mutation in the correct structural context. Mutations are discarded if they fail to increase dG_ins,pred_ by a user-specified amount or are energetically unfavorable (i.e., result in an increase in Rosetta score beyond a user-specified threshold), usually due to steric clashes. Designs that pass these filters (for details, see Materials and Methods) are output as potential candidates for experimental evaluation. We found that a small number of mutations was often sufficient to substantially raise the dG_ins,pred_ of the segment of interest. For example, the dG_ins,pred_ of residues 14-32 in the I53_dn5 pentamer increased from +0.779 to +2.812 kcal/mol with only one mutation to a charged amino acid (Figure 1d).

### Improving the secretion yield of existing designed protein nanoparticle components

As a first test of our protocol, several previously designed two-component nanoparticle proteins that were originally expressed in bacteria were screened for secretion from Expi293F cells (Bale et al. 2016; Ueda et al. 2020b). Three had no predicted segments of low dG_ins,pred_, needing no Degreaser application, and secreted with yields between 3 and 34 μg/mL of cell culture supernatant (Supplementary Figure S2). However, seven others yielded between 0.6 and 3 μg/mL, precluding purification and *in vitro* assembly experiments. In two of the latter cases, the Degreaser was applied to create several single-residue variants intended to improve secretion of I53_dn5A, the pentameric component of the icosahedral nanoparticle I53_dn5 (Ueda et al. 2020b), and O43-38B, the tetrameric component of a designed octahedral nanoparticle, O43-38 (Supplementary Figure S3a), respectively. The two naturally occurring proteins from which these nanoparticle components were derived (including an N29S mutation in 1e4c to knock out a potential N-linked glycosylation site) both secreted at high levels, around 100 μg/mL (Figure 2a). The Degreased variants were predicted to have no significant effects on protein stability, as indicated by the <5% change in Rosetta score compared to the input structures (Supplementary Figure S4). In each case, variants with significant improvements in secretion yield were obtained. For example, two of four I53_dn5A pentamer variants tested, W16E and L26E, increased dG_ins,pred_ by more than 1.1 kcal/mol and improved secretion yield more than twenty-fold (Figure 2a; amino acid sequences of all novel proteins used in this study are provided in Supplementary Table S1). For the O43-38B tetramer, two single-residue variants, A141E and R147N, boosted secretion eight-fold and two-fold, respectively.

To assess the orthogonality and compatibility of the Degreaser with unrelated mutations, some nanoparticle components were further redesigned for other purposes. Because existing methods for designing protein stability, solubility, or expression do not explicitly penalize the introduction of contiguous hydrophobic segments, the variants generated were generally orthogonal to those of the Degreaser. In the case of I53_dn5A, we mutated a pair of cysteine residues retained from the naturally occurring pentameric scaffold to alanine to prevent the formation of off-target disulfide bonds. Ten additional nearby mutations and a three-residue deletion were also included to maintain protein stability. These unrelated mutations were dG_ins,pred_-agnostic and did not substantially change the hydrophobicity of already-hydrophobic segments within the protein (Supplementary Figure S5). In these contexts, the Degreaser mutations again positively impacted protein secretion without negatively impacting protein assembly: the stabilized O43-38B variant DEO5 (which incorporated two Degreaser mutations: A141E and N146D) and the I53_dn5A.2 variant (incorporating the A23E Degreaser mutation) had higher secretion yield than their originally designed counterparts (8.2 and 234 μg/mL, respectively; Figure 2a and b). To confirm that the secretion phenotype was dependent on the Degreaser-identified mutations, we reverted these positions to their originally designed identities, which resulted in substantial decreases in dG_ins,pred_ and secretion yield as expected (1.8 and 2.3 μg/mL, respectively; orange data points and bars). Importantly, these nanoparticle components retained their ability to assemble *in vitro* (Figure 2c), as validated by dynamic light scattering (DLS), size exclusion chromatography (SEC), and negative stain electron microscopy (nsEM) of *in vitro* assembly reactions containing their respective second components. Taken together, these results demonstrate that the Degreaser can be used to generate nanoparticle component variants with improved secretion that retain their ability to assemble to designed nanoparticle architectures.

### Improving the secretion yield of protein nanoparticles by degreasing cryptic transmembrane domains

Encouraged by the ability of the Degreaser to improve secretion of the oligomeric components of designed two-component protein nanoparticles, we next evaluated whether it could also improve the secretion of one-component or homomeric nanoparticles that self-assemble during secretion. Several naturally occurring homomeric protein nanoparticles such as lumazine synthase and ferritin have been explored in biotechnological applications such as nanoparticle vaccine design and drug delivery (López-Sagaseta et al. 2016; Palombarini et al. 2020). We compared the secretion of these two naturally occurring protein nanoparticles to the computationally designed one-component icosahedral nanoparticle I3-01 (Hsia et al. 2016) and found that they secrete at levels roughly ten- and one hundred-fold higher, respectively (Figure 3a). This observation is consistent with the lack of low dG_ins,pred_ segments in lumazine synthase and ferritin, whereas I3-01 has two segments of low dG_ins,pred_, one near the N terminus and a second near the C terminus, both of which include polar to hydrophobic mutations at the designed nanoparticle interface (Figure 3b).

**Figure 3.**
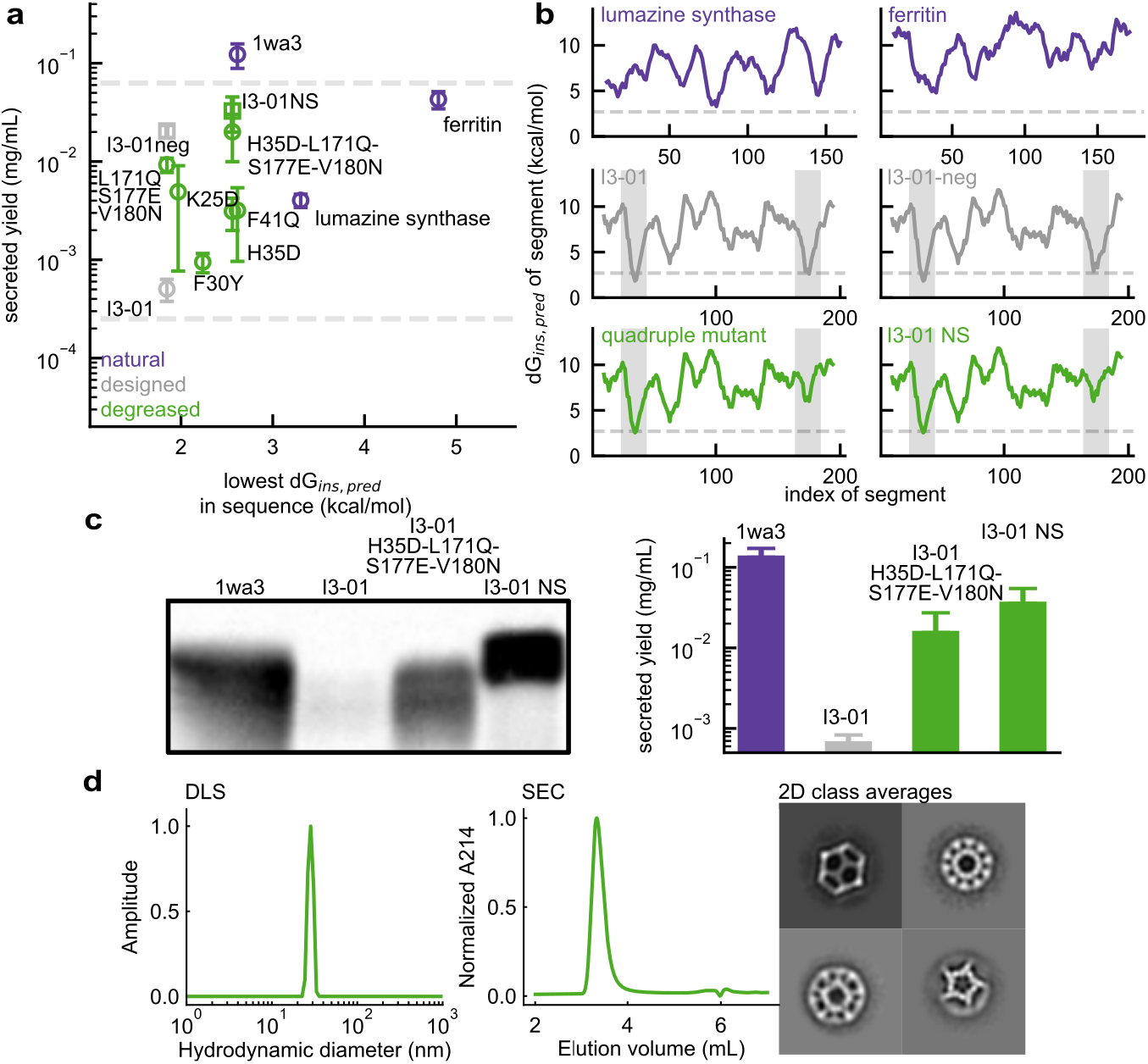
Retroactive Degreasing of a designed protein nanoparticle improves secretion yield. (a) I3-01 can be Degreased to boost secretion yield; square markers represent variants that incorporate mutations from orthogonal redesign for improvement of other phenotypes. (b) Comparison of the dG_ins,pred_ per segment of wild-type (gray) and best-secreted (green) variants of I3-01 with secreted natural protein assemblies (purple). Lowest regions of wild-type I3-01 are shaded in gray, and the horizontal dashed line denotes the +2.7 kcal/mol threshold. (c) *Left*, representative western blot of individual samples across I3-01 designs. *Right*, secreted yield quantification of the same series measured in triplicate. (d) Characterization of the Degreased I3-01NS purified from mammalian cell supernatants by DLS, SEC, and nsEM with 2D class averaging confirmed its structure is identical to that of I3-01 expressed in bacteria (Hsia et al., 2016).

We used the Degreaser to identify substitutions that could increase the dG_ins,pred_ of both the N- and C-terminal cryptic transmembrane domains of I3-01 (Supplementary Figure S5). The single substitution that had the greatest effect on both dG_ins,pred_ and secretion was H35D in the N-terminal region, though K25D also substantially increased secretion yield (Figure 3a). A triple mutation in the C-terminal region (L171Q/S177E/V180N) also increased secreted yield. Combining mutations in both regions resulted in a quadruple mutant (H35D/L171Q/S177E/V180N) that secreted at ten-fold higher levels than any of the tested single-residue variants. We again evaluated the utility of Degreaser mutations in the context of orthogonal mutations by introducing the four Degreaser-identified substitutions into a negatively ‘supercharged’ I3-01 variant, I3-01-neg. While the I3-01-neg mutations did not affect the predicted transmembrane insertion potential of I3-01 (Supplementary Figure S5), they did substantially improve overall protein expression (Supplementary Figure S11b), which may explain our observation of improved secreted yield for this variant even in the absence of Degreaser mutations (Figure 3a). Nevertheless, incorporation of the four Degreaser-identified mutations into I3-01-neg further improved its secretion, resulting in a 64-fold increase in secreted yield compared to I3-01 (Figure 3c). This variant, I3-01NS, retained its ability to assemble to the known icosahedral architecture of I3-01: SEC, DLS, and nsEM all indicated monodisperse particles that were indistinguishable from previously published I3-01 data (Figure 3d; (Hsia et al. 2016)). These results establish that elimination of cryptic transmembrane domains through sequence redesign can be used to improve the secretion of homomeric self-assembling protein nanoparticles, including the I3-01 nanoparticle that has been used as a hyperstable scaffold in several biotechnological applications (Rajoo et al. 2018; Bruun et al. 2018; Cohen et al. 2021; He et al. 2021).

### *De novo* design of secretion-optimized one-component protein assemblies

Given the success of the Degreaser in retroactively improving the secretion of several nanoparticle components and a computationally designed nanoparticle, we next tested its prospective use and compatibility with large-scale design protocols by incorporating it into the design of a set of new one-component nanoparticles intended to secrete robustly from mammalian cells. We used as building blocks a set of 1,094 models of trimeric proteins consisting of *de novo* helical bundles fused to designed helical repeat proteins as previously described (Hsia et al. 2021). These building blocks were docked as rigid bodies into three target architectures containing three-fold symmetry axes: icosahedral (I3), octahedral (O3), and tetrahedral (T3) (Figure 4a). After docking, residues at interfaces with adjacent building blocks were designed using Rosetta to enable spontaneous self-assembly to the target architecture. Three fully automated design protocols were compared: OG, ND, and DG (Figure 4a). The OG protocol used a conventional, dG_ins,pred_-agnostic protocol and therefore generated designs that had dG_ins,pred_ values both above and below +2.7 kcal/mol. The ND protocol simply applied a post-design filter to the OG design set that rejected any designs with dG_ins,pred_ less than +2.7 kcal/mol. Finally, the DG protocol incorporated the Degreaser after the interface design step and also filtered out any designs with dG_ins,pred_ less than +2.7 kcal/mol. In this benchmark design set, we used the conservative approach of allowing the Degreaser to change at maximum one residue per design, focusing on decreasing the hydrophobicity of the lowest dG_ins,pred_ segment. It is important to note that not all DG designs harbor a Degreaser mutation, as designs that do not have segments of low dG_ins,pred_ pass the Degreaser step and are accepted without modification.

**Figure 4.**
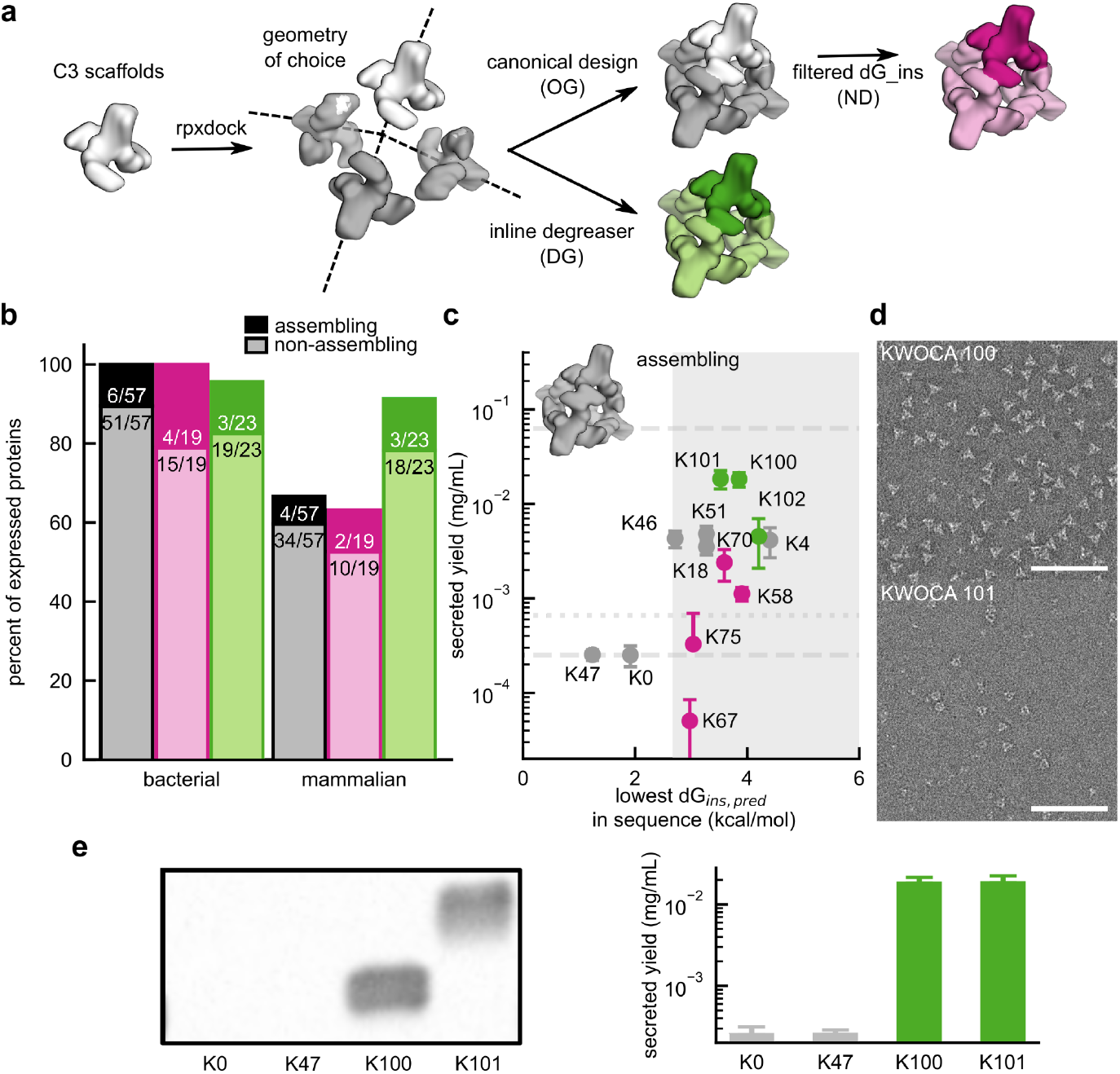
Incorporation of the Degreaser prospectively during design to generate *de novo* designed secreted protein assemblies. (a) Trimeric building blocks were docked into a desired geometry: tetrahedral, octahedral, or icosahedral. For the KWOCAs, designs were run independently for DG and OG sets, while ND designs were selected from a filtered subset of all OG designs. (b) Expression and secretion characterization of KWOCAs shows the benefit of the Degreaser on secreted yield (positive expression in mammalian cells determined as greater secretion than I3-01). Assemblies validated by nsEM are highlighted in darker color. (c) nsEM-verified assembling (Supplementary Figure S9) secreted proteins partitioned into OG, ND, and DG groups show the enhanced secreted yield of DG designs. (d) Constructs purified from mammalian material assembled into well-defined particles, indistinguishable from those expressed in bacteria (Supplementary Figure S9). Scale bar, 100 nm. (e) *Left*, representative western blot of KWOCAs with lowest dG_ins,pred_ and secretion yield (K0 and K47) and highest secretion yield (K100 and K101). *Right*, quantification of secreted yield, measured in triplicate.

Incorporation of Degreaser-guided design into an otherwise conventional design protocol did not substantially perturb the structural metrics typically used to gauge the quality of designed nanoparticle interfaces. Within the DG design set, 420 of the 1,048 designs (40%) were actually mutated by the Degreaser, while mutations meeting the Degreaser criteria were not identified for 18 designs which were therefore rejected. Notably, there was an increase in dG_ins,pred_ of the Degreaser-guided design models before any sequences were filtered on dG_ins,pred_ (47% > +2.7 kcal/mol compared to 36% for the OG design models). After filtering, DG designs that were not mutated by the Degreaser had an average dG_ins,pred_ of +3.97 kcal/mol, while those bearing mutations had an average dG_ins,pred_ of +3.38 kcal/mol. Because sequences with originally high dG_ins,pred_ are not mutated, the lower average dG_ins,pred_ of sequences with mutations is due to the low original dG_ins,pred_ of those sequences. The Degreaser mutations led to an average increase in dG_ins,pred_ of 1.41 kcal/mol but no significant differences in other design metrics (Supplementary Figure S6a). A slight shift in the distribution of ddG (the Rosetta-predicted energy of interface formation) was observed, which was expected due to Degreaser-introduced polar residues in what remained predominantly hydrophobic interfaces, accompanied by a small shift in the distribution of the interface shape complementarity (Sc; (Lawrence and Colman 1993)). On the other hand, the solvent-accessible surface area buried at the interface (sasa) showed a nearly identical distribution to that of conventional (OG) and dG_ins,pred_-filtered (ND) designs. After filtering on several structural metrics and visual inspection of the top-scoring designs by ddG, we selected 99 KWOCAs (Khmelinskaia-Wang one-component assemblies) for experimental characterization. These included 57 OG, 19 ND, and 23 DG designs, which differed in dG_ins,pred_ but not other structural metrics (Supplementary Figure S6b). 8 of the selected DG designs included mutations introduced by the Degreaser.

We first expressed the KWOCAs in the cytoplasm of *E. coli* to determine which ones successfully assembled to the intended structures regardless of secretion from mammalian cells. All KWOCAs but one yielded sufficient protein in the soluble fraction of clarified *E. coli* lysates for purification and characterization (Figure 4b). During SEC purification, 22 of 99 KWOCAs yielded a peak with an elution volume corresponding to a protein complex larger than that expected for a trimer, but smaller than unbounded aggregates (Supplementary Figure S7a). Most of these also showed peaks at elution volumes corresponding to unassembled trimeric protein. DLS of fractions from the early peaks indicated that the majority of these 22 designs formed monodisperse assemblies (Supplementary Figure S7b), and nsEM confirmed that 13 assembled to homogeneous nanoparticle structures (Supplementary Figure S8). The proportion of designs confirmed to adopt homogeneous structures was identical between conventionally designed proteins (OG+ND) and Degreaser-designed candidates (10/76 (13%) vs. 3/23 (13%), respectively; Fig. 4b). Crystallization of several KWOCAs resulted in high-resolution structures of trimeric building blocks of one non-assembling KWOCA (KWOCA 39; Supplementary Figure S9) and four assembling KWOCAS, including one confirmed by nsEM (KWOCAs 60, 65, 73, and 102) (Supplementary Figure S10a). The crystal structures matched closely the monomeric subunit of the design models (1.0–2.2 Å Cɑ rmsd), with larger deviations observed at the level of the trimers (2.2–4.0 Å Cɑ rmsd). The differences suggest that multiple aspects of the designed proteins, such as the helical bundle interface, flexibility within the designed helical repeat domains, and flexibility at the junction of the helical repeat domains and trimeric helical bundles, contribute subtle structural deviations that propagate across the trimer, which may prevent formation of the target architecture in the crystal.

We next evaluated secretion of the KWOCAs from transfected Expi293F cells by measuring the levels of myc-tagged protein in clarified harvest fluid by western blot (see Methods). A majority of the KWOCAs (72%) secreted with greater yield than I3-01, our benchmark modestly secreted protein nanoparticle (Figure 4b,c). The higher success in secretion within the DG KWOCA set (87% compared to 67% and 63 in the OG and ND sets, respectively) confirm the utility of the Degreaser in predicting and improving protein secretion. For further analysis, we separated the experimentally characterized KWOCAs into two categories: non-assembling proteins (Supplementary Figures S9 and S11) and confirmed assemblies (Supplementary Figures S8, S10 and S11). The non-assembling proteins show only a weak trend of higher secretion yield with higher dG_ins,pred_ (Supplementary Figure 11a), though this is confounded by the varying overall expression levels of these proteins (Supplementary Figure S11b). Although we did not observe consistent differences in secretion yield among the OG, ND, and DG sets of non-assembling proteins, inline application of the Degreaser (DG) tended to provide a greater benefit to secreted yield than filtering on dG_ins,pred_ after conventional design (ND) (Supplementary Figure S11a). Of the 13 EM-validated assembling designs (Supplementary Figure S8), 9 secrete at higher levels than the original I3-01 design (Figure 4c), and the highest secreted yield (KWOCA 101) was within two-fold of the highest-secreting redesigned variant of I3-01 (I3-01NS; Figure 3c). Characterization of eight of these KWOCAs by SEC, DLS, and nsEM revealed that each was indistinguishable from its bacterially produced counterpart (Figure 4d and Supplementary Figures S7 and S8). Only two of the verified assemblies (KWOCAs 0 and 47, from the OG design set) have segments of low dG_ins,pred_ (<+2.7 kcal/mol; Figure 4c). Those two assemblies were secreted at very low levels, while inline application of the Degreaser led to the highest yield of secreted nanoparticles (KWOCAs 100 and 101; Figure 4e). Together, these data indicate that the Degreaser can be applied to improve secreted yield from mammalian cells while maintaining a similar success rate in design outcome.

Comparison of several pairs of closely related designs yielded additional insights into secretion determinants. For example, KWOCA 51 and 101, which form closely related tetrahedral assemblies, used the same input scaffold for design and differ by only two residues. However, KWOCA 101 has a higher lowest dG_ins,pred_ and secreted with a roughly four-fold greater yield than KWOCA 51 (Figure 4c), highlighting how small changes in protein sequence can lead to considerable changes in secreted yield. Also related are the Degreased KWOCA 100 and the conventionally-designed KWOCA 46, both confirmed assemblies (Supplementary Figure S9), in which a one-residue difference led to a +1.13 kcal/mol change in dG_ins,pred_ and a five-fold increase in secretion. In both of these cases, the conventional design pipeline resulted in assemblies that secrete poorly and would require retrospective application of the Degreaser. Two other pairs of designs suggested that assembly state may affect secreted yield. The non-assembling KWOCA 88 differs from the octahedral KWOCA 4 by only two residues, but KWOCA 88 secretes with a roughly four-fold higher yield (Figure 4c and Supplementary Figure S11a). Finally, even though KWOCA 73 differs from KWOCA 41 by only two residues, the former showed higher-order material by SEC and DLS whereas the latter did not (Supplementary Figure S8), and KWOCA 73 secretes at about half the yield of KWOCA 41 even though its lowest dG_ins,pred_ value is much higher. Thus, although there appears to be a general secretion penalty for self-assembling proteins, these data further support that in-line use of the Degreaser during design can improve secreted nanoparticle yield.

We obtained single-particle cryoEM reconstructions of two highly secreted assemblies, KWOCA 51 and KWOCA 4, to evaluate our design protocol at high resolution. DLS and SEC indicated that both designs assemble into monodisperse nanoparticles, with KWOCA 51 forming a smaller particle than KWOCA 4 (∼19 and 26 nm hydrodynamic diameter, respectively) as expected by design (∼17 and 32 nm, respectively) (Figure 5a). Comparing calculated to experimental SAXS profiles further revealed that KWOCA 51 homogeneously assembles into the intended tetrahedral geometry, while KWOCA 4 significantly deviates from the design model (Figure 5b). Indeed a single-particle cryo-EM reconstruction of KWOCA 51 at 5.1 Å resolution closely matched the design model, and relaxing the model into the density led to only minor deviations within each subunit (1.3 Å Cɑ rmsd) that mainly reflect slight structural flexibility of the helical repeat domain (Figure 5c and Supplementary Figure S10b). In contrast, a cryo-EM map of KWOCA 4 at 6.6 Å resolution revealed that the protein does not form the computationally designed icosahedral assembly, instead identifying an octahedral nanoparticle as the only species present in the assembly fraction from SEC (Figure 5d and Supplementary Figure S10b,c). Accordingly, a SAXS profile calculated from a cryo-EM model obtained by fitting and relaxing trimeric building blocks into the density closely matched the experimental data (Supplementary Figure S10d). Interestingly, only minor structural deviations within the trimeric building blocks were observed when comparing the computational design model to the relaxed cryo-EM model (2.2 Å Cɑ rmsd), indicating that the off-target assembly must be due to differences in the computationally designed interface between the trimers. Indeed, the angle between two contiguous subunits in the cryo-EM model is rotated by 27°, resulting in a deviation of 18 Å of the contiguous subunit compared to the design model (Figure 5e and Supplementary Figure S10c). This rotation is further accompanied by a 3 Å transverse translation of the center of mass of the designed interface past the C2 symmetry axis, suggesting that the residues on the periphery of the originally designed interface were loosely packed and only weakly contributing to the interface energy. To our knowledge, this is the first report of a high-resolution structure of a *de novo* computationally designed protein nanoparticle that forms a well-defined architecture distinct from the one intended. Nevertheless, the two structures together establish that KWOCAs 4 and 51, which secrete from mammalian culture at higher levels than lumazine synthase and I3-01, are highly ordered, monodisperse *de novo* designed nanoparticles.

**Figure 5.**
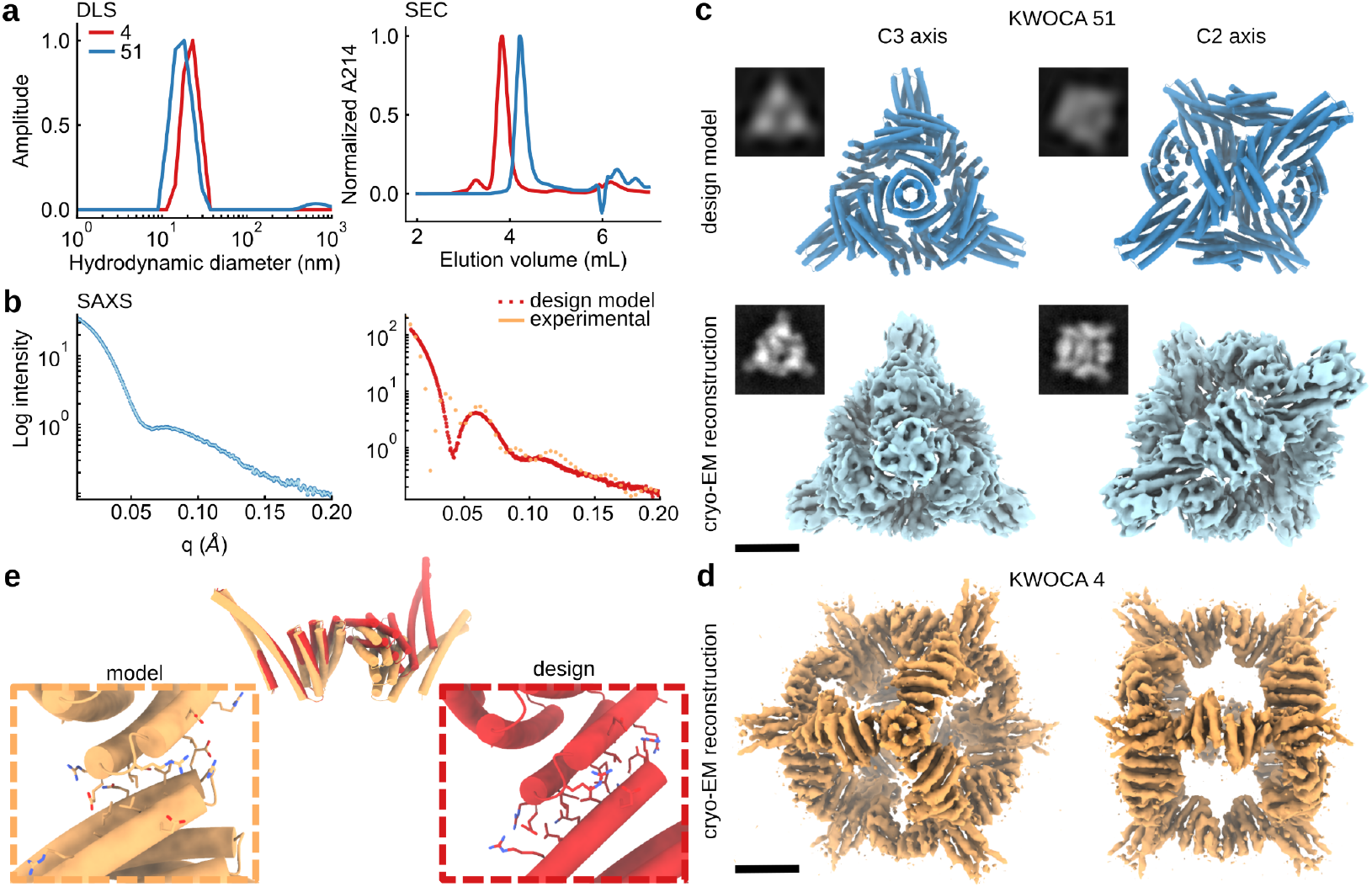
Structural characterization of KWOCA 4 and KWOCA 51. (a) DLS, SEC traces and (b) SAXS profiles of KWOCA 51 (blue) and KWOCA 4 (red/orange). (c) Design model and cryo-EM density map of KWOCA 51. (d) Cryo-EM density map of KWOCA 4. (e) Overlay of two KWOCA 4 subunits across the designed nanoparticle interface, highlighting the interface contact angle difference between the design model and the best-fitting cryo-EM model. Theoretical SAXS profiles calculated from the design models (dotted darker lines) are overlaid with the experimentally obtained SAXS profiles (b). The fits to the theoretical SAXS profiles calculated to the best-fitting cryo-EM models can be found in Supplementary Figure S10d. Scale bar, 5 nm (c,d).

## Discussion

Computational protein design methodologies are advancing rapidly, enabling access to previously unexplored spaces in protein structure and function (Huang, Boyken, and Baker 2016; Baek and Baker 2022). In addition to increasing our fundamental understanding of proteins, these advances have made commercial application of computationally designed proteins a reality. For example, computationally designed cytokine mimetics (Silva et al. 2019), enzymes for gluten degradation (Gordon et al. 2012), and nanoparticle vaccines (Marcandalli et al. 2019; Boyoglu-Barnum et al. 2021; J. G. Jardine et al. 2015) have recently advanced to clinical trials, and a designed nanoparticle vaccine for COVID-19 (Walls et al. 2020) was recently approved for use in South Korea. As designed proteins become increasingly useful, methods for optimizing various phenotypes other than structure and stability become more important. Recent examples include the ability to design heterodimers while maintaining the solubility of the individual components (Sahtoe et al. 2022) and methods for modulating the brightness and chromophore specificity of de novo mini fluorescent proteins (Dou et al. 2018; Klima et al. 2021). To enable their modular implementation in a wide variety of design applications, design methods like these must be able to optimize such phenotypes without sacrificing structural stability. The Degreaser was explicitly constructed to be modular—as showcased by our redesign of existing proteins as well as our application of the Degreaser in-line during the design of new secretable protein assemblies—while preserving structural stability and integrity. These features in principle enable its application to any protein. Furthermore, application of the Degreaser in-line during design is minimally invasive: it only mutates proteins that require elimination of cryptic transmembrane domains, and it identifies the minimal sufficient perturbation. As we showed during KWOCA design, this approach allows in-line implementation of the Degreaser that should eliminate the requirement for retroactive re-design of poorly secreting proteins.

More broadly, any method for improving the yield of recombinant biologics is valuable. For example, the decades of effort invested in optimizing and industrializing the production of monoclonal antibodies now underpins the biologics industry (Kelley 2009). Methods like the Degreaser that encode improved yield or performance in the sequence of the molecule itself are especially desirable, as they make the improvements “automatic”: they do not require other actions like the use of specialized cell culture media or co-transfection of chaperones. Numerous examples of enzyme and antigen re-design to improve yield highlight the utility of this approach (Dalvie, Brady, et al. 2021; Dalvie, Rodriguez-Aponte, et al. 2021; Ellis et al. 2021; Hsieh et al. 2020; Rawi et al. 2020; Pallesen et al. 2017; Malladi et al. 2020; Peleg et al. 2021; Campeotto et al. 2017). Methods that encode improvements genetically are of increasing importance now that genetic delivery of biologics has become clinical reality with the recent licensure of several AAV-based gene therapies and mRNA vaccines (D. Wang, Tai, and Gao 2019; Chaudhary, Weissman, and Whitehead 2021). Furthermore, secreted protein nanoparticle immunogens are beginning to be explored as a strategy for improving the potency of mRNA vaccines (Konrath et al. 2022; Z. Xu et al. 2020; Mu et al. 2021). The Degreaser and the new highly secretable KWOCAs we describe here could pave the way towards the design of mRNA-launched nanoparticle vaccines with atomic-level accuracy. This approach will enable structural and functional optimization of the nanoparticle scaffolds in ways that are not possible when relying on naturally occurring scaffolds. For example, we previously showed that the *de novo* design of novel nanoparticle scaffolds enabled precise optimization of termini placement for genetic fusion of functional domains (Ueda et al. 2020b).

Several features in our data suggest potential future directions for the Degreaser. First, to test the suitability of the Degreaser as part of a “set it and forget it” large-scale design protocol, we used the conservative strategy of only allowing one mutation maximum in the DG KWOCA designs. However, the results we obtained retroactively degreasing I3-01 and the individual scaffolds of two-component nanoparticles indicate that more aggressive use of the Degreaser (i.e., enabling multiple mutations) could lead to larger improvements in the secretion of specific proteins, and could lead to a higher success rate in designing highly secretable proteins while still preserving the intended structure and function. Second, although we have shown the presence of cryptic transmembrane domains significantly influences protein secretion, there are other factors that affect secretion and should be considered when designing secreted proteins (Dalvie, Brady, et al. 2021; Dalvie, Rodriguez-Aponte, et al. 2021). For example, a recent high-throughput study of the secretion of protein fragments in yeast identified several sequence-based features that could be incorporated in future secretion models (Boone et al. 2021). Finally, here we applied the Degreaser to homomeric nanoparticles and the oligomeric components of two-component nanoparticles due to the many potential applications of highly secretable designed protein nanoparticles. Future application of the Degreaser to two-component nanoparticles, other designed proteins, and naturally occurring proteins that have not evolved to secrete at high levels could be used to improve the secretion of many different potential biologics and enable new applications of designed proteins.

## Methods

### The Degreaser

The Degreaser is a Mover in the Rosetta software suite that takes as input a protein structure, represented within Rosetta as a Pose, and evaluates its sequence for segments where dG_ins,pred_ is less than a user-specified threshold. Calculations and equations used to calculate dG_ins,pred_ of protein sequences are described in (Hessa et al. 2007). In brief, each 19-residue segment (though non-19-residue windows can be used with a length correction term) within a sequence is considered as approximately helical, and the position-specific amino acid contribution to dG_ins,pred_ is calculated for each position within that segment, then summed to give the dG_ins,pred_ of the segment at that sequence position. We set the detection threshold of the DegreaserMover to +3.5 kcal/mol in order to capture any potential regions of hydrophobicity. This value was chosen to include even marginally hydrophobic segments.

Within each detected segment, one residue at a time (the ‘focus residue’) is allowed to mutate to a user-defined set of amino acid identities, which are by default polar residues (DEKRQNSTY). Histidine was excluded from this set as its unprotonated form is potentially not polar enough to perturb the dG_ins,pred_ of a given structure. The Rosetta PackRotamersMover, or Packer (described in detail in (Leaver-Fay et al. 2011)), alters the identity of the focus residue using a Monte Carlo search method, and simultaneously samples rotamers of the surrounding residues without altering their identities (repacking). The resultant candidate Pose is accepted as the most energetically favorable perturbation of the focus residue.

The perturbed Pose is then evaluated for its effect on both dG_ins,pred_ as well as on Rosetta score. We set an upper threshold for the change in Rosetta score of +15 REU, which corresponds to less than 1% of the total score of most of the previously designed protein assemblies that were part of our first testing set. We note that setting this threshold higher led to the incorporation of many residues that yielded subtle clashes within the design models. However, this score threshold can be raised in order to more aggressively perturb existing structures, especially in cases where multiple mutations may be necessary. For perturbed Poses where the first Packer-generated residue exceeds the allowed increase in Rosetta score, the focus position is reverted to wild-type and not considered further. For Pose candidates in which the first Packer-generated residue meets the scoring criteria but dG_ins,pred_ is not increased (above a user-given threshold amount, default dG_ins,pred_ > 0.27 kcal/mol), the identity of that candidate is disallowed at that focus residue and the process is repeated. In this manner, the Degreaser semi-exhaustively samples possible mutations at each position within a hydrophobic segment that minimally perturbs the structure while still satisfying the desired increases in dG_ins,pred_. When a Pose is accepted, its properties are saved and logged. Importantly, after all positions in all segments are sampled, the Degreaser ‘chooses’ the Pose that gives the largest increase in dG_ins,pred_ at the lowest initially-identified dG_ins,pred_ in the sequence. This logic is fundamental as, in its absence, the dG_ins,pred_ of one segment within a protein can be significantly increased but the resulting lowest dG_ins,pred_ of the sequence does not change. By maximally perturbing the lowest dG_ins,pred_ segment within a Pose, we guarantee the largest and most impactful dG_ins,pred_ change.

In the examples of Degreaser-guided protein nanoparticle (re)design reported here, we allowed only one mutation per input structure. Although only interfacial residues were allowed to design within the conventional design protocol, the Degreaser is allowed to change any of the residues it identifies to be within hydrophobic segments. However, the Degreaser can be specified to only operate on a subset of residues within a given model, much as any other Mover can be. By allowing only one mutation, our goal was to minimally disturb the interfaces resulting from conventional design, which contain between 7 and 24 residues that participate in the hydrophobic interface and may not be able to easily accommodate several mutations. However, the Degreaser is amenable to allowing an arbitrary number of mutations per hydrophobic segment. Furthermore, not every hydrophobic segment identified is in the vicinity of the designed interface. Both considerations warrant further investigation.

Further relaxation of individual Degreaser variants yielded varying results, as the incorporation of polar residues into regions near designed nonpolar interfaces or other segments of the original model led to differing consequences. For example, some design models were already relaxed, and further relaxation of its Degreaser variant caused unexpected backbone movements due to the introduction of small clashes. Combinations of Degreaser-identified variants for previously-designed protein nanoparticles were generated manually. These combinations were directly additive with respect to dG_ins,pred_, while the effects on Rosetta score varied depending on the distance between the variant positions in 3D space. We found that combinations of variants in different segments within one protein can be considered almost entirely independently, while combinations of variants within a given segment were considered more cautiously to make sure that the original structure was not too significantly perturbed.

### Computational design of protein nanoparticles

Trimeric scaffolds were generated by helical fusion of previously designed trimeric helical bundles (Boyken et al. 2019) and *de novo* helical repeat domains (Brunette et al. 2015), following the protocol described in (Hsia et al. 2021). Symmetrical docking of the top scoring 1094 trimers was performed using the rpxdock protocol (https://github.com/willsheffler/rpxdock). Briefly, the three-fold symmetry axis of each trimeric scaffold was aligned with corresponding three-folds in one of the target symmetries: T, O, or I. These aligned trimers may rotate around and translate along their respective symmetry axis while maintaining the symmetry of the complex (King et al. 2012, 2014). These two degrees of freedom, radial displacement and axial rotation, were sampled in increments of 1 Å and 1°, respectively. For each docked configuration in which no clashes between the backbone and beta carbon atoms of adjacent building blocks were present, an rpx (rotamer pair transform) designability score was calculated (Fallas et al. 2017). High-scoring docked configurations with intermediately sized interfaces (ncontacts < 75) were selected for full-atom interface design using Rosetta scripts as previously described (King et al. 2014; Hsia et al. 2016). Briefly, the design protocol took a single-chain input pdb and a symmetry definition file containing information for a target cubic point group symmetry (DiMaio et al. 2011). The oligomers were then aligned to the corresponding axes of the target symmetry using the Rosetta SymDofMover, taking into account the rigid body translations and rotations retrieved from the .pickle file output from the docking protocol (King et al. 2014; Hsia et al. 2016). The conventional symmetric interface design protocol was modified for in-line Degreaser design by adding the DegreaserMover step after the final step of conventional design, before any filters were applied to a particular design (see Supplementary Information). Individual design trajectories were filtered by the following criteria: difference between Rosetta energy of bound and unbound states (ddG) less than −20.0 REU, interface surface area greater than 500 Å^2^, shape complementarity (sc) greater than 0.6, and at least three alpha helices at the interface. Designs arising from the conventional design protocol were further filtered to ensure that dG_ins,pred_ > 2.7 to generate the ND pool, and DG designs were also filtered to ensure that dG_ins,pred_ > 2.7. Designs that passed these criteria were manually inspected and a set of 99 designs selected for experimental characterization: 57 OG, 19 ND, and 23 DG.

### Cell culture, protein expression and purification

All bacterial protein expression was performed with Lemo21(DE3) *E. coli* (NEB), all bacterial plasmid propagation with NEB5ɑ *E. coli* (NEB), and all mammalian protein expression with Expi293F cells (ThermoFisher Scientific). All bacterial expression was performed from a pET29b(+) vector with open reading frames inserted between the NdeI and XhoI restriction sites. All mammalian expression was performed from a pCMV/R vector (Barouch et al. 2005) with open reading frames inserted between the XbaI and AvrII restriction sites, and all constructs used the same IgGκ secretion signal. 6×His tags for purification, myc tags for detection, as well as GS linkers and a tryptophan residue to enable protein quantitation by UV/vis spectroscopy were placed at the N or C termini of constructs depending on available 3D space after manual inspection of design models (complete lists of gene and protein sequences can be found in Supplementary Tables S1-S2).

For bacterial expression and purification of previously described nanoparticle component proteins, see previous methods (Bale et al. 2016; Hsia et al. 2016; Ueda et al. 2020b). For bacterial expression and purification of KWOCAs, proteins were expressed by autoinduction using TBII media (Mpbio) supplemented with 50×5052, 20 mM MgSO_4_, and trace metal mix, under antibiotic selection at 18°C for 24 h after initial growth for 6-8 h at 37°C. Cells were harvested by centrifugation at 4000 *g* and lysed by sonication or microfluidization after resuspension in lysis buffer (50 mM Tris pH 8.0, 250 mM NaCl, 20 mM imidazole, 5% glycerol), followed by addition of bovine pancreatic DNaseI (Sigma-Aldrich) and protease inhibitors (Thermo Scientific). Clarified lysate supernatants were batch bound with equilibrated Ni-NTA resin (QIAGEN). Washes were performed with 5-10 column volumes of lysis buffer, then eluted with 3 column volumes of the same buffer containing 500 mM imidazole. Concentrated or unconcentrated elution fractions were further purified using a Superose 6 Increase 10/300 GL (Cytiva) on an AKTA Pure (Cytiva) into 25 mM Tris pH 8.0, 150 mM NaCl, 5% glycerol. Instrument control and elution profiles analysis were performed with Cytiva software (Cytiva).

For purification of plasmid DNA for Expi293F transfection, bacteria were cultured and plasmids were harvested according to the QIAGEN Plasmid Plus Maxi Kit protocol (QIAGEN). For mammalian expression and purification of KWOCAs and other secreted proteins, Expi293F cells were passaged according to manufacturer protocols (ThermoFisher Scientific). Cells at 3.0×10^6^ cells/mL were transfected with 1 μg of purified plasmid DNA mixed with 3 μg/μg PEI-MAX per mL cell culture. For secretion yield measurements, cells were harvested at 72 h post-transfection by centrifugation for 5 minutes at 1,500 *g*. For protein purification, cells were harvested at 120 h post-transfection by centrifugation of cells and subsequent sterile filtering of supernatant. Filtered supernatant was adjusted to 50 mM Tris pH 8.0 and 500 mM NaCl, then bound to Ni Sepharose Excel (Cytiva) with agitation overnight. Pelleted resin was washed with 50 mM Tris pH 8.0, 500 mM NaCl, 30 mM imidazole, then eluted with the same buffer containing 300 mM imidazole. Concentrated elution fractions were purified by size-exclusion chromatography as described above.

Protein content and purity at each step of expression and purification were analyzed by SDS-PAGE using Criterion precast gels and electrophoresis systems (BIO-RAD). Purified protein concentration measurements were measured using UV absorbance at 280 nm, and calculated using theoretical molar extinction coefficients (ExPasy). Proteins were concentrated with 30,000 MWCO concentrators (Millipore). Purified, concentrated, and buffer-exchanged proteins were snap-frozen in liquid nitrogen and stored at −80°C only if aggregates were absent as determined by DLS.

### Secretion yield quantification

Cells were centrifuged to separate medium from cells, and pelleted cells were resuspended in the same volume of removed medium in phosphate-buffered saline (PBS). All samples were then treated for 10 min at 37°C with 0.05% Triton-X 100 (Sigma) containing a 1:400 dilution of Benzonase endonuclease (EMD Millipore) to permeate membranes and prevent nucleic acid aggregation, facilitating quantitative gel loading. Internal myc tag protein standard was also added to a final concentration of 0.06 mg/mL. Treated samples were then diluted into 4× SDS loading buffer (200 mM Tris pH 6.8, 40% glycerol, 8% SDS, bromophenol blue, 4 mM DTT) and incubated at 95°C for 5 min. 14.3 μL of boiled samples were loaded onto Criterion 4-20% precast polyacrylamide gels (BIO-RAD). Precision Plus WesternC standards were included in each gel (BIO-RAD). Gels were run using BIO-RAD Criterion gel boxes and power supplies, then transferred using the Trans-Blot Turbo system onto 0.2 μm nitrocellulose membranes according to manufacturer instructions (BIO-RAD). Transferred blots were blocked in 3% milk in wash buffer (10 mM Tris pH 8.0, 150 mM NaCl, 0.1% Tween-20) for 30 min, then incubated with a 1:20,000 dilution of mouse anti-myc tag antibody (9B11, Cell Signaling Technology) with agitation, either 75 min at room temperature or 16 h at 4°C. Blots were then washed three times with wash buffer and incubated 75 min at room temperature with a 1:10,000 dilution of goat anti-mouse HRP conjugated antibody (Cell Signaling Technology). After three washes with wash buffer, blots were developed with Clarity ECL substrates according to manufacturer directions on a Gel Doc XR+ Imager with Image Lab software (BIO-RAD).

Western blot images were analyzed using ImageJ/FIJI software for quantification. Calibration curves of known myc-tagged protein were used to establish a linear range (data not shown), and four points for each blot were included to allow absolute concentration determination. Three biological replicates (i.e., independent transfections) were included for each construct. For some constructs, the measured cellular level of protein was higher than the linear range of the calibration curve. However, for nearly all measurements, the secretion yield measurement was within linear range.

### Protein characterization

Dynamic light scattering measurements (DLS) were performed using the default Sizing and Polydispersity method on the UNcle (Unchained Labs). 8.8 μL of SEC-purified elution fractions were pipetted into the provided glass cuvettes. DLS measurements were run with ten replicates at 25°C with an incubation time of 1 s; results were averaged across runs and plotted using Python. Other DLS measurements were also obtained using a DynaPro NanoStar (Wyatt) DLS setup with ten acquisitions per measurement, and three measurements per protein sample.

For negative stain electron microscopy, samples were diluted to 0.1-0.02 mg/mL and 3 µL was negatively stained using Gilder Grids overlaid with a thin layer of carbon and 2% uranyl formate as previously described (Veesler et al. 2014). Data were collected on an Talos L120C 120 kV electron microscope equipped with a CETA camera.

To identify the molecular mass of each protein, intact mass spectra were obtained via reverse-phase LC/MS on an Agilent 6230B TOF on an AdvanceBio RP-Desalting column, and subsequently deconvoluted by way of Bioconfirm using a total entropy algorithm. For LC, buffers were water with 0.1% formic acid and acetonitrile with 0.1% formic acid; the proteins were eluted using a gradient of 10% to 100% acetonitrile buffer over 2 min. Except for KWOCA 39 (Supplementary Figure S9), for which the purification profile according to the method described above is shown, SEC profiles for the KWOCAS (Figures 2,3,5 and Supplementary Figure S7) were obtained by high pressure liquid chromatography on an Agilent Bio SEC-5 column (Agilent) at a flow rate of 0.35 mL/min by injection of 10 μL of purified eluate using a mobile phase of Tris-buffered saline (50 mM Tris pH 8, 150 mM NaCl, 5% v/v glycerol).

### Yeast surface display

*Saccharomyces cerevisiae* EBY100 strain cultures were grown in C-Trp-Ura medium supplemented with 2% (w/v) glucose (CTUG). To induce expression, yeast cells initially grown in CTUG were transferred to SGCAA medium supplemented with 0.2% (w/v) glucose and induced at 30 °C for 16–24 h. Cells were washed with PBSF (PBS with 1% (w/v) BSA) and labeled for 30 minutes at room temperature with FITC-conjugated anti-Myc at 10 μg/ml (Immunology Consultants Lab, CYMC-45F). After incubation, cells were washed once more and resuspended in PBSF for cell sorting (Attune NxT Flow Cytometer, Thermo Fisher Scientific).

### Small-angle X-ray scattering

Selected SEC fractions were concentrated to 1-5 mg/mL in buffer containing 2% glycerol. The flowthrough was used as a blank for buffer subtraction during SAXS analysis. Samples were then centrifuged (13,000 *g*) and passed through a 0.22 µm syringe filter (Millipore). These proteins and buffer blanks were shipped to the SIBYLS High-Throughput SAXS ALS Advanced Light Source in Berkeley, California to obtain scattering data (Putnam et al. 2007; Hura et al. 2009; Classen et al. 2013; Dyer et al. 2014). Scattering traces were analyzed and fit to theoretical models using the FOXS 15 server (https://modbase.compbio.ucsf.edu/foxs/) (Schneidman-Duhovny et al. 2013, 2016).

### X-ray crystallography

All crystallization trials were carried out at 20°C in 96-well format using the sitting-drop method. Crystal trays were set up using Mosquito LCP by SPT Labtech and monitored by JANSi UVEX imaging system. Drop volumes ranged from 200 to 400 nL and contained protein to crystallization solution in ratios of 1:1, 2:1, and 1:2. Diffraction quality crystals appeared in 0.2 M NH_4_CH_3_CO_2_, 0.1 M Na_3_C_6_H_5_O_7_ pH 5.6, 30% (v/v) MPD for KWOCA 39. Diffraction quality crystals appeared in 0.1 M Tris pH 7.0, 20% (w/v) PEG-2000 MME for KWOCA 73. Diffraction quality crystals appeared in 0.2 M MgCl_2_, 0.1 M Tris pH 7.0, 10% (w/v) PEG-8000 for KWOCA 60. Diffraction quality crystals appeared in 0.2 M MgCl_2_, 0.1 M Imidazole pH 8.0, 35% (v/v) MPD for KWOCA 65. Diffraction quality crystals appeared in 0.1 M NaCH_3_CO_2_ pH 4.5, 35% (v/v) MPD for KWOCA 102. Crystals were subsequently harvested in cryo-loops and flash frozen directly in liquid nitrogen for synchrotron data collection.

Data collection from crystals of KWOCA 39, 60, and 65 were performed with synchrotron radiation at the Advanced Photon Source (APS) on beamline 24ID-C. Data collection from crystals of KWOCA 73 and 102 were performed with synchrotron radiation at the Advanced Light Source (ALS) on beamline 8.2.1/8.2.2.

X-ray intensities and data reduction were evaluated and integrated using XDS (Kabsch 2010) and merged/scaled using Pointless/Aimless in the CCP4 program suite (Winn et al. 2011). Structure determination and refinement starting phases were obtained by molecular replacement using Phaser (McCoy et al. 2007) using the design model for the structures. Following molecular replacement, the models were improved using phenix.autobuild (Adams et al. 2010); efforts were made to reduce model bias by setting rebuild-in-place to false, and using simulated annealing and prime-and-switch phasing. Structures were refined in Phenix (Adams et al. 2010). Model building was performed using COOT (Emsley and Cowtan 2004). The final model was evaluated using MolProbity (Williams et al. 2018). Data collection and refinement statistics are recorded in Supplementary Table S3. Data deposition, atomic coordinates, and structure factors reported in this paper will be deposited in the Protein Data Bank (PDB), http://www.rcsb.org/, prior to publication.

### Cryo-electron microscopy

Cryo-EM grids were prepared using Vitrobot mark IV (Thermo Fisher Scientific). Vitrobot settings were as follows: KWOCA 4 - *T* = 10°C, Humidity = 100%, Blotting force = 0, Wait time = 10 s, Blotting time = 5 s; KWOCA 51 - *T* = 22°C, Humidity = 100%, Blotting force = 1, Wait time = 10 s, Blotting time = 7 s.

KWOCA 4 sample was at a concentration of 3.58 mg/mL in 25 mM Tris pH 8.0, 150 mM NaCl, 5% glycerol. 0.5 µL of 0.04 mM lauryl maltose neopentyl glycol (LMNG) stock solution was mixed with 3.5 µL of the KWOCA 4 sample and 3 µl were immediately loaded onto the UltrAuFoil R 1.2/1.3 grids (Au, 300-mesh; Quantifoil Micro Tools GmbH). Prior to sample loading the grids were subjected to Ar/O_2_ plasma cleaning for 10 s on a Solarus 950 plasma cleaner (Gatan). After the blotting step, the grids were plunge-frozen into liquid-nitrogen-cooled liquid ethane. For KWOCA 51, we applied 2 μL of 3 mg/mL protein in 25 mM Tris, 150 mM NaCl, pH 8.0, 100 mM glycine to glow-discharged C-flat CF-2/2 C-T-grids (TED PELLA).

KWOCA 4 cryo-EM samples were imaged on Talos Arctica transmission electron microscope (Thermo Fisher Scientific) operating at 200 kV. The microscope was equipped with a sample autoloader and a K2 summit direct electron detector (Gatan) operating in counting mode. Nominal exposure magnification was 36,000 with a resulting pixel size at the specimen plane of 1.15 Å. KWOCA 51 data collection was performed on an FEI Titan Krios Electron Microscope operating at 300 kV. The microscope was equipped with a Gatan Quantum GIF energy filter and a K3 Summit direct electron detector (Li et al. 2013) operating in electron-counting mode. Nominal exposure magnification was 105,000 with a resulting pixel size at the specimen plane of 0.85 Å. Automated data collection was performed using Leginon software (Carragher et al. 2000; Suloway et al. 2005). Data collection information is presented in Supplementary Table S4.

For KWOCA 4, raw frames were aligned and dose weighted using MotionCor2 (Zheng et al. 2017). GCTF (K. Zhang 2016) was applied for estimation of CTF parameters. From 1,341 collected micrographs, we extracted 126,365 clean KWOCA 4 particles. Initial data processing steps (particle picking, 2D classification, initial model generation) were performed in cryoSPARC.v3.2.0 (Punjani et al. 2017) and the particles were subsequently transferred to Relion/3.0 (Zivanov et al. 2018) for further processing. Octahedral symmetry was applied for all 3D classification and refinement steps. For KWOCA 51, all data processing was carried out in CryoSPARC. Alignment of movie frames was carried out using Patch Motion with an estimated B-factor of 500 Å^2^. Defocus and astigmatism values were estimated using Patch CTF. ∼800,000 particles were picked in a reference-free manner using Blob Picker and extracted with a box size of 440 Å. An initial round of 2D classification was performed in CryoSPARC using 100 classes and a maximum alignment resolution of 6 Å. ∼398,000 selected particles were re-centered and re-extracted with a box size of 360 Å. An *ab initio* reconstruction was generated using a C1 symmetry operator on the dose-weighted and re-centered particles. A homogeneous refinement was next performed using this *ab initio* model as a starting reference. Tetrahedral symmetry was applied during this refinement, leading to an initial estimated map resolution of 5.95 Å. Local motion within single movies was corrected using an estimated B-factor of 500 Å, and particles were re-extracted with a box size of 360 Å. A second round of homogeneous refinement was performed, resulting in an improved resolution estimate of 5.8 Å. Particles were next split into separate optics groups and re-refined to a final estimated resolution of 5.6 Å. Relevant information is presented in Supplementary Figure S10. The resulting maps will be deposited to the Electron Microscopy Data Bank (EMDB) prior to publication. Copies of the design models of each trimeric scaffold were fit into the respective cryo-EM density map using Chimera (Pettersen et al. 2004). The resulting model was relaxed, starting from a single monomer to which a symmetry operator was applied, guided by the cryo-EM density using Rosetta (DiMaio et al. 2015; R. Y.-R. Wang et al. 2016).

To calculate the root-mean-square (rms) deviation of the experimentally obtained crystal structures or relaxed cryo-EM models to the computational design models, the pair_fit function of PyMol was used on the common Cɑ carbons of the monomeric subunit of the pair of models to be compared. Additionally, the rms of the whole trimer was calculated using the rms_cur function on the common Cɑ carbons.

### Other methods

Figures created using Inkscape. Data processing and plotting were performed with LibreOffice Calc and Python. Protein structure rendering was performed in PyMOL (http://www.pymol.org/pymol) or ChimeraX (Pettersen et al. 2021).

## Supporting information

design script

## Author Contributions

Conceptualization: J.Y.W., A.Kh., N.P.K.; Methodology: J.Y.W., A.Kh.; Software: J.Y.W., A.Kh., W.S.; Formal analysis: J.Y.W., A.Kh., A.A., A.J.B., A.B.; Investigation: J.Y.W., A.Kh., A.A., A.J.B., C.S., U.N., D.E., C.W., M.C.M., M.A., S.C., A.Ka., H.N., C.S., B.S., M.W., A.B.; Resources: Y.H., D.B.; Data curation: J.Y.W., A.Kh., A.A., A.J.B., A.B.; Writing – original draft preparation: J.Y.W., A.Kh., and N.P.K.; Writing – review & editing: All authors; Visualization: J.Y.W., A.Kh., A.A., A.J.B.; Supervision: L.C., B.F., M.M., A.B.W., N.P.K.; Funding acquisition: D.B., A.B.W., N.P.K.

## Acknowledgments

The authors thank Bill Anderson, Hannah L. Turner, and Jean-Christophe Ducom (The Scripps Research Institute) for their assistance with electron microscopy experiments; Ratika Krishnamurty for program management; and members of the King laboratory for comments on the manuscript. This work was supported by the Bill & Melinda Gates Foundation (OPP1156262 to D.B. and N.P.K., OPP1115782 to A.B.W.), a generous gift from the Audacious Project at the Institute for Protein Design, the National Institutes of Health (P50 AI150464 to N.P.K.), the Defense Threat Reduction Agency (HDTRA1-18-1-0001 to D.B. and N.P.K.) and the Collaboration for AIDS Vaccine Discovery (INV-002916 to A.B.W.). A.A. is supported by amfAR Mathilde Krim Fellowship in Biomedical Research #110182-69-RKVA. U.N. was supported in part by a PHS National Research Service Award (T32GM007270) from NIGMS. SAXS experiments were conducted at the Advanced Light Source (ALS) a national user facility operated by Lawrence Berkeley National Laboratory on behalf of the Department of Energy, Office of Basic Energy Sciences, through the Integrated Diffraction Analysis Technologies (IDAT) program, supported by DOE Office of Biological and Environmental Research. Additional support comes from the National Institute of Health project ALS-ENABLE (P30 GM124169) and a High-End Instrumentation Grant S10OD018483. X-ray diffraction data were collected at the APS Northeastern Collaborative Access Team beamlines, which are funded by the National Institute of General Medical Sciences from the National Institutes of Health (P30 GM124165), and ALS beamline 8.2.2/8.2.1 at Lawrence Berkeley National Laboratory. The Berkeley Center for Structural Biology is supported in part by the NIH, National Institute of General Medical Sciences, and the Howard Hughes Medical Institute. Molecular graphics and analysis performed with UCSF Chimera and UCSF ChimeraX, developed by the Resource for Biocomputing, Visualization, and Informatics at the University of California, San Francisco, with support from NIH P41-GM103311, NIH R01-GM129325 and the Office of Cyber Infrastructure and Computational Biology, National Institute of Allergy and Infectious Diseases.

## Supplementary Figures

**Supplementary Figure S1:**
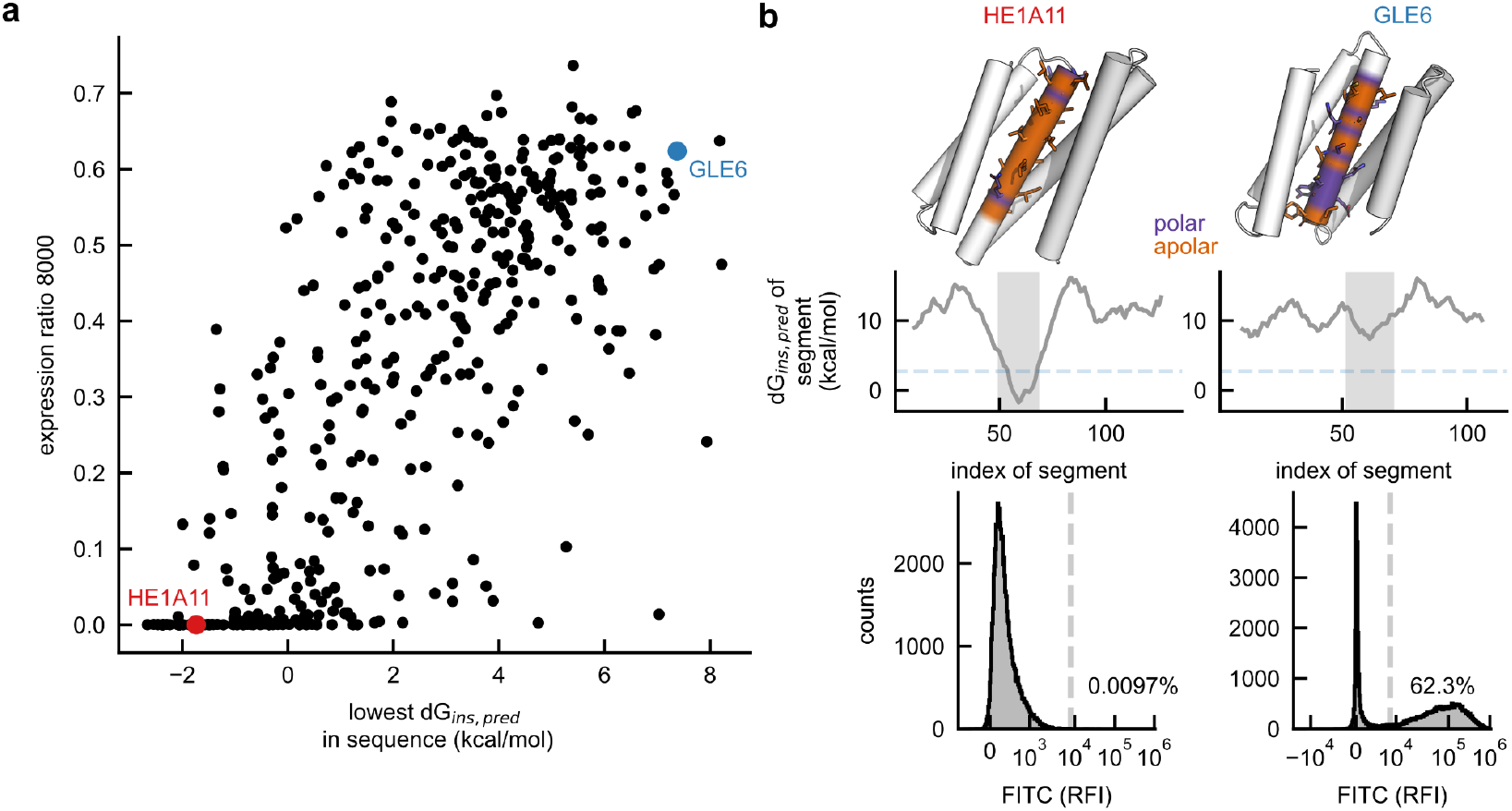
*De novo* designed proteins with low dG_ins,pred_ display poorly on the surface of yeast. (a) Of 480 *de novo* designed proteins, sequences with higher dG_ins,pred_ had stronger expression on the surface of yeast as measured by the ratio of events with FITC relative fluorescence intensity (RFI) greater than 8,000 (see vertical dashed lines in b) to total events. Because these proteins are displayed with C-terminal tags to which a fluorescently labeled antibody binds, translocation is mandatory for exterior display and subsequent fluorescent detection. (b) Two examples, one with low dG_ins,pred_ (left, HE1A11) and one with high dG_ins,pred_ (right, GLE6), highlight how proteins with similar topologies but different lowest dG_ins,pred_ lead to different surface display outcomes.

**Supplementary Figure S2:**
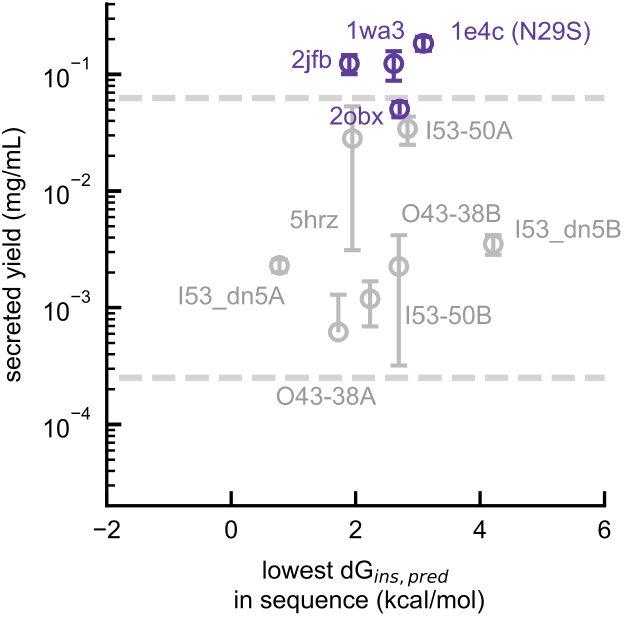
Poorly secreting designed nanoparticle proteins contain segments of low dG_ins,pred_. Material from transiently transfected cell culture was fractionated and analyzed by quantitative Western blot to assess the secreted yield of previously designed proteins and the naturally occurring scaffolds from which they were derived (I53_dn5A and O43-38B as well as their natural counterparts are repeated from Figure 2a). Horizontal dashed lines represent the upper and lower limits of the linear range of a western blot standard curve generated by densitometry.

**Supplementary Figure S3:**
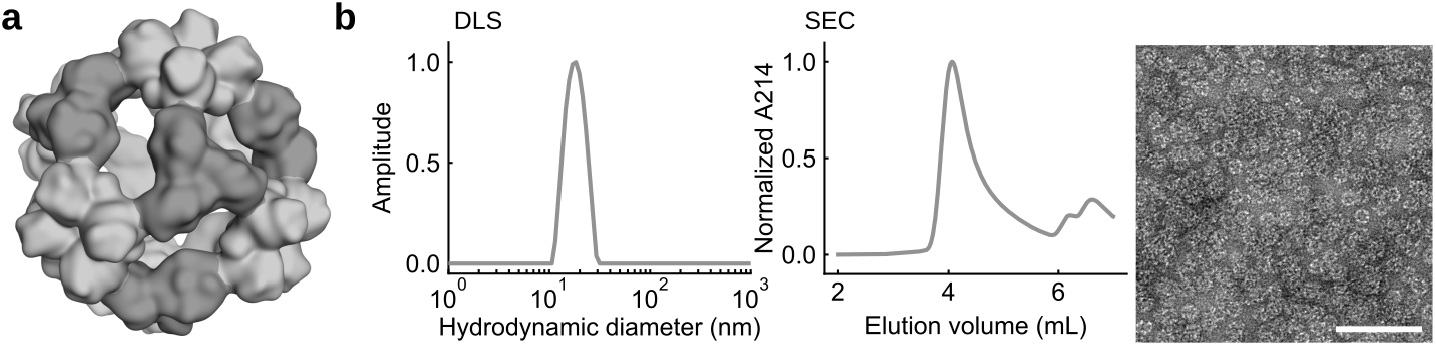
Two-component octahedral nanoparticle O43-38. (a) Design model and (b) DLS, SEC, and nsEM of *in vitro*-assembled O43-38 subunits expressed in *E. coli*. Scale bar, 100 nm.

**Supplementary Figure S4:**
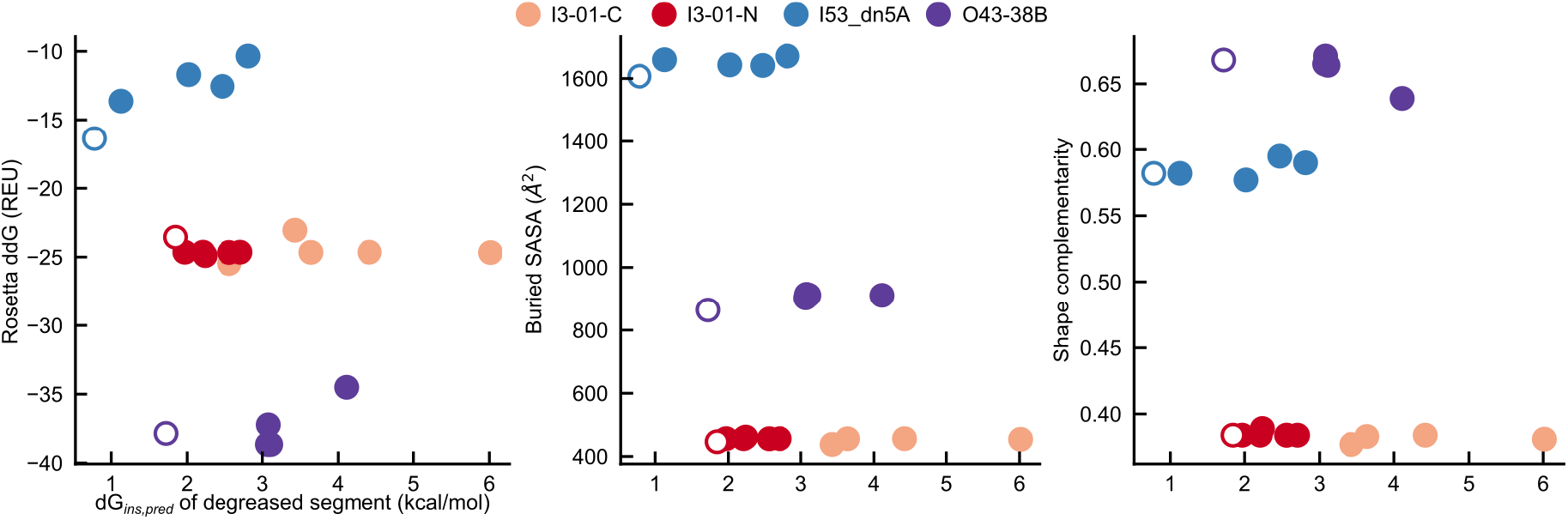
Degreaser-identified mutations do not substantially affect Rosetta metrics. Design metrics typically used to filter nanoparticle designs are shown for the I3-01, I53_dn5A, and O43-38B proteins discussed in Figures 2 and 3. I3-01 redesigns in which the N-terminal and C-terminal cryptic transmembrane domains were modified by the Degreaser are indicated separately. The original designs, before application of the Degreaser, are represented by hollow markers.

**Supplementary Figure S5:**
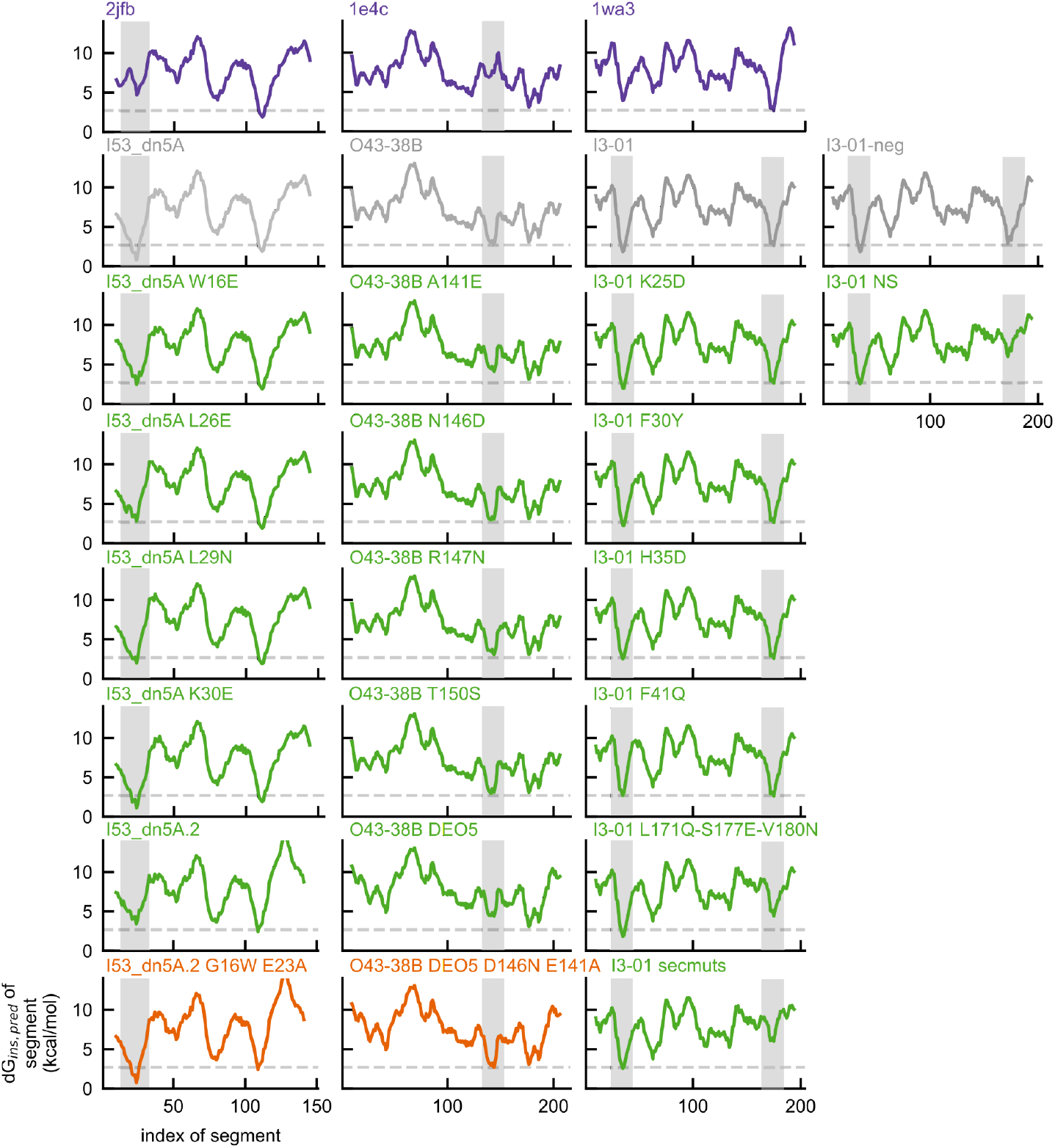
dG_ins,pred_ of proteins characterized for secretion yield in Figures 2 and 3. Effect of degreaser-identified mutations and orthogonal redesign (green), as well as the reversion of degreaser-identified mutations (orange) on the lowest dG_ins,pred_ segment in the sequence in comparison to the original designs (gray) and their naturally occurring counterparts (purple). Regions corresponding to the lowest dG_ins,pred_ segment(s) of the original designed proteins are shaded in gray. The horizontal dashed line indicates the +2.7 kcal/mol threshold.

**Supplementary Figure S6:**
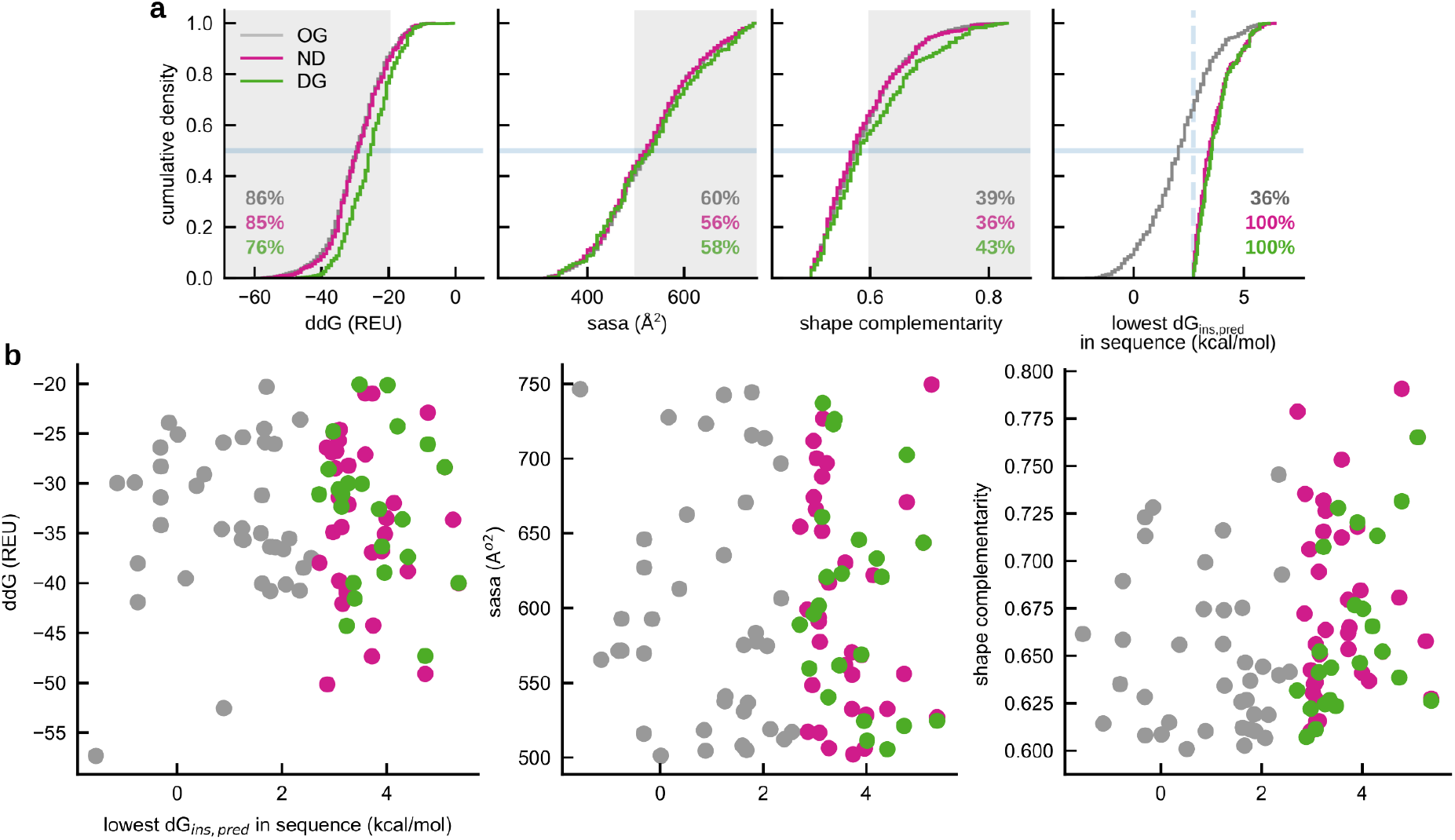
Design metrics do not substantially differ between KWOCA OG, ND, and DG designs. (a) Distributions of design metrics for each design set. Shaded regions highlight the proportion of designs accepted for each metric. Proportions of design with dG_ins,pred_ >+2.7 kcal/mol (dotted line) resulting from each protocol are also shown. (b) The distribution of lowest dG_ins,pred_ does not correlate with other design metrics in KWOCA designs chosen for experimental characterization (see Figure 4).

**Supplementary Figure S7:**
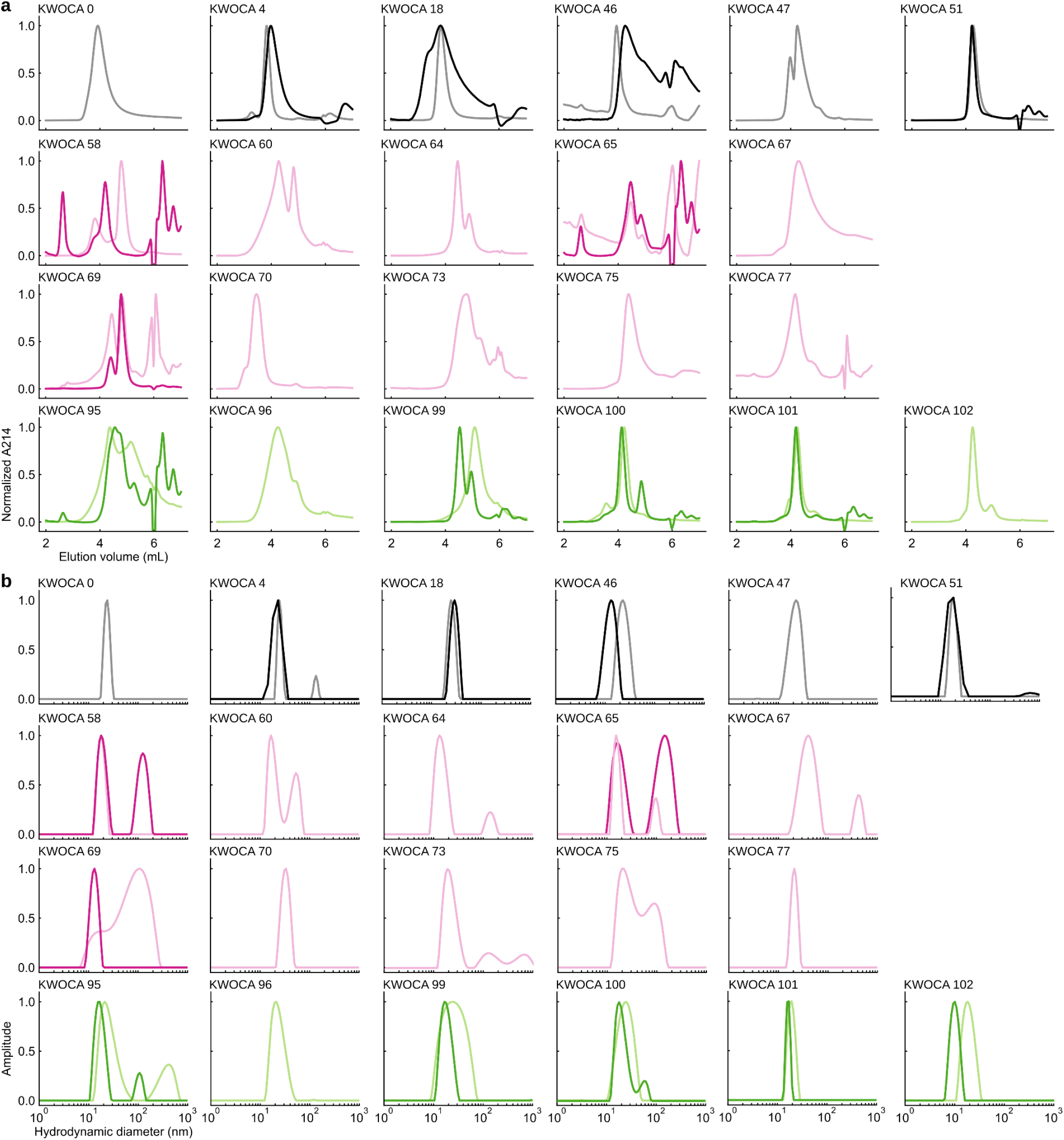
SEC and DLS suggest that 22 of 99 KWOCAs self-assemble. (a) SEC and (b) DLS profiles of proteins purified from bacterial cell lysates (light color) and clarified supernatants of mammalian cells (dark color) are shown. Constructs purified both from bacterial and mammalian material generally showed similar profiles.

**Supplementary Figure S8:**
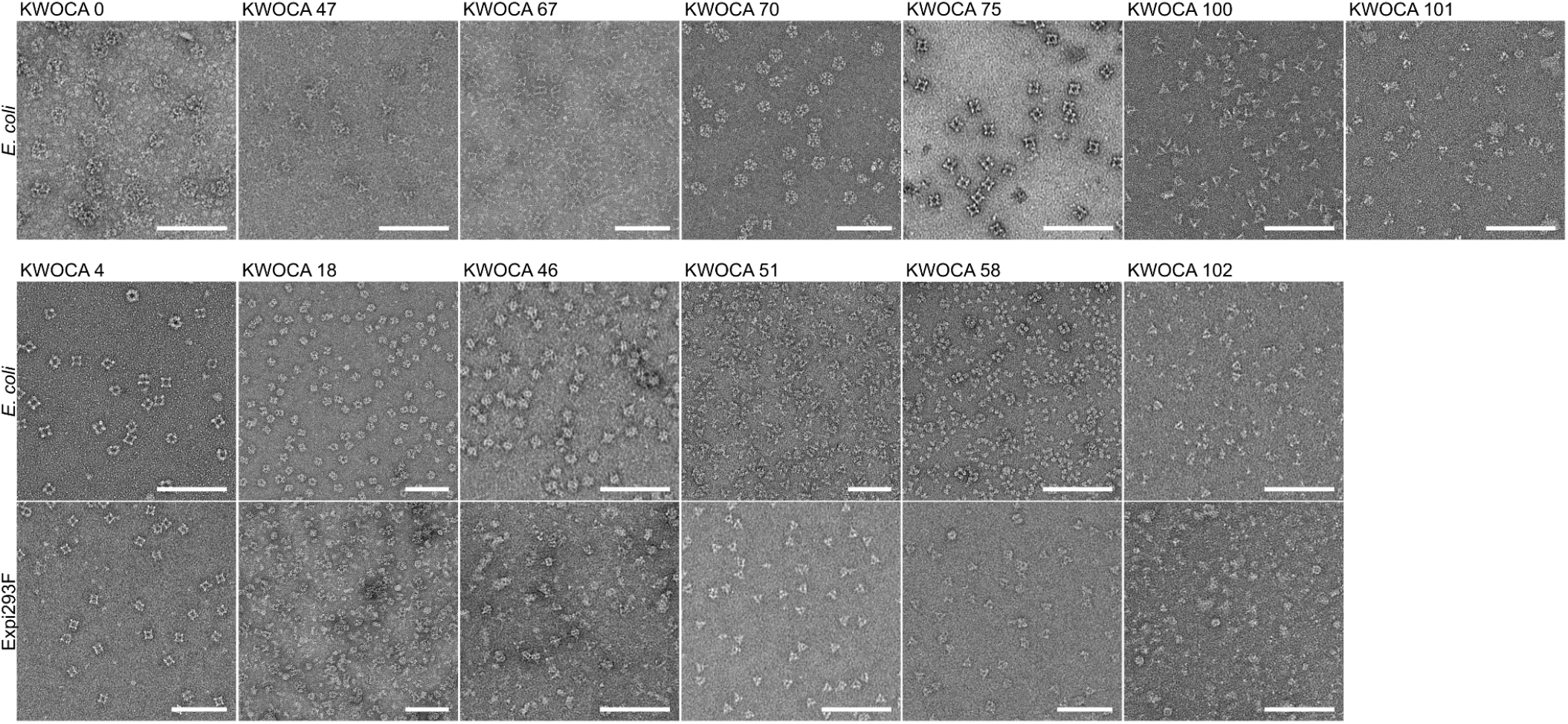
Confirmation of 13 of 22 potentially assembling KWOCAs by nsEM. Constructs purified from mammalian material (Figure 4d) are indistinguishable from those purified from bacterial material. Scale bar, 100 nm.

**Supplementary Figure S9:**
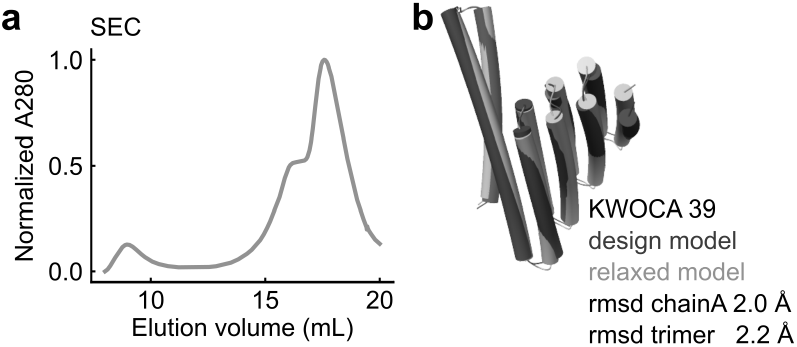
SEC and crystal structure of a non-assembling KWOCA. (a) SEC profile and (b) crystal structure of the non-assembling KWOCA 39. The structure is aligned to the backbone of a single subunit of the computational design model, and rmsd values for the monomer and trimer are shown.

**Supplementary Figure S10:**
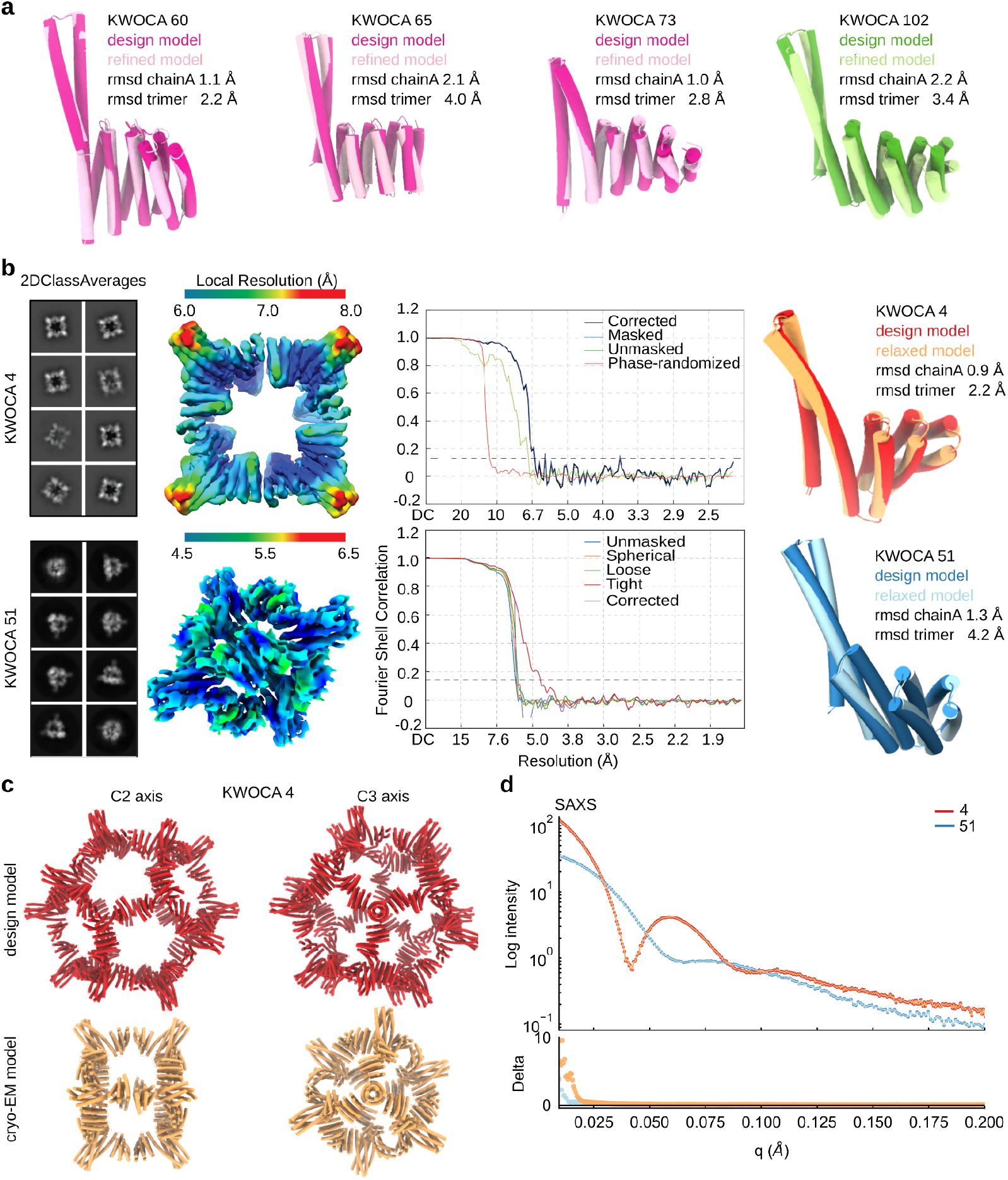
Additional structural information on assembling KWOCAs. (a) Crystal structures of the trimeric building blocks of KWOCAs 60, 65, 73 (possible assemblies, although not validated by nsEM) and 102 (nsEM-validated assembly) overlaid with their corresponding design models, aligned by the backbone of the single monomer shown. (b) Representative 2D class averages, local resolution estimation maps, and FSC curves for the cryo-EM densities displayed in Figure 5. Overlays of the monomer of each design model with the fitted model relaxed into the cryo-EM density are also shown. (c) KWOCA 4 icosahedral design model and octahedral relaxed model obtained from fitting eight copies of the trimer to the cryo-EM density. (d) Theoretical SAXS profiles (dotted dark lines) calculated from the cryo-EM models are overlaid with the experimentally obtained profiles for KWOCA 4 and KWOCA 51.

**Supplementary Figure S11:**
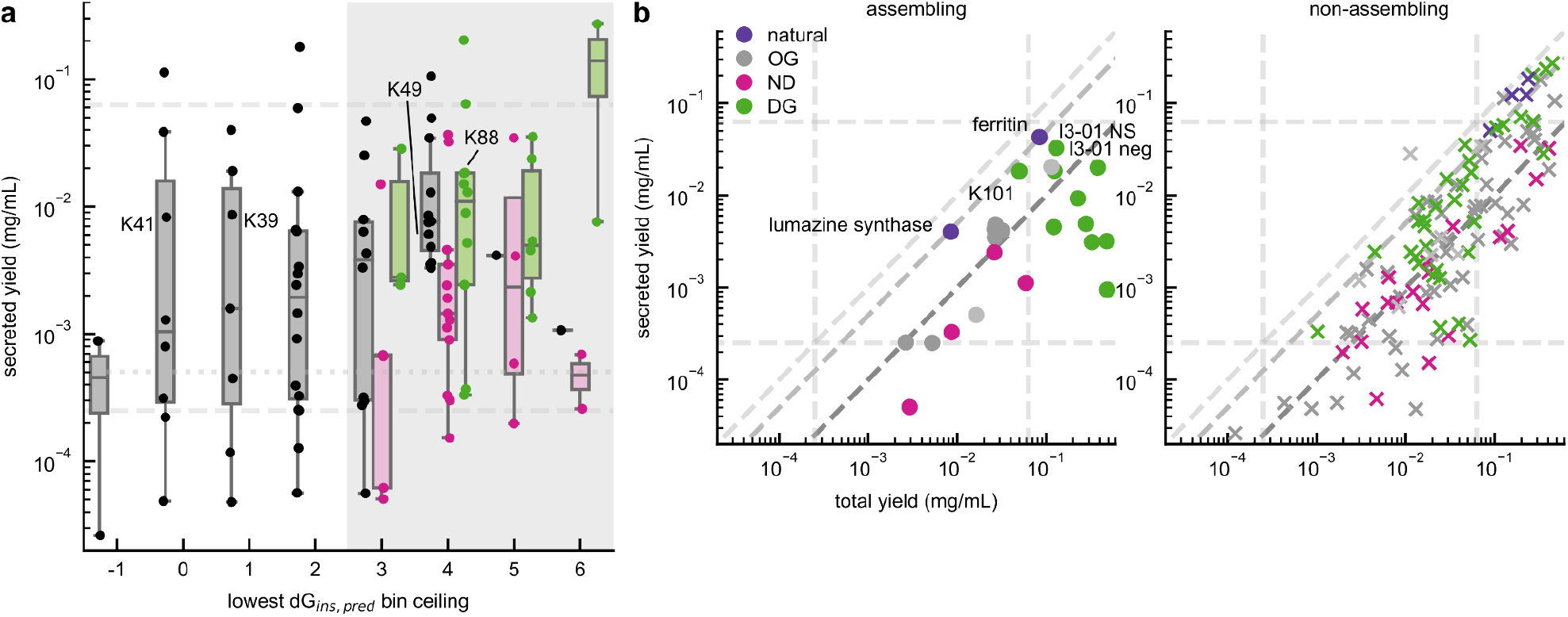
Degreased proteins have improved total yield and secretion efficiency. (a) Secreted yield vs dG_ins,pred_ for non-assembling KWOCAs, binned in 1 kcal/mol increments. The boxes represent median and quartile values, while the whiskers indicate data points within 1.5 inter-quartile ranges. (b) Secreted yield vs total yield for natural protein nanoparticles, I3-01 variants, and assembling and non-assembling KWOCAs. For some constructs, the total yield values, which represent the summed yield of intracellular and secreted protein obtained by western blot, were above the linear calibration curve (bounds represented by dashed lines). Diagonal dashed lines represent secretion efficiencies (secreted yield divided by total yield) of 1, 0.5, and 0.1 (top left to bottom right).

**Supplementary Table S1.**
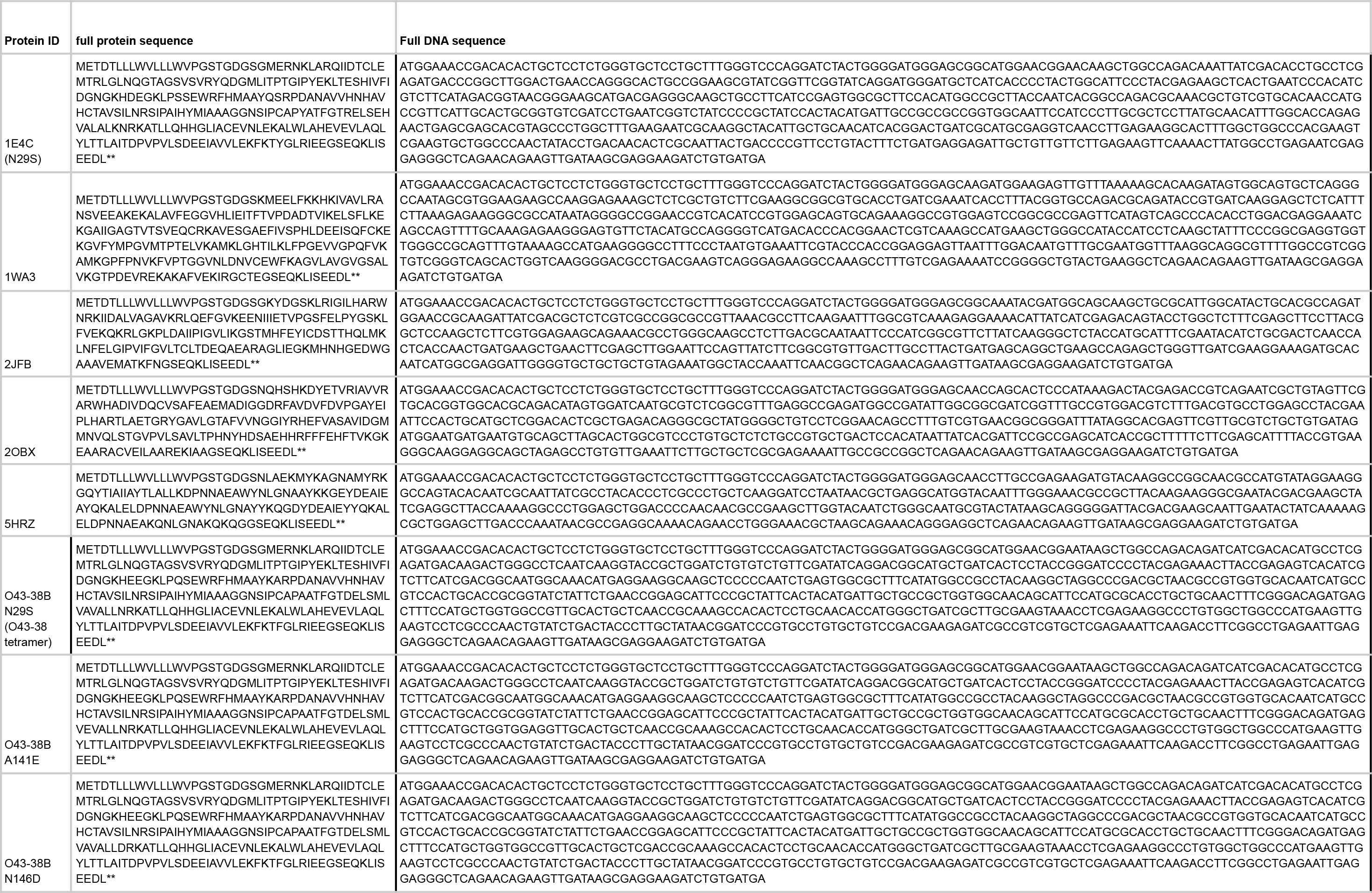
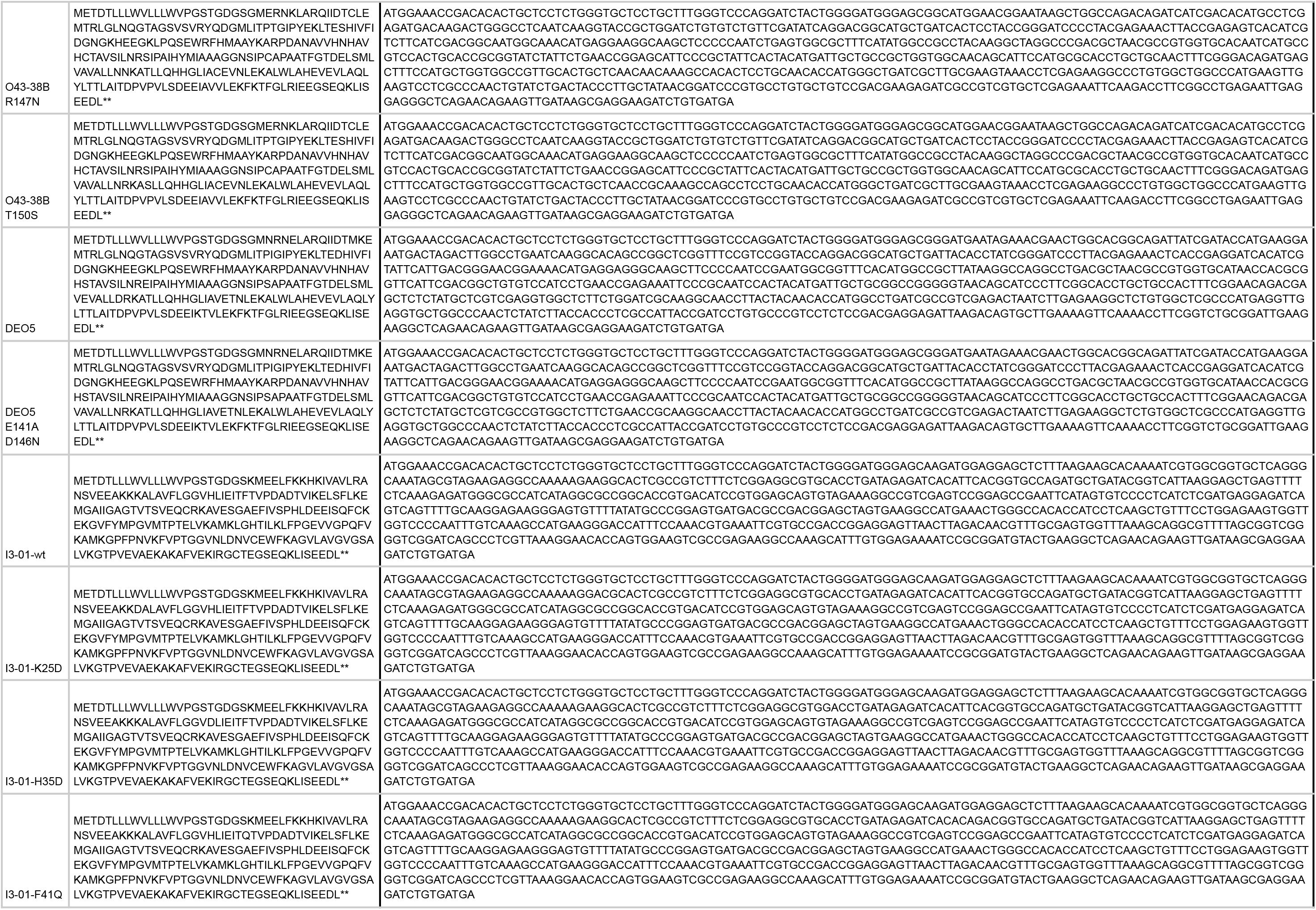
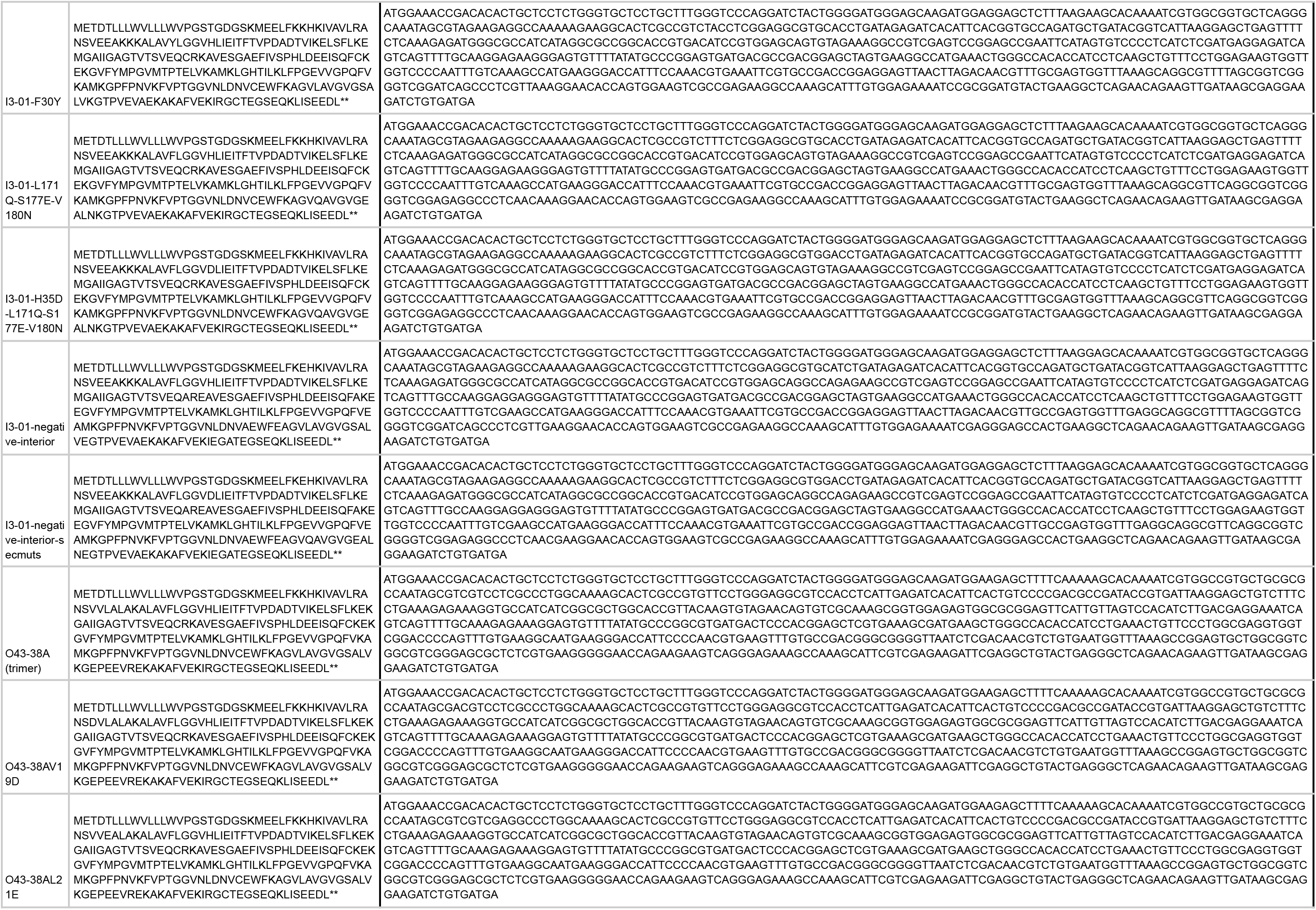
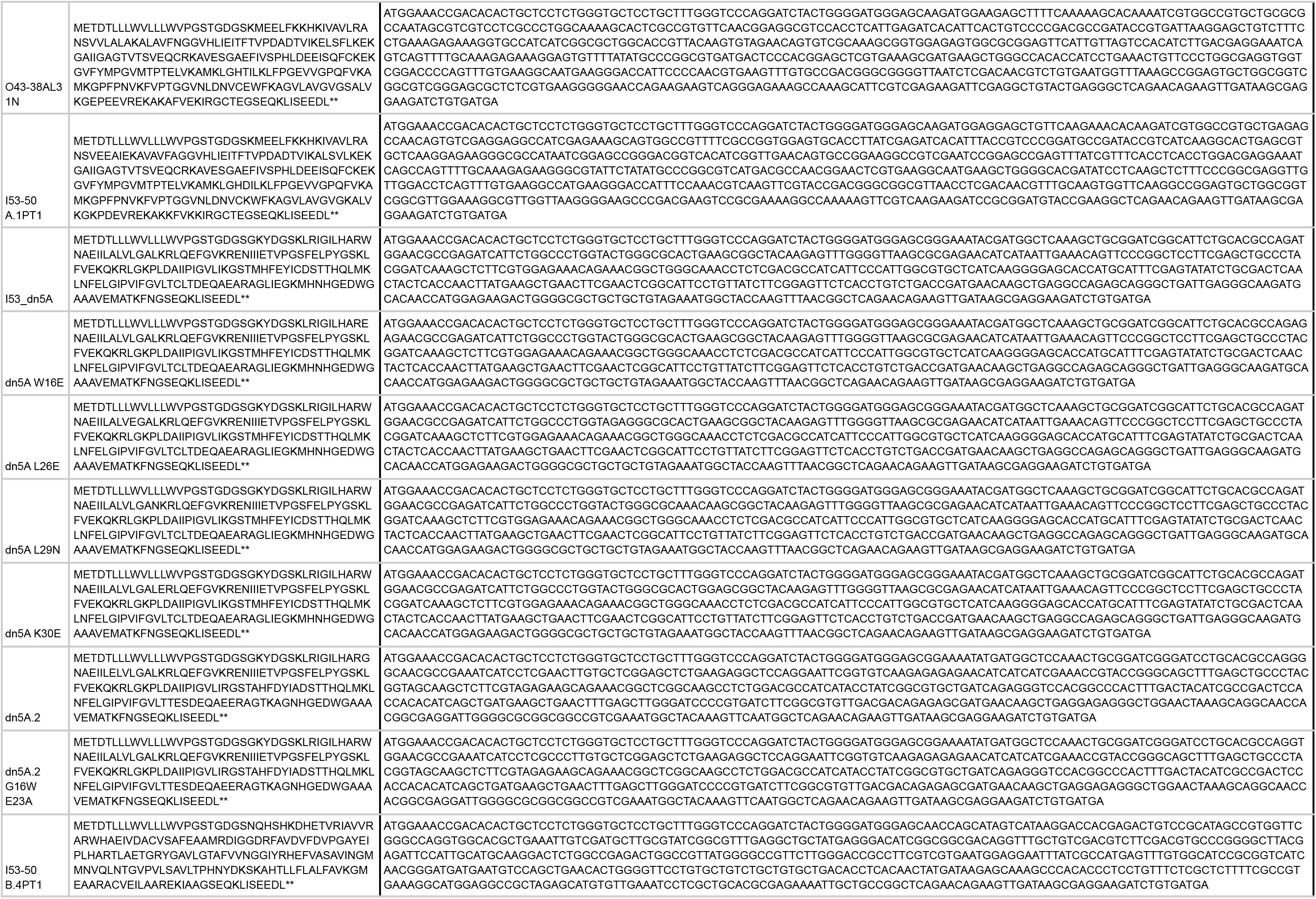
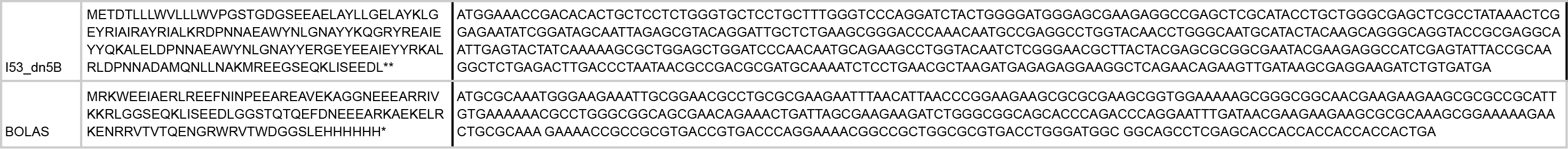
Full protein and DNA sequences for mammalian protein expression of retrospectively Degreased previously designed nanoparticle and nanoparticle components, as well as their designed and natural counterparts.

**Supplementary Table S2.**
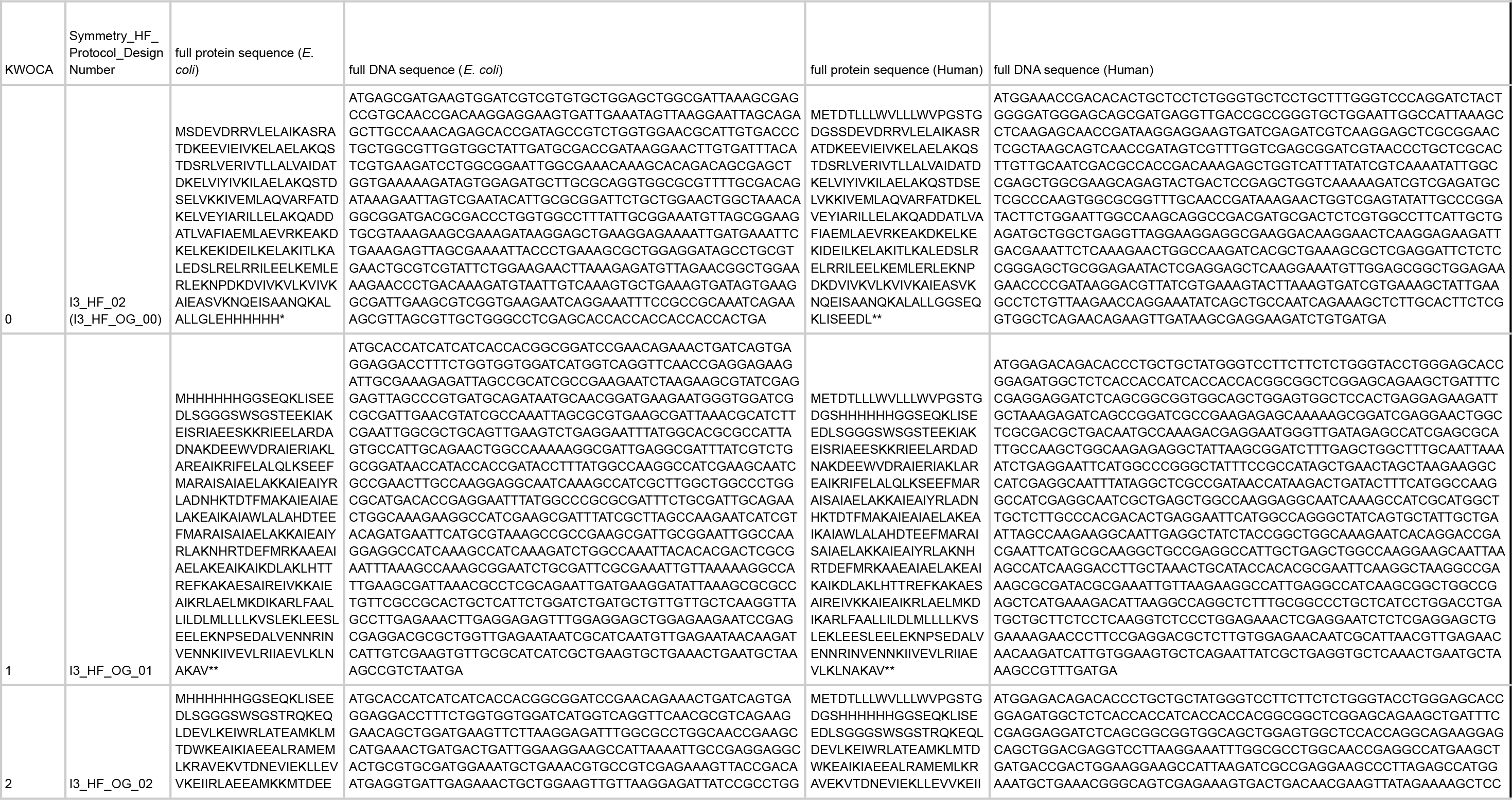
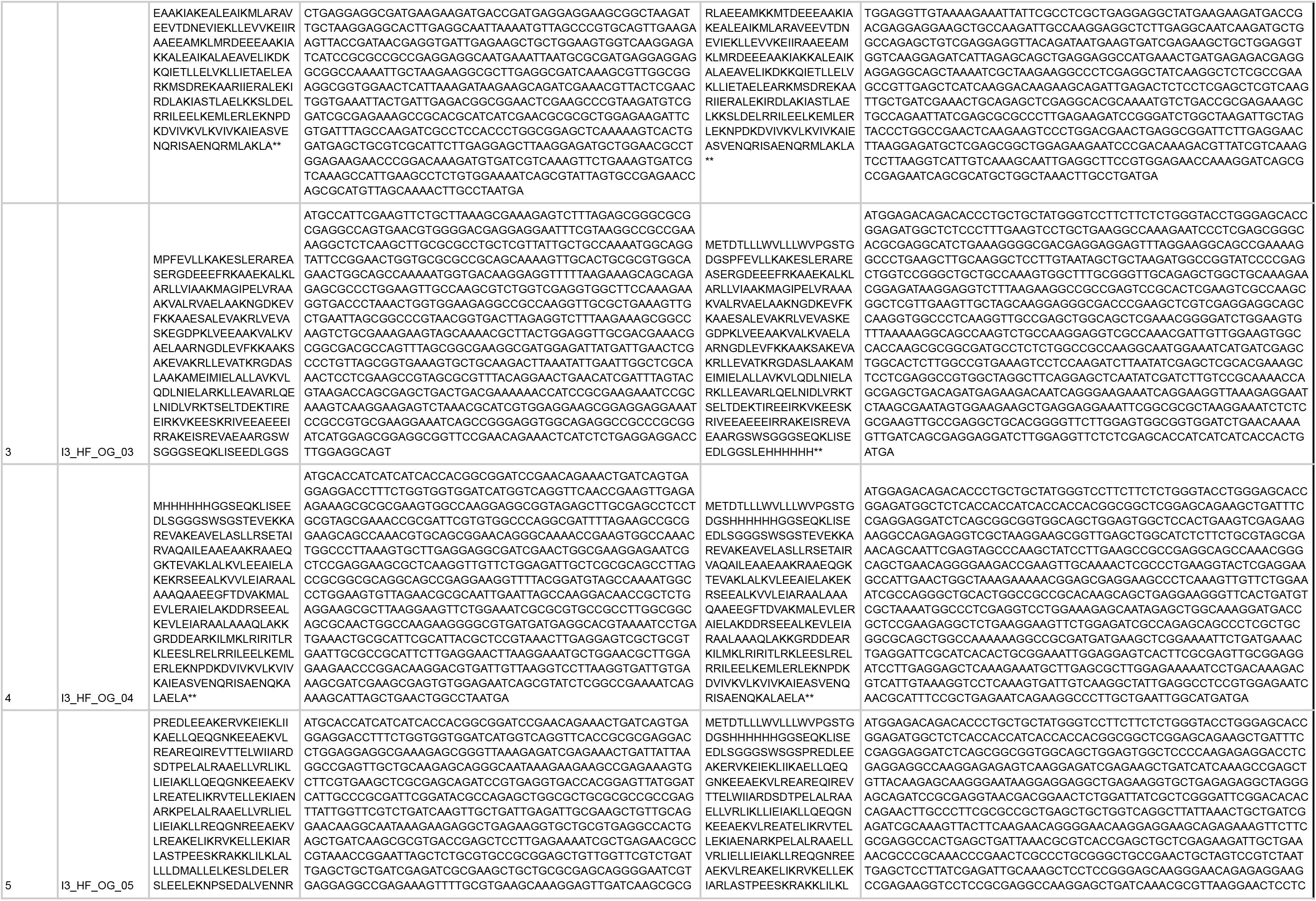
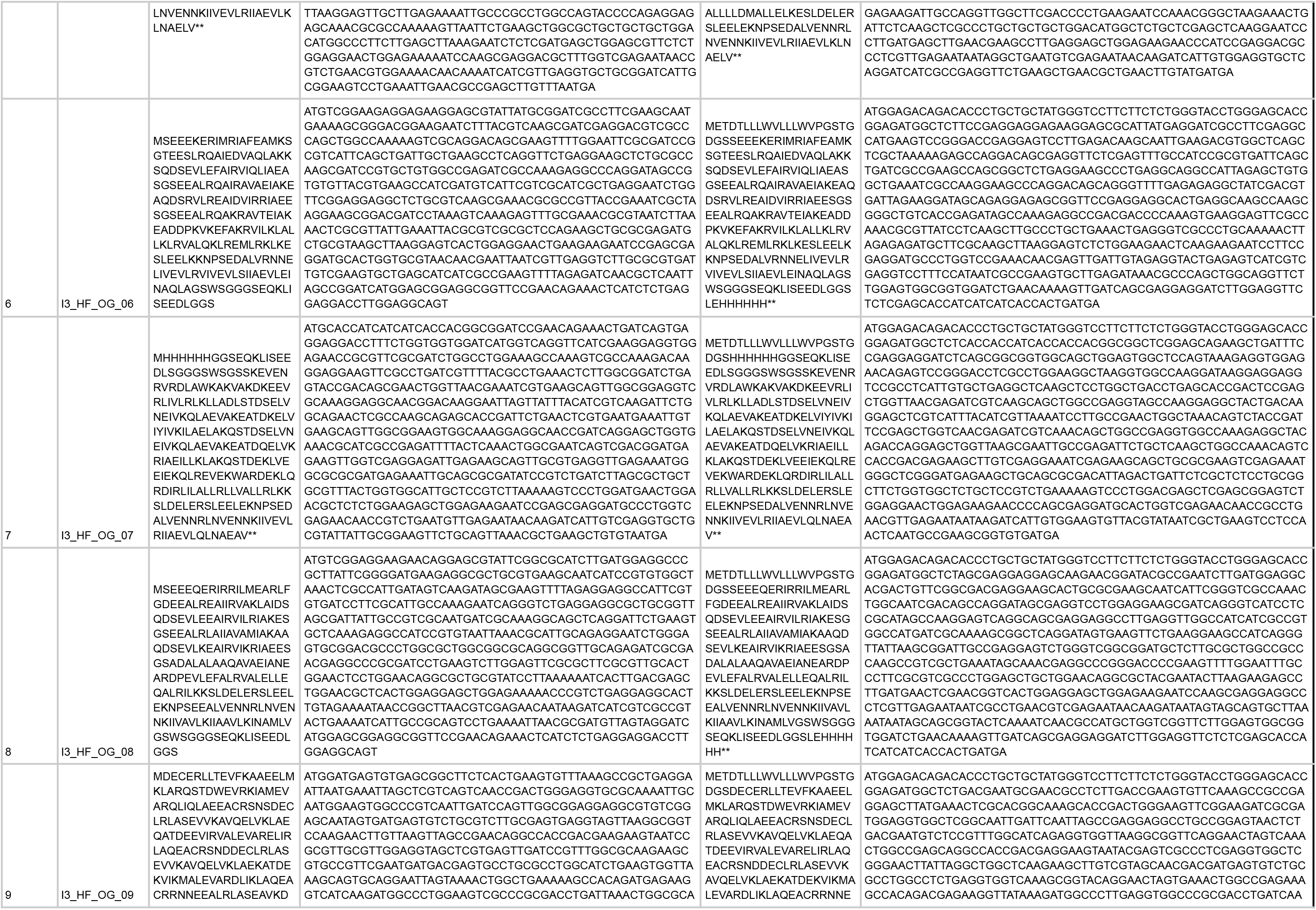
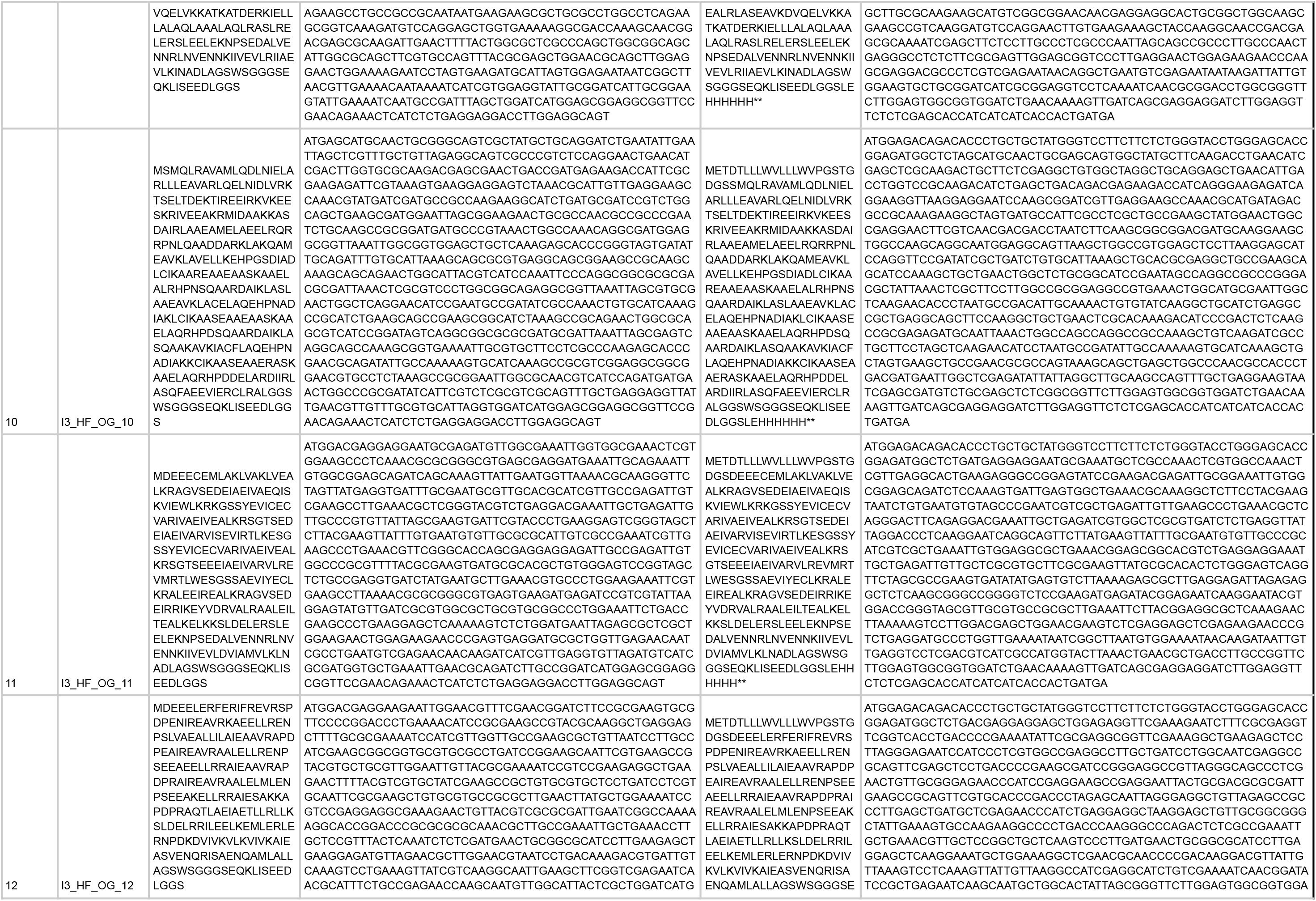
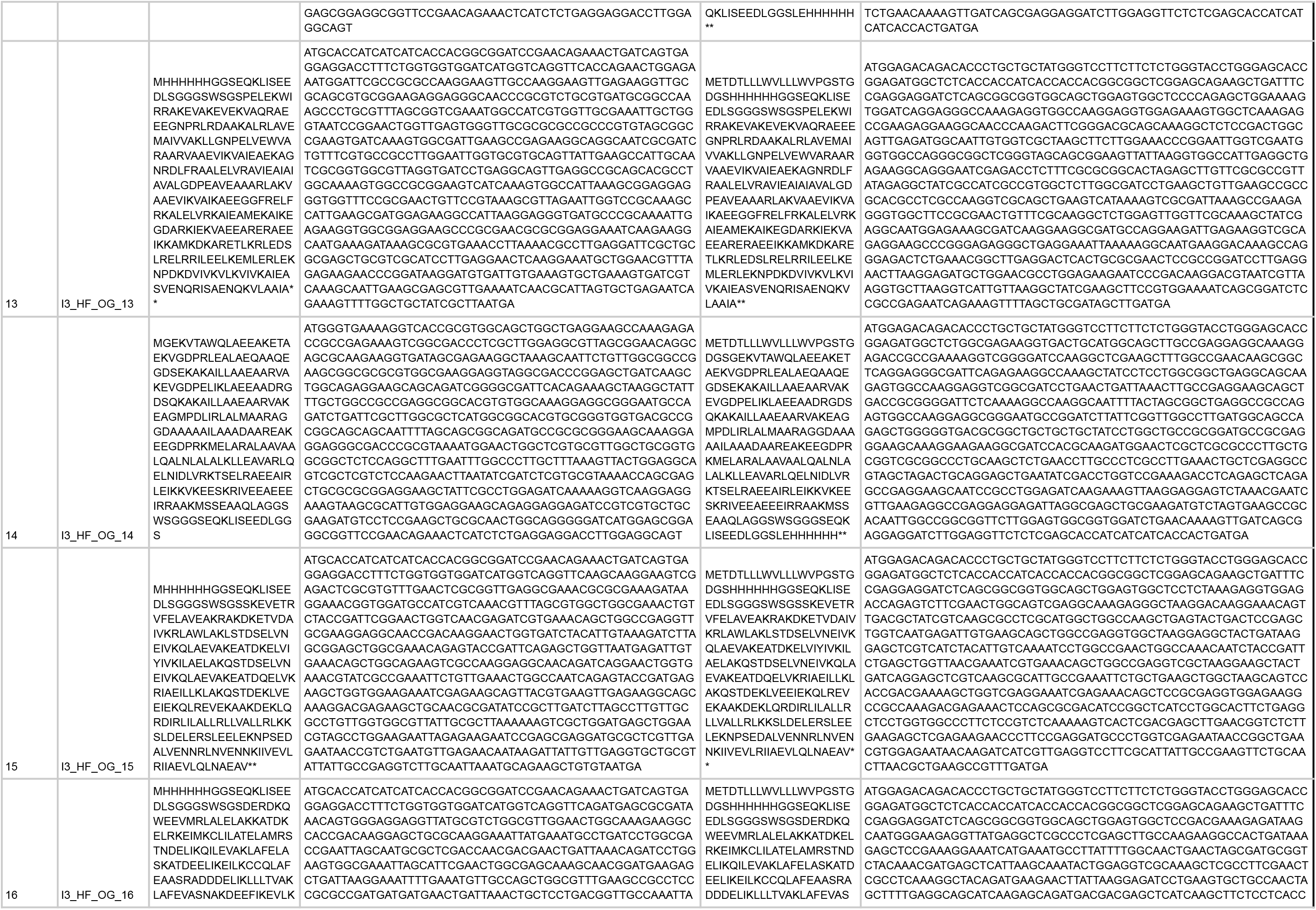
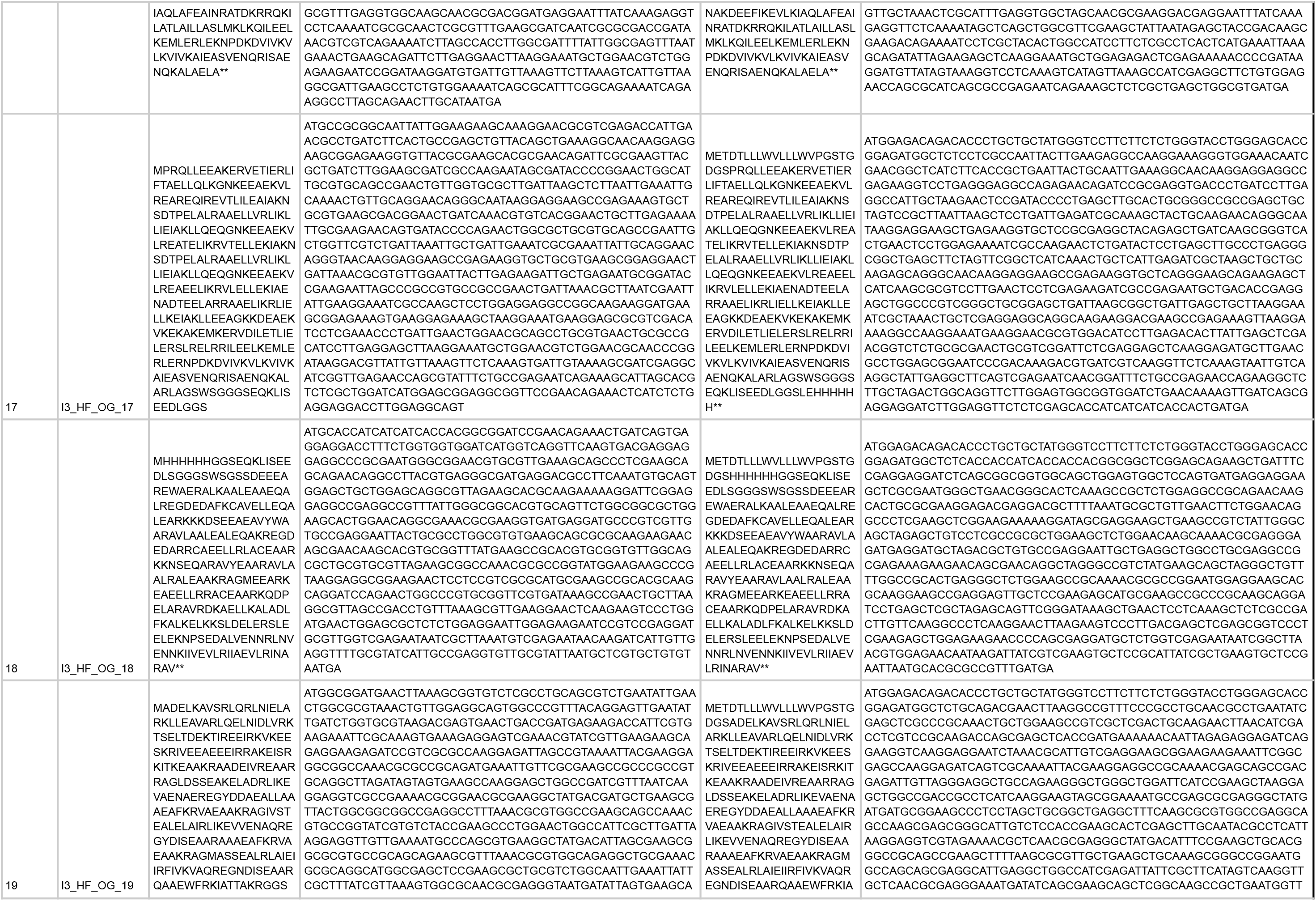
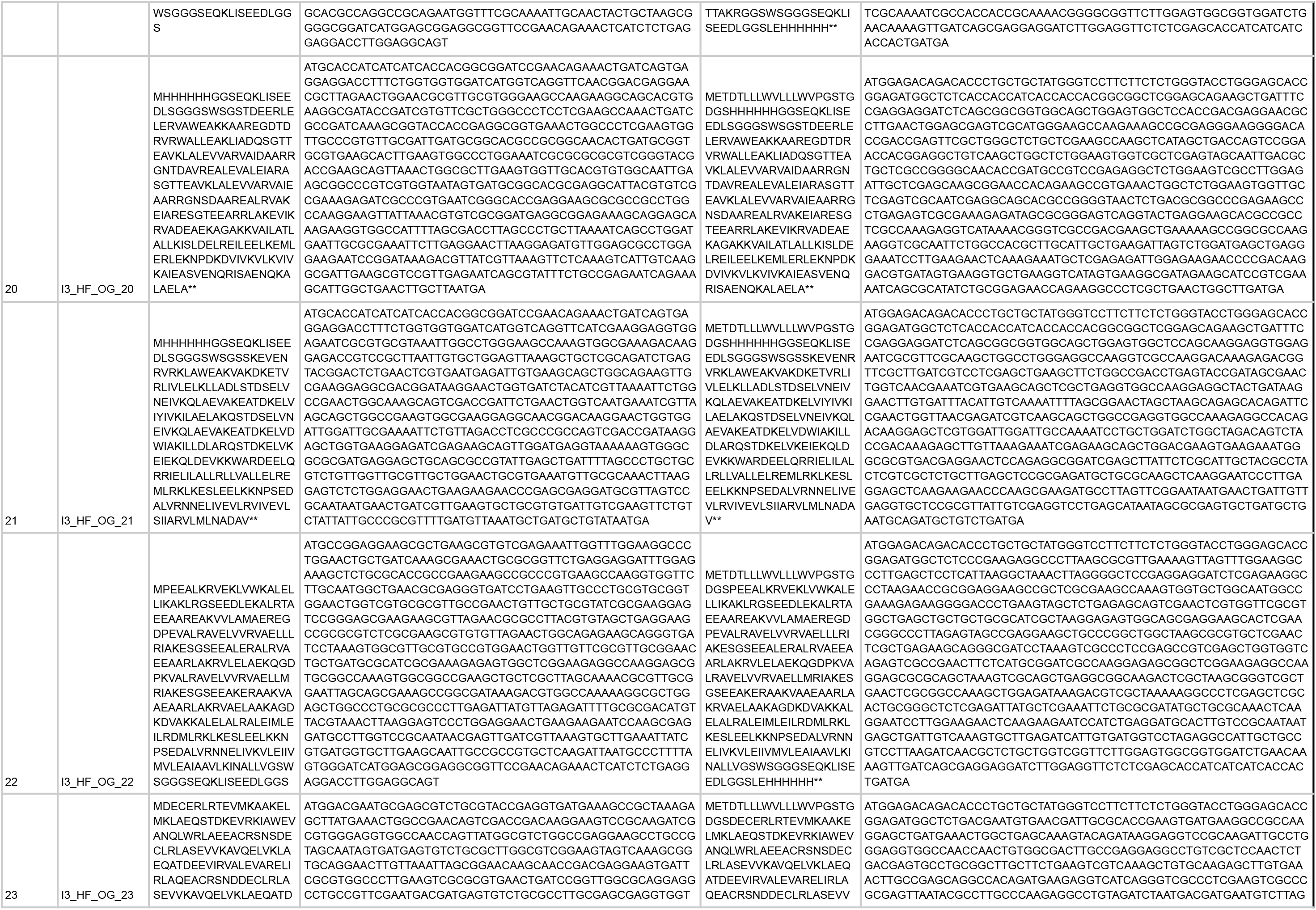
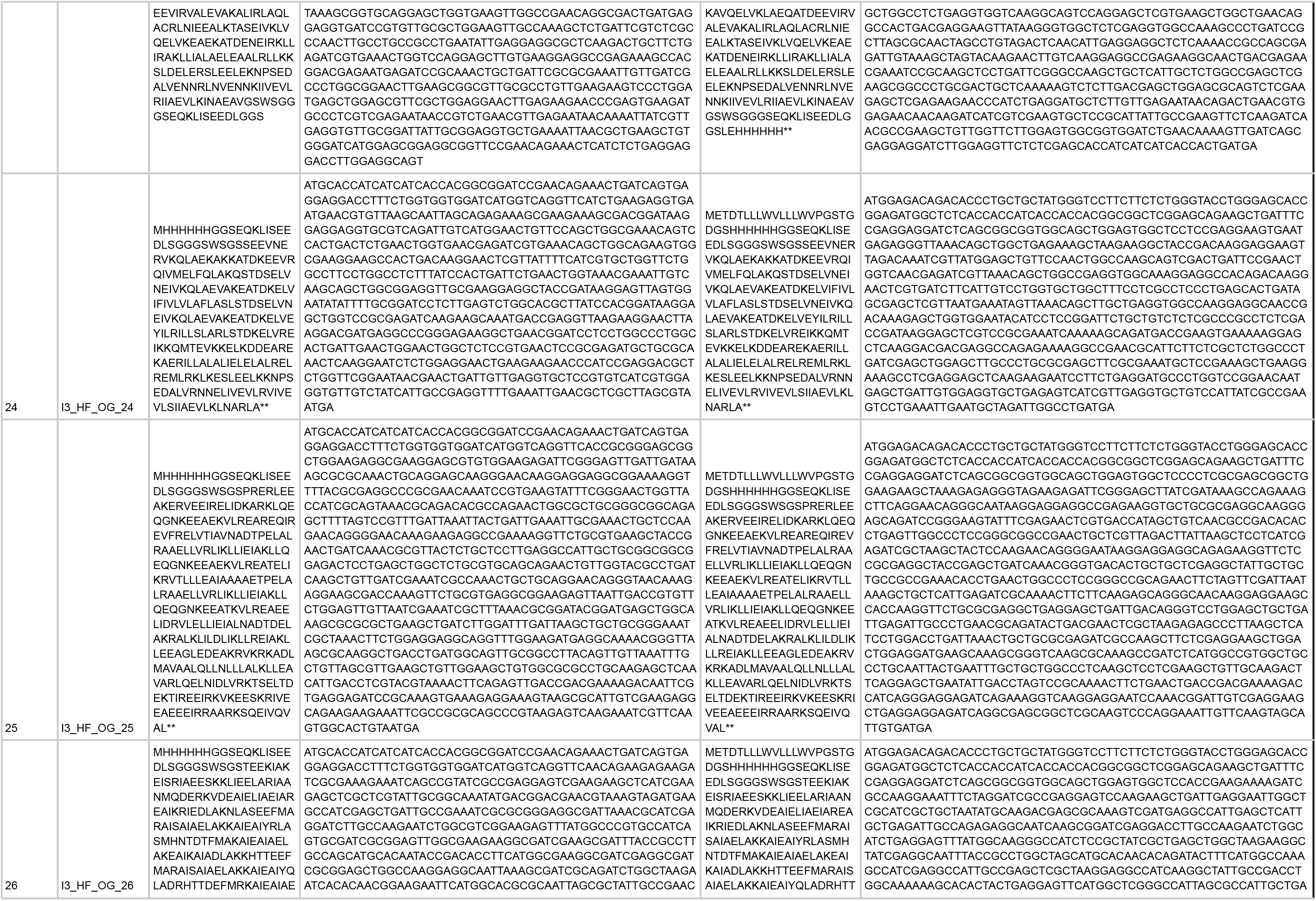
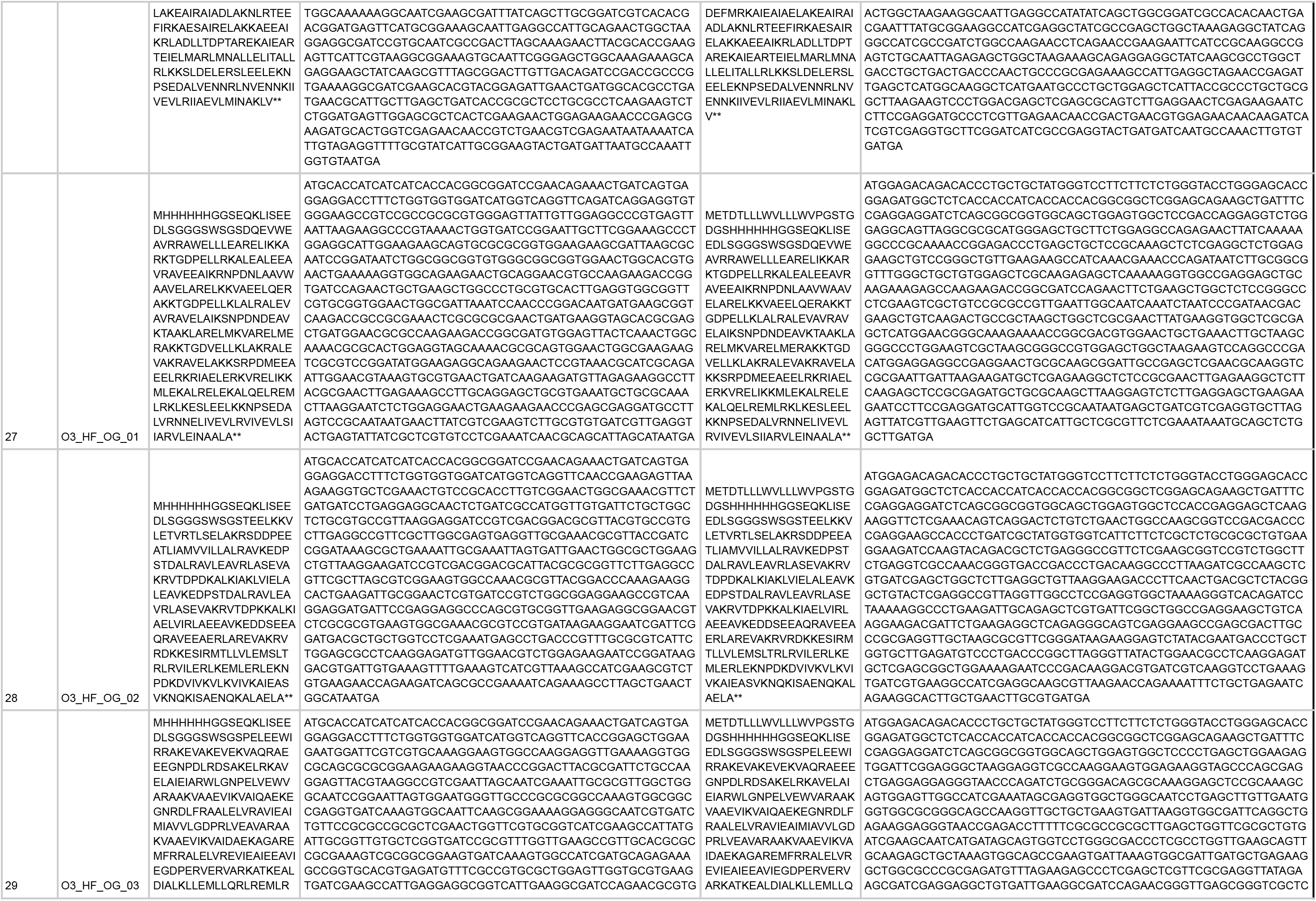
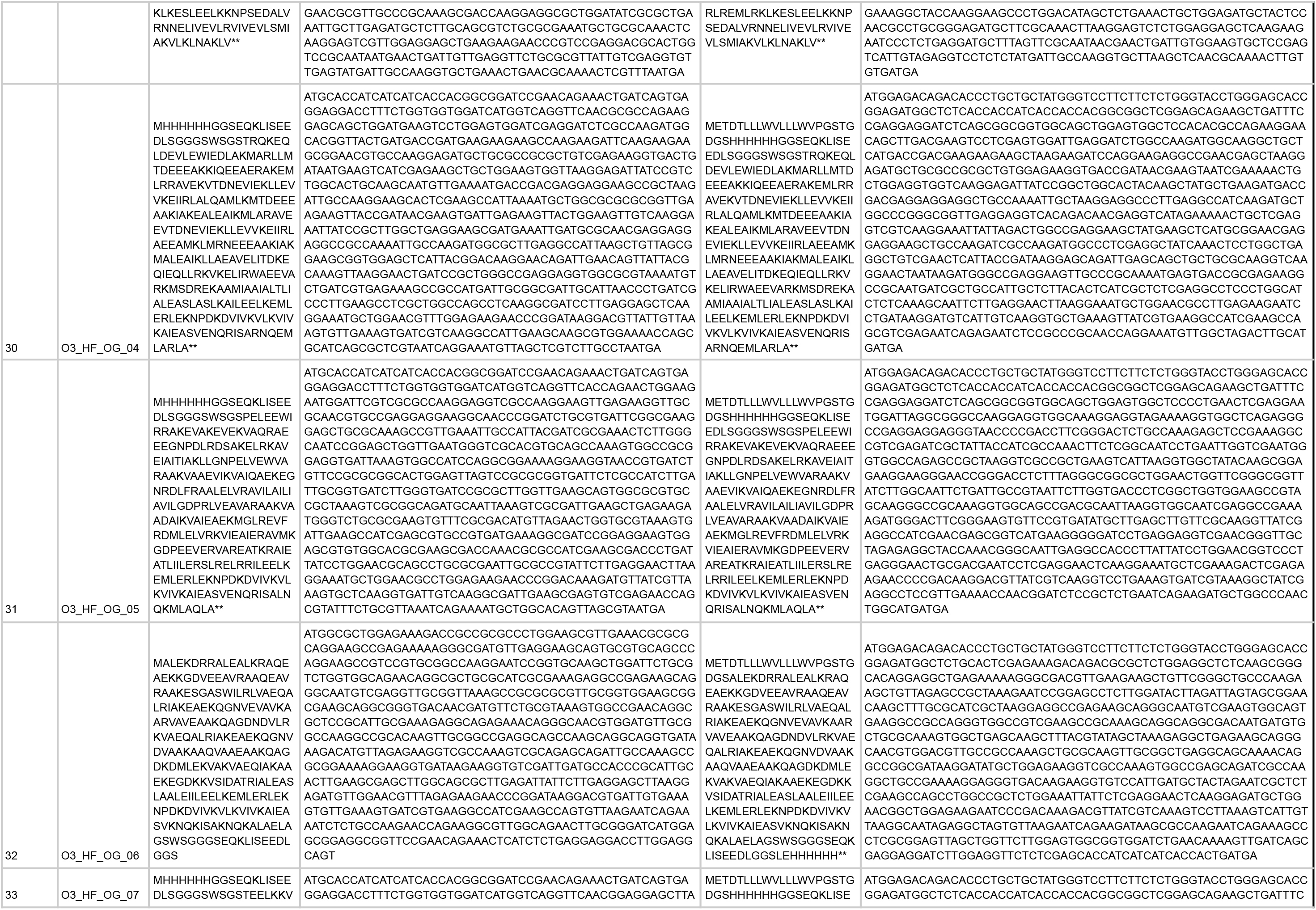
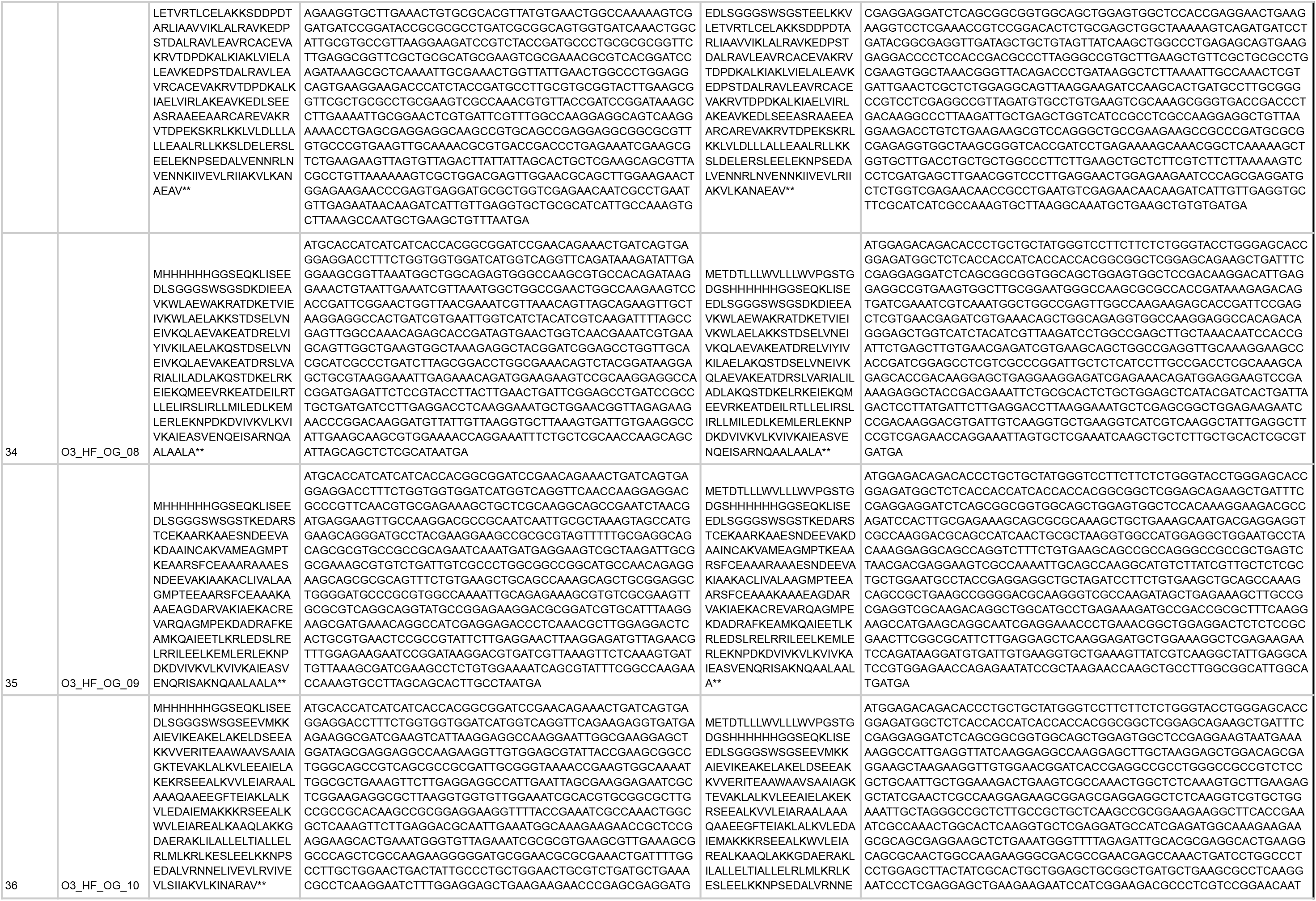
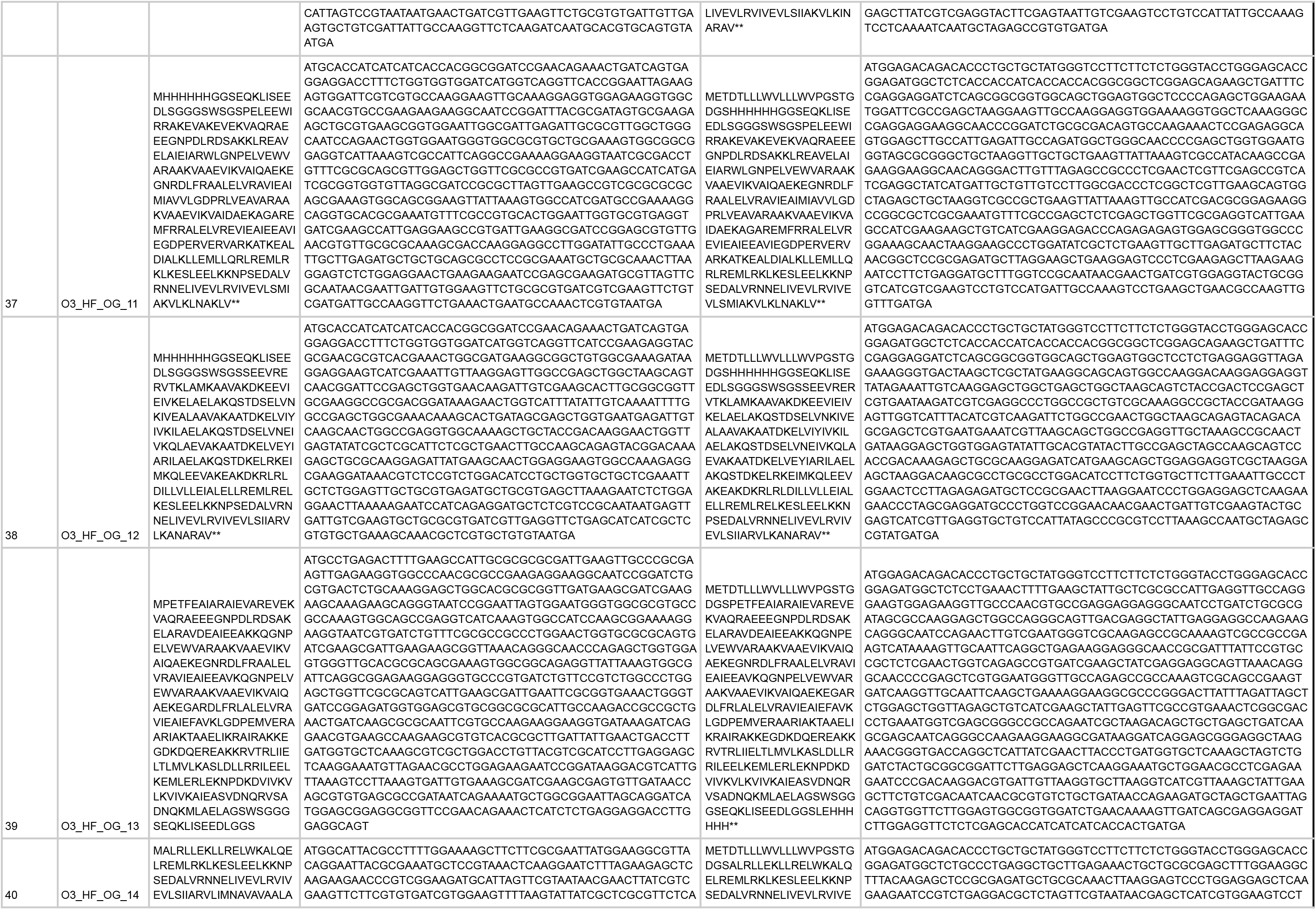
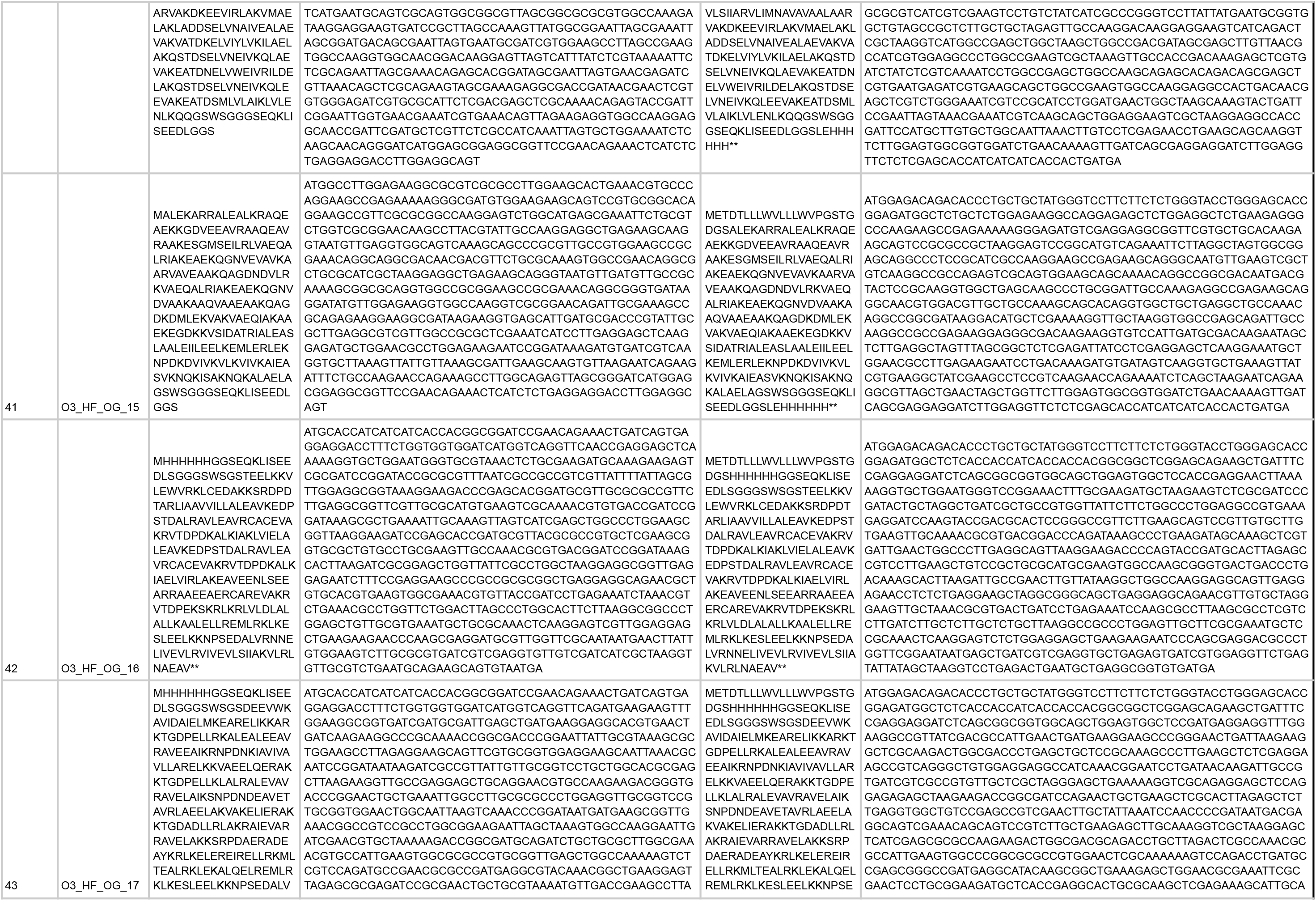
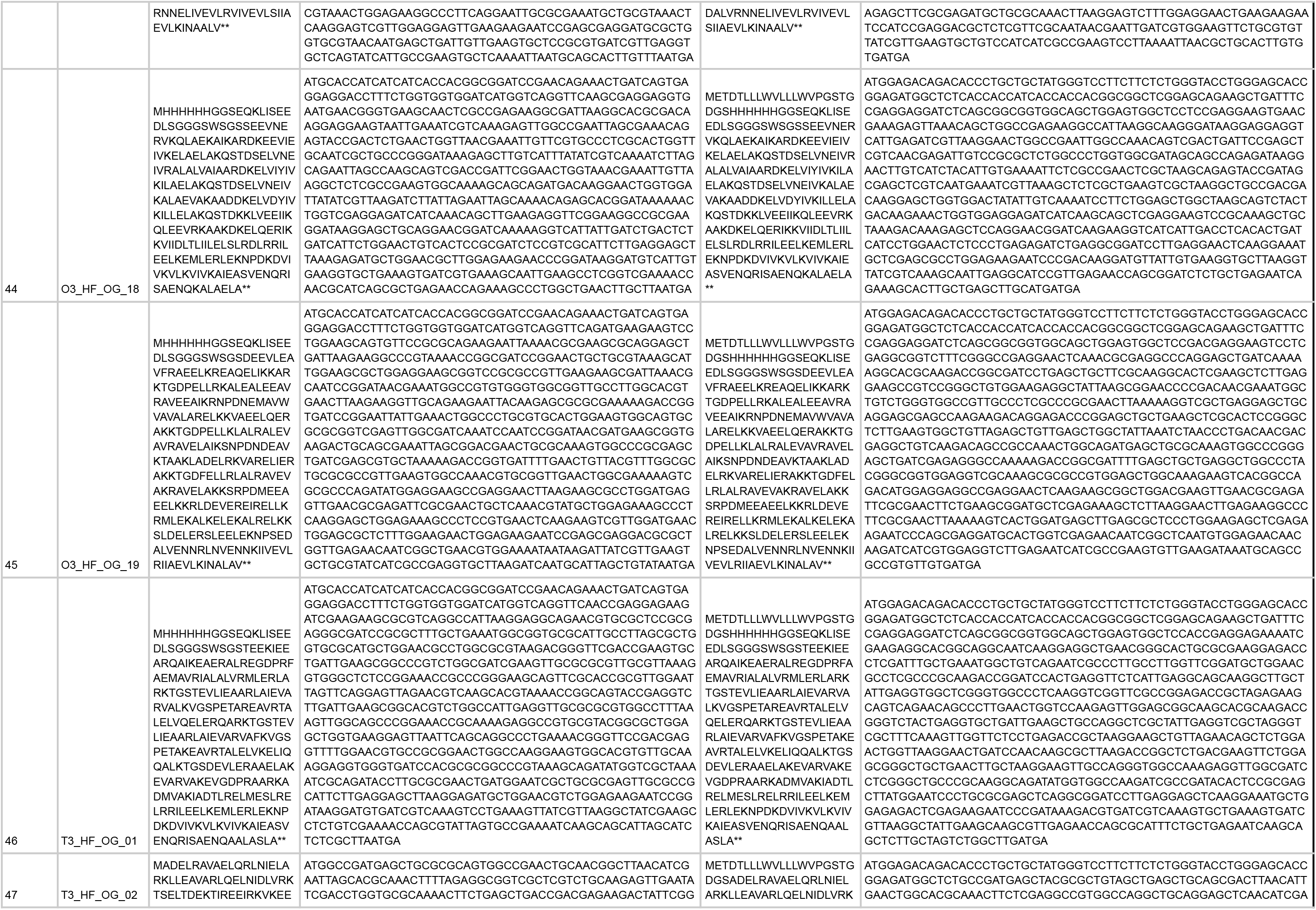
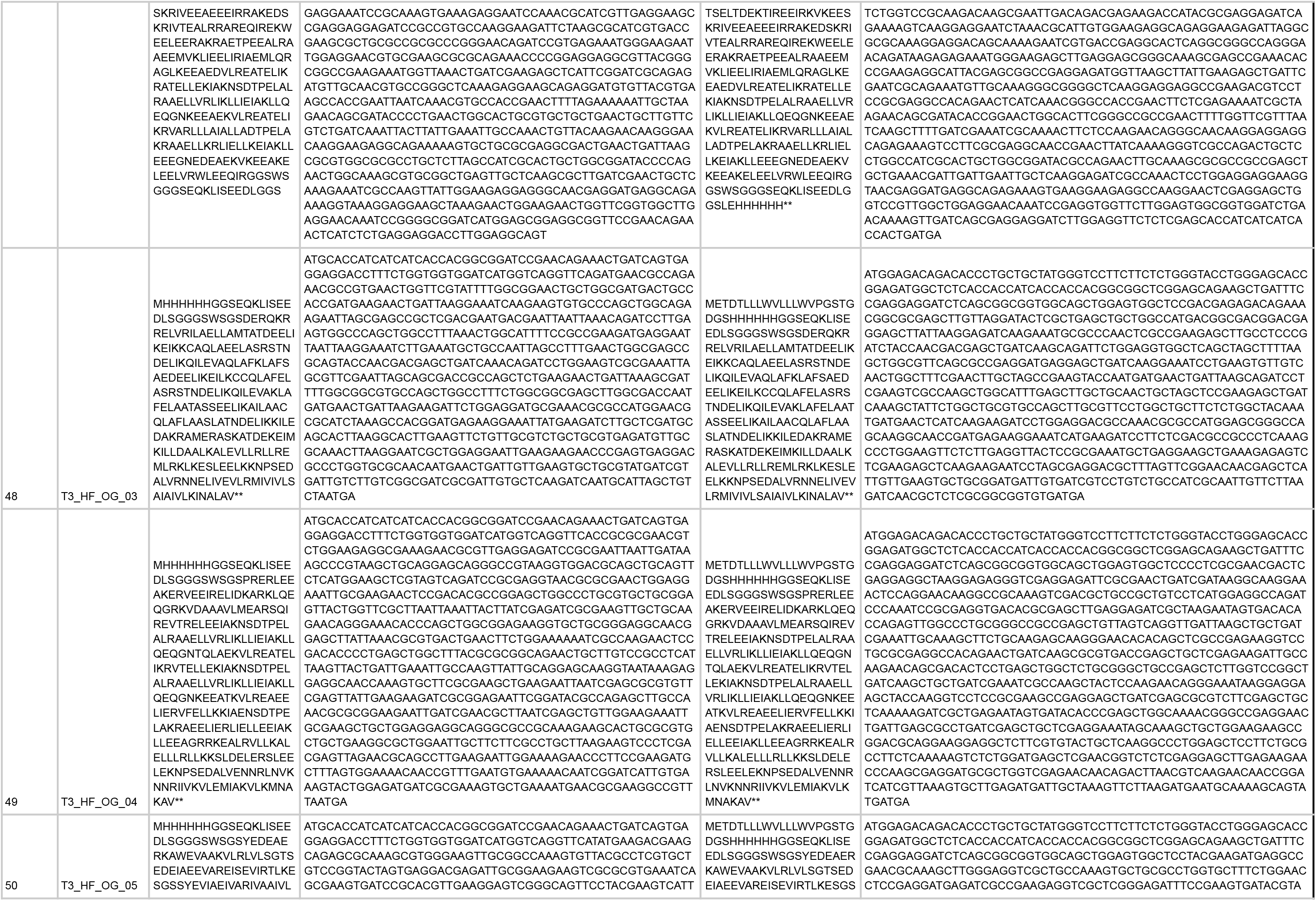
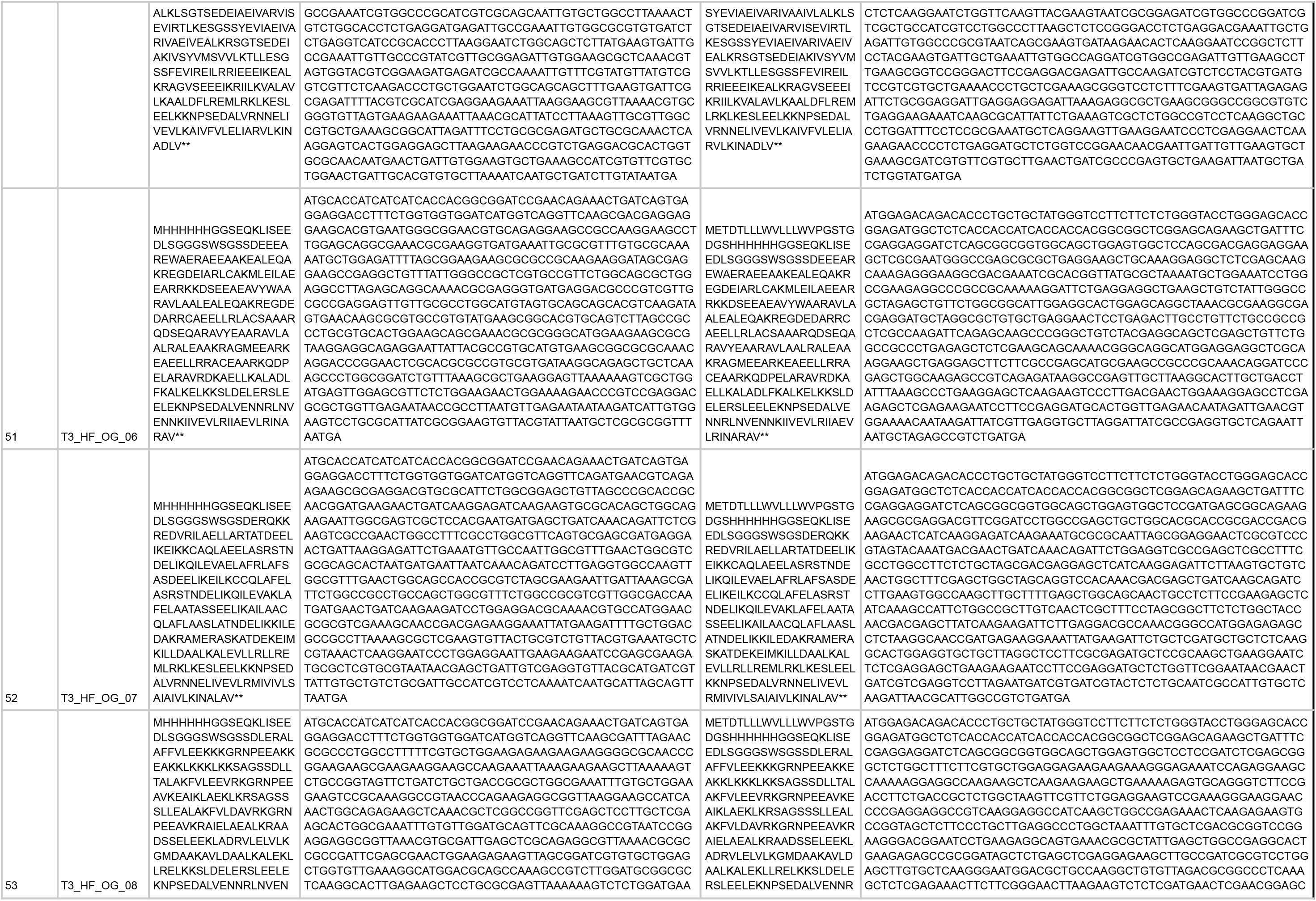
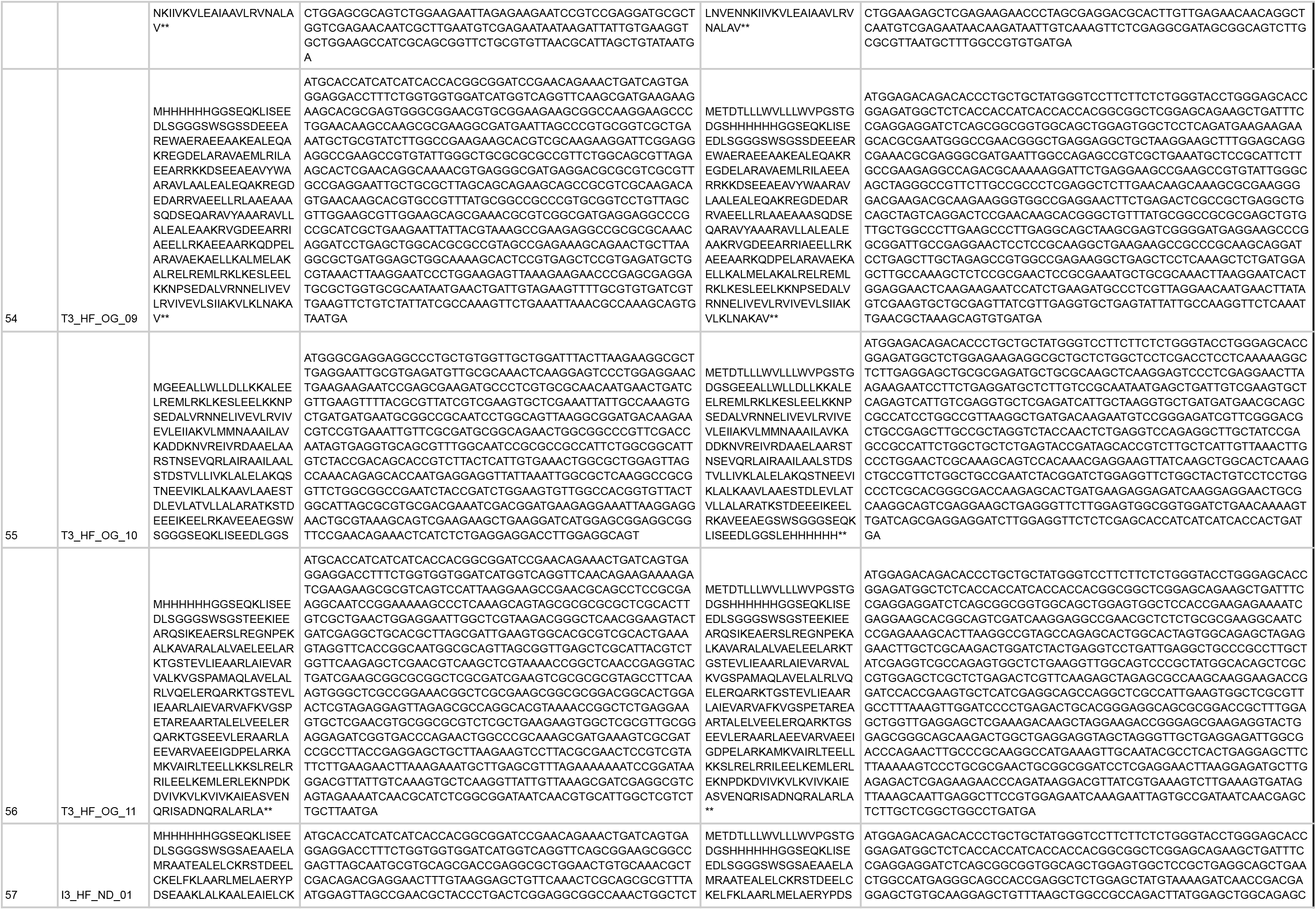
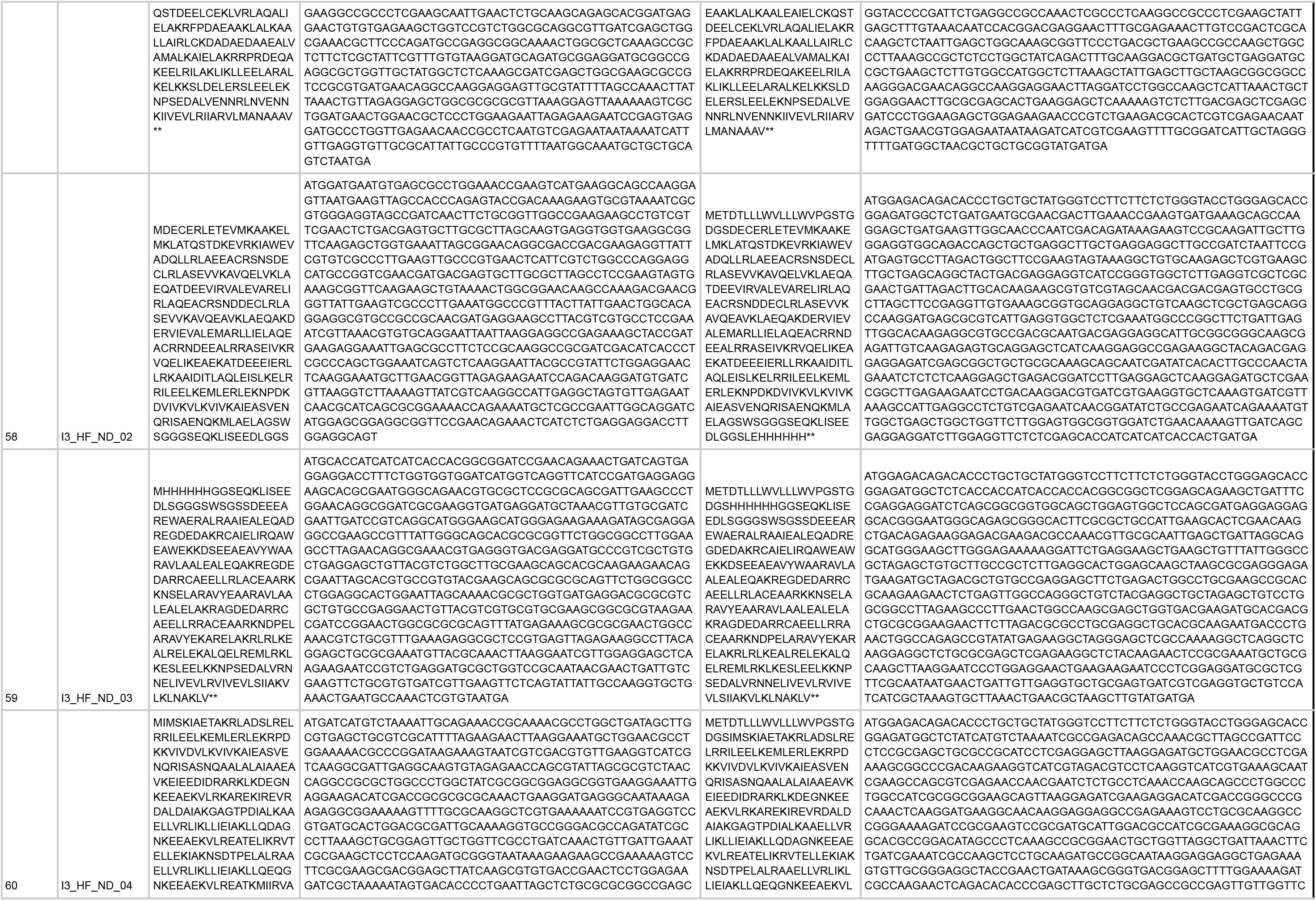
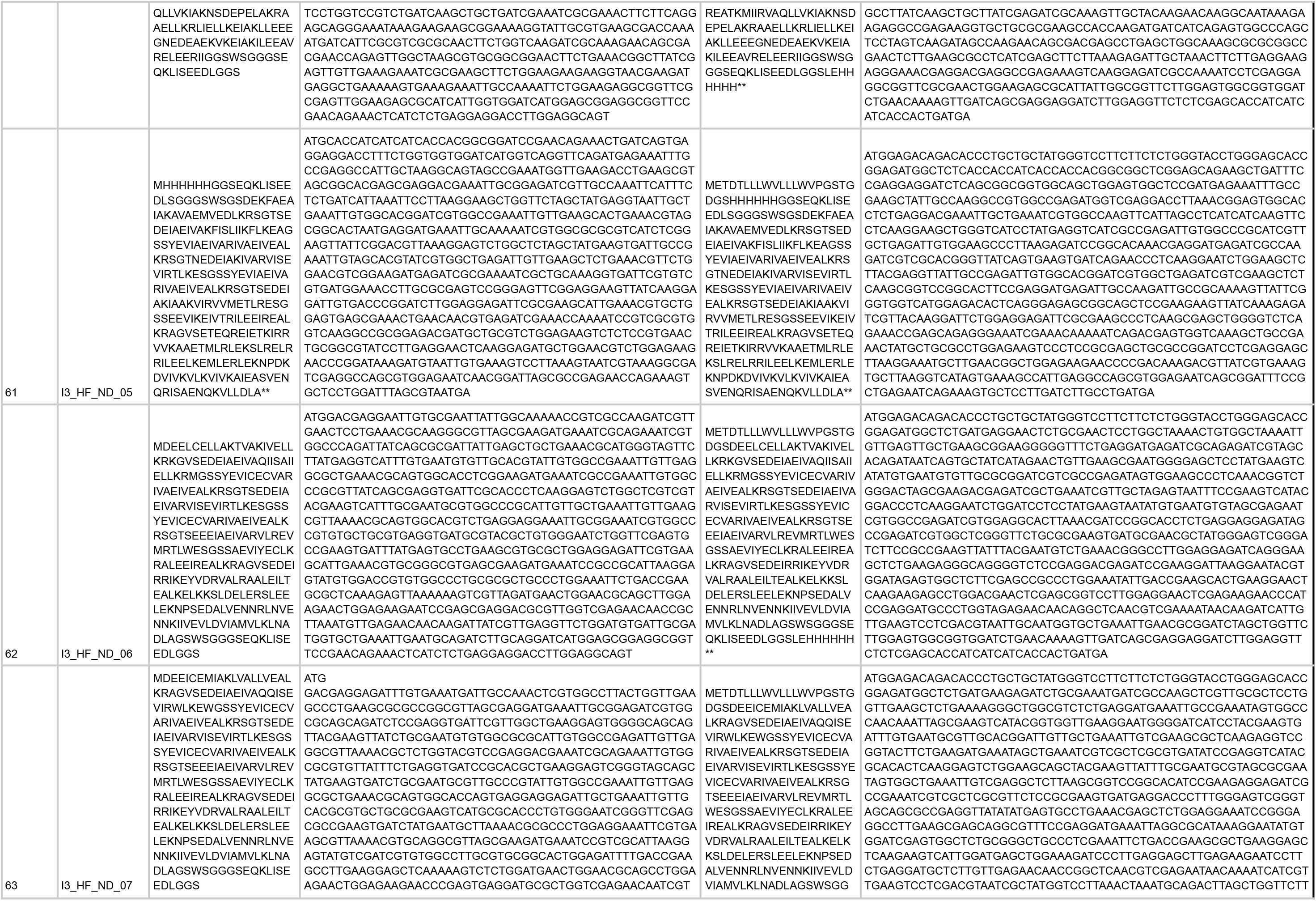
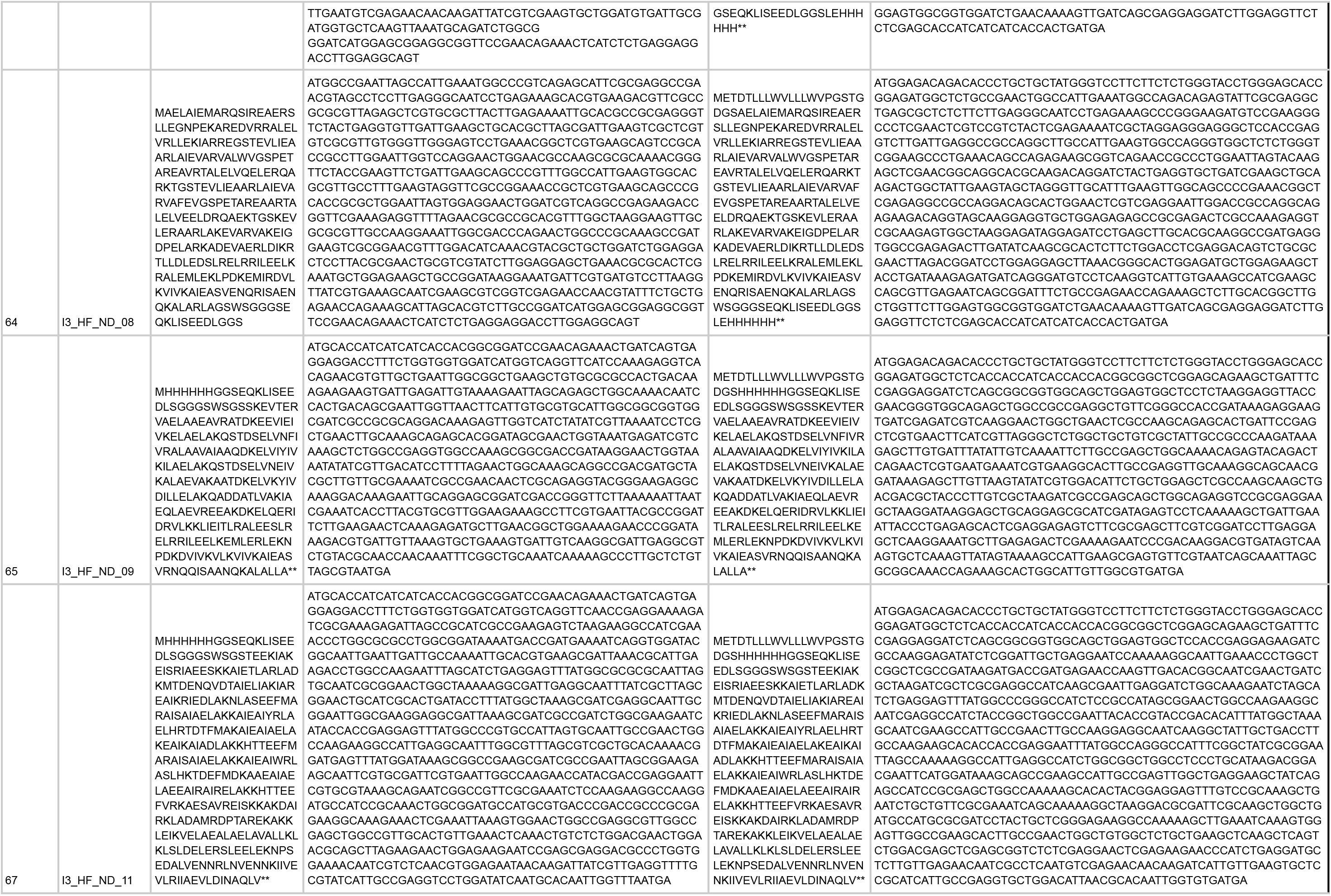
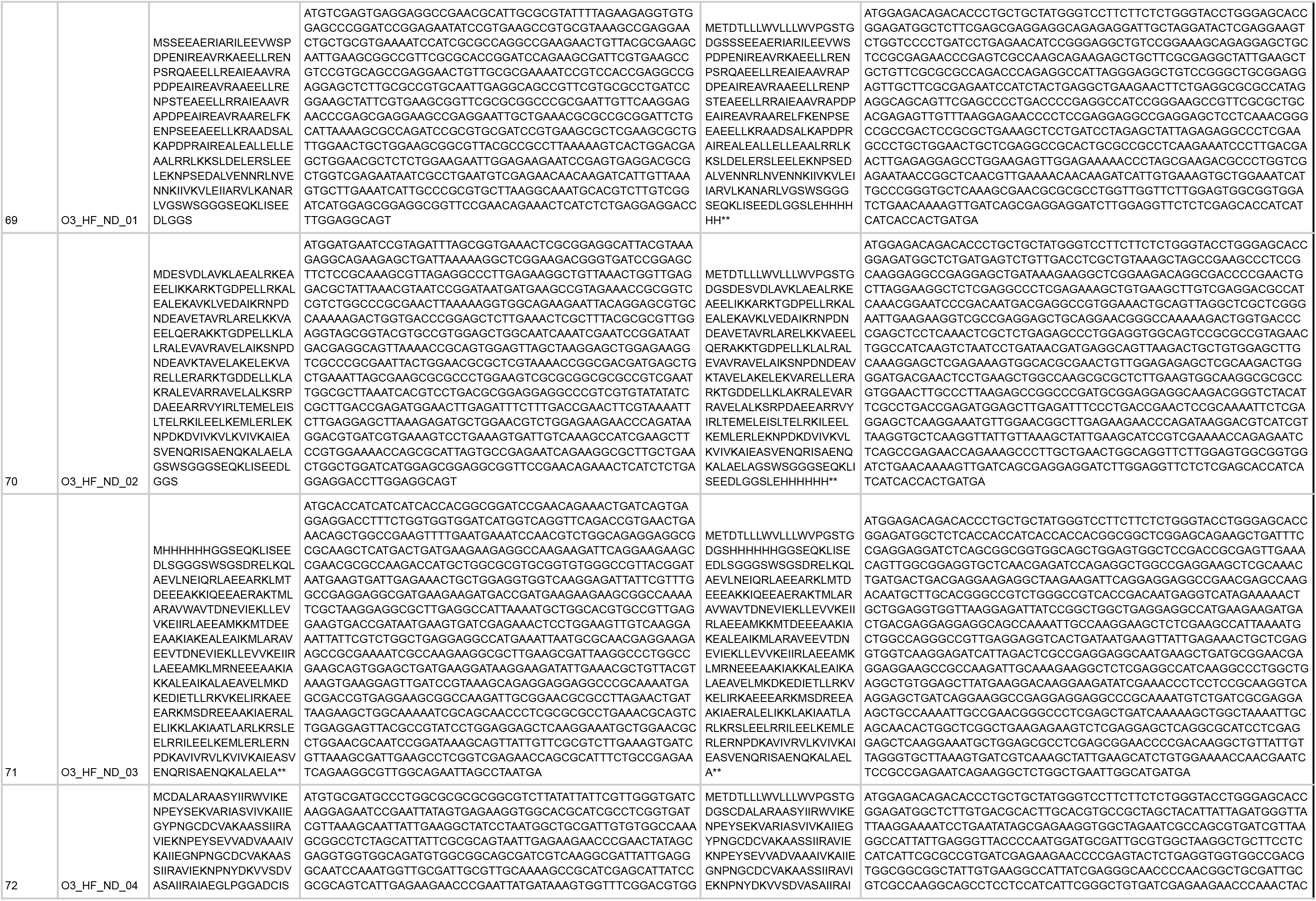
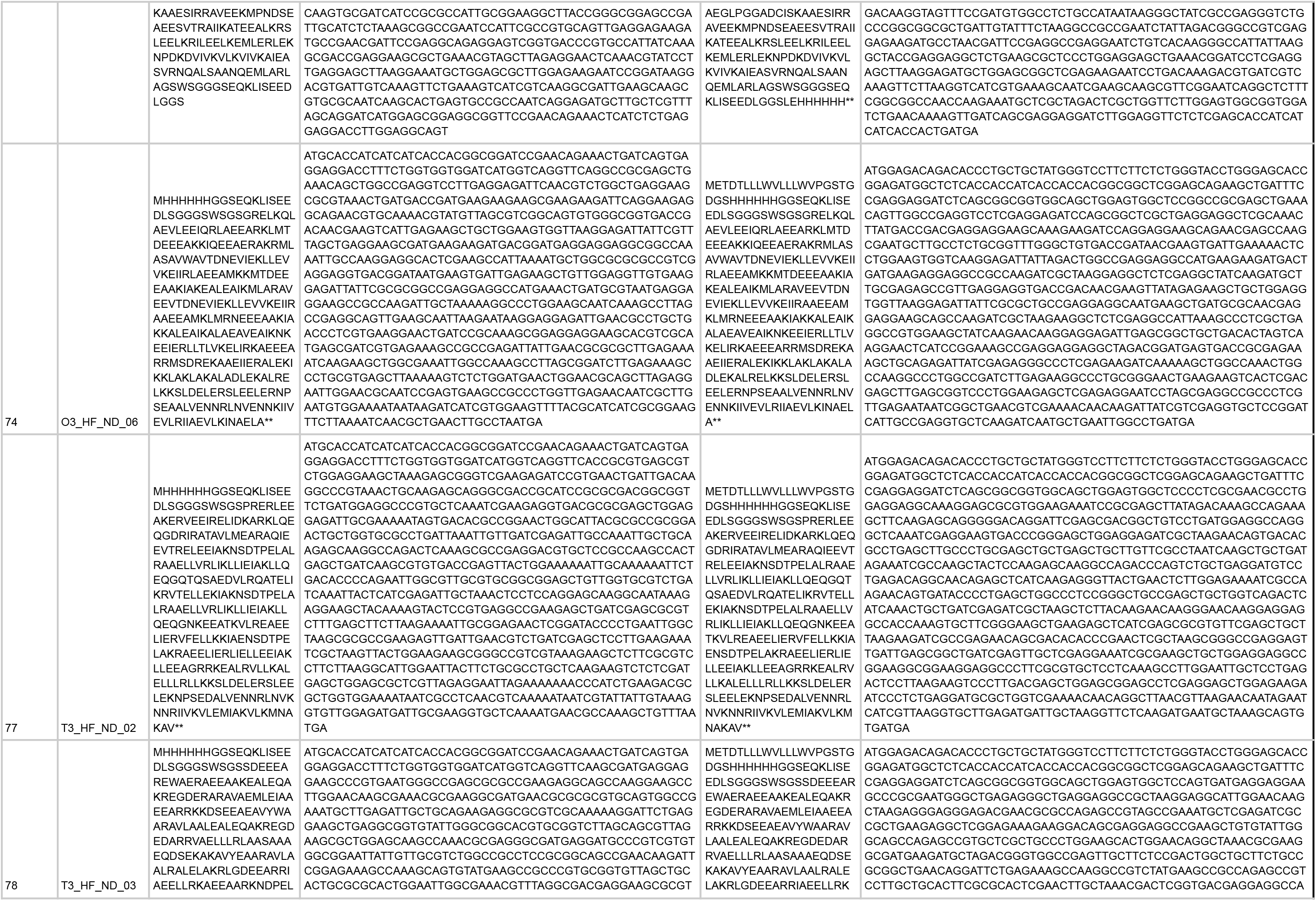
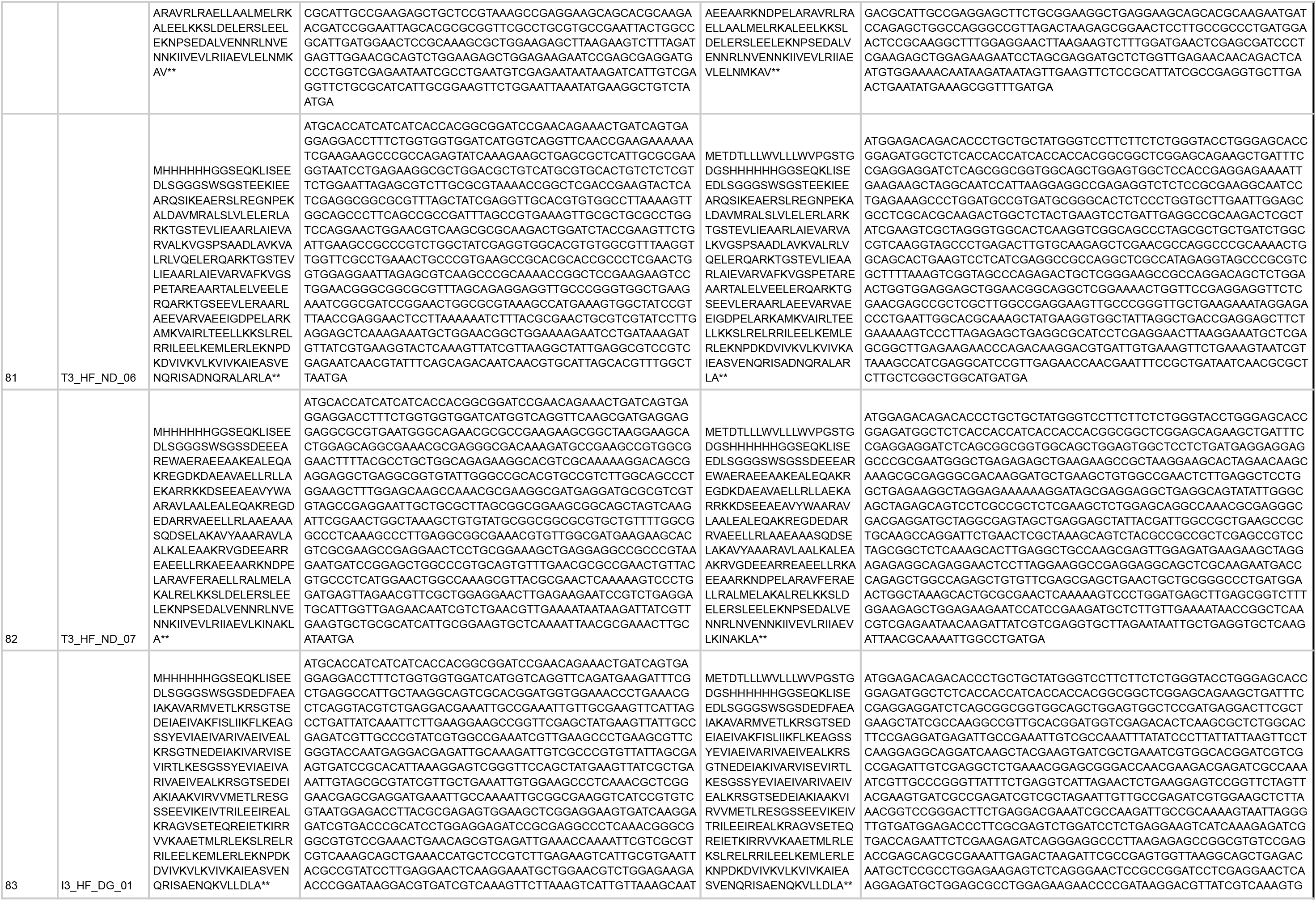
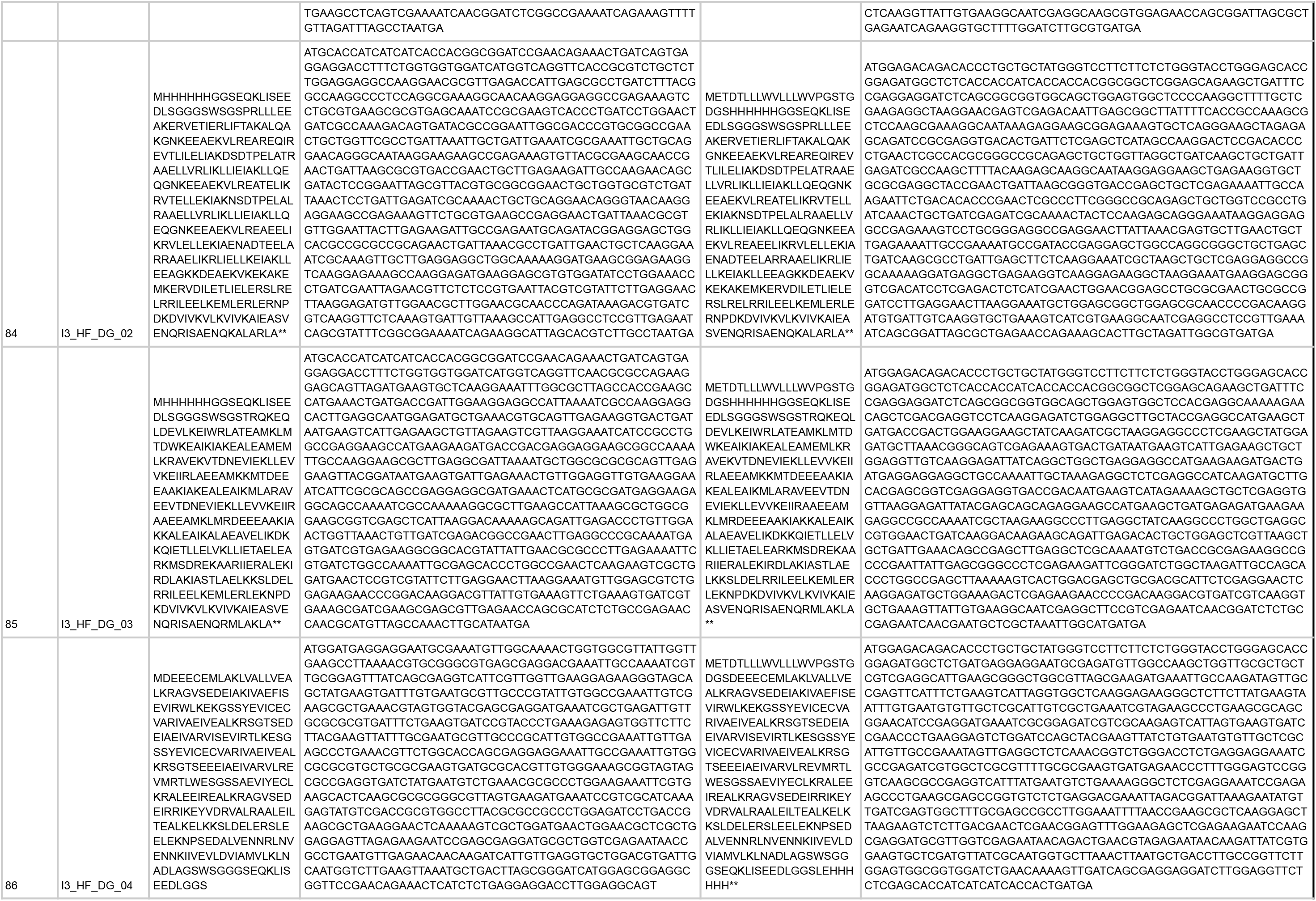
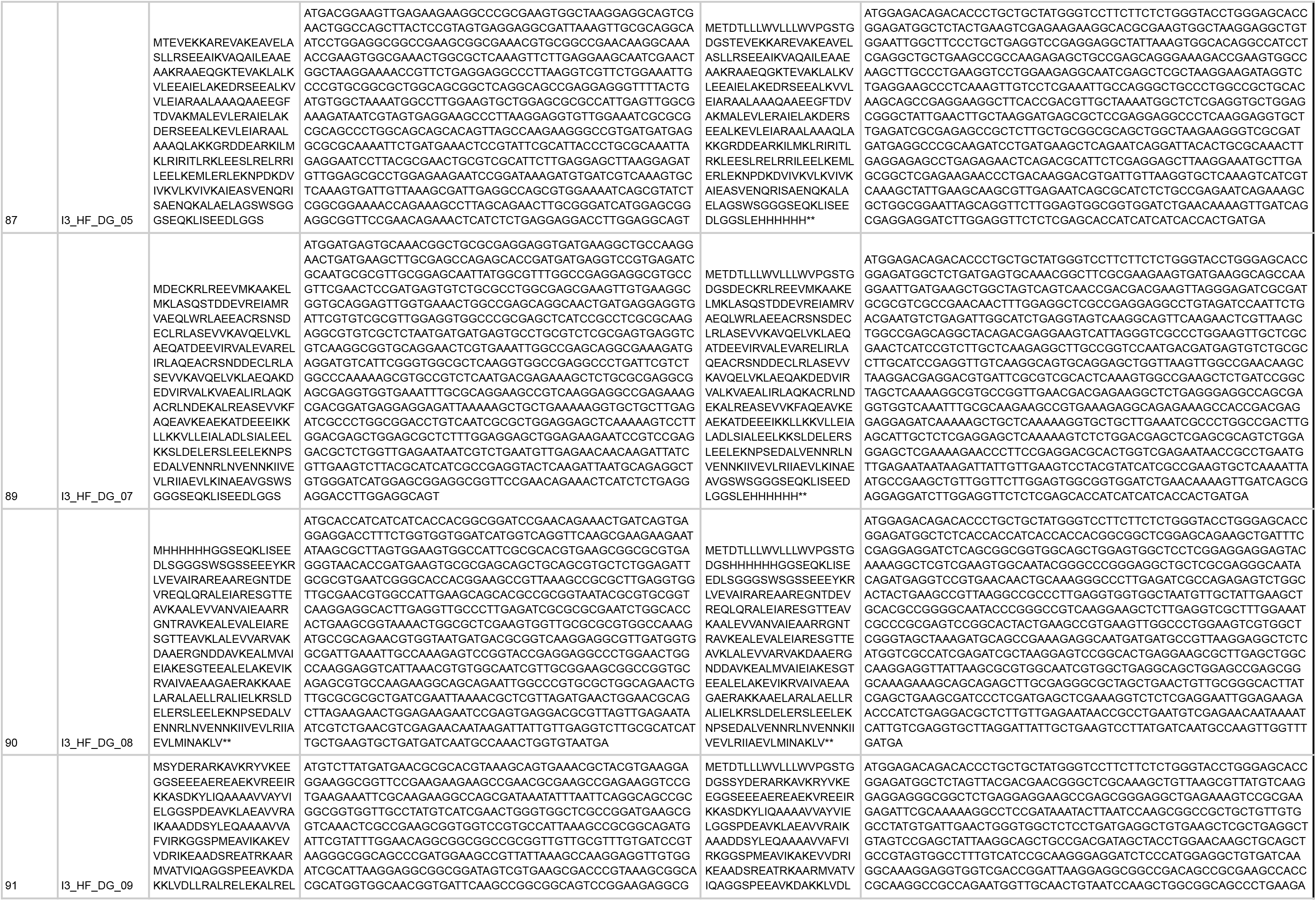
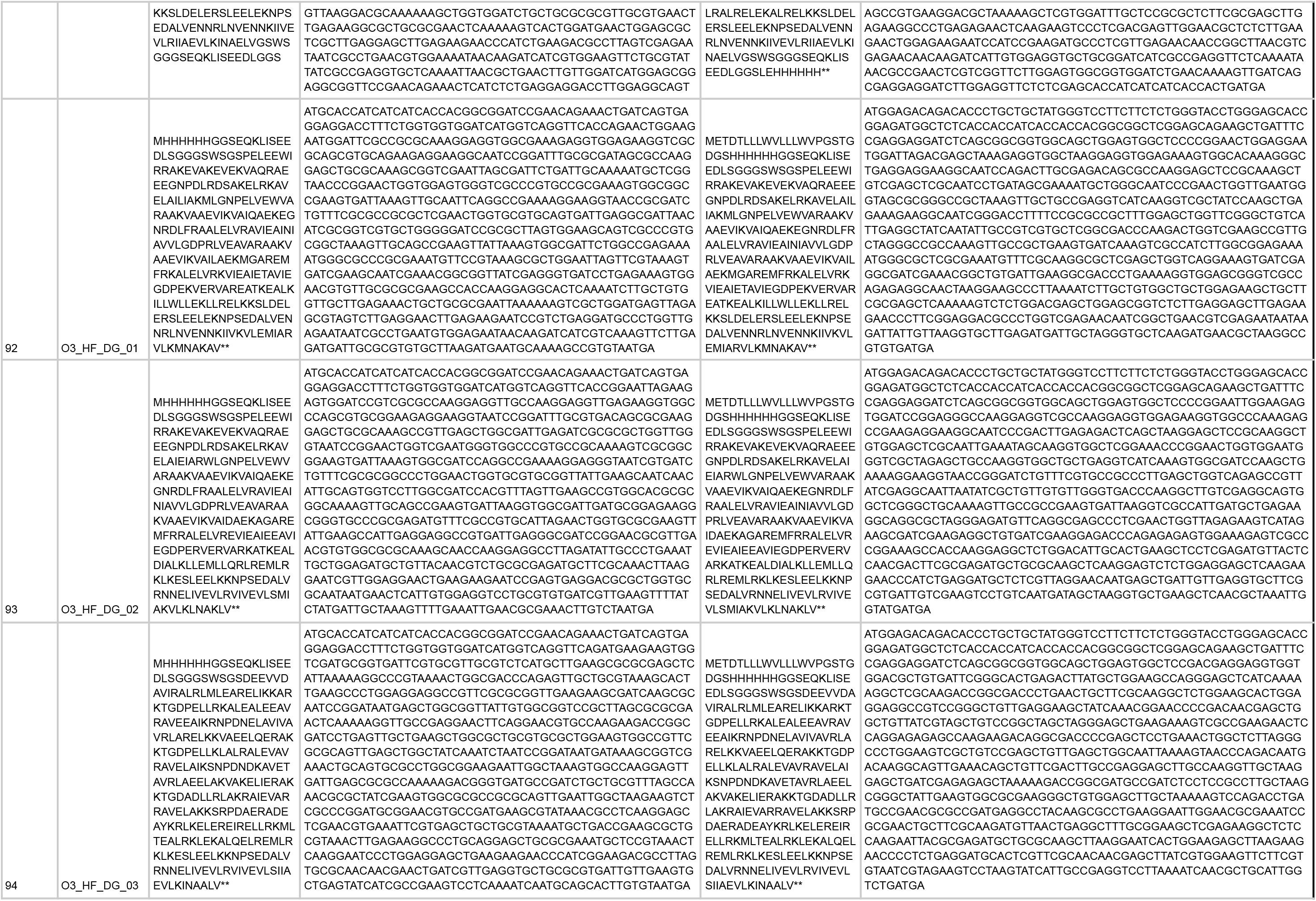
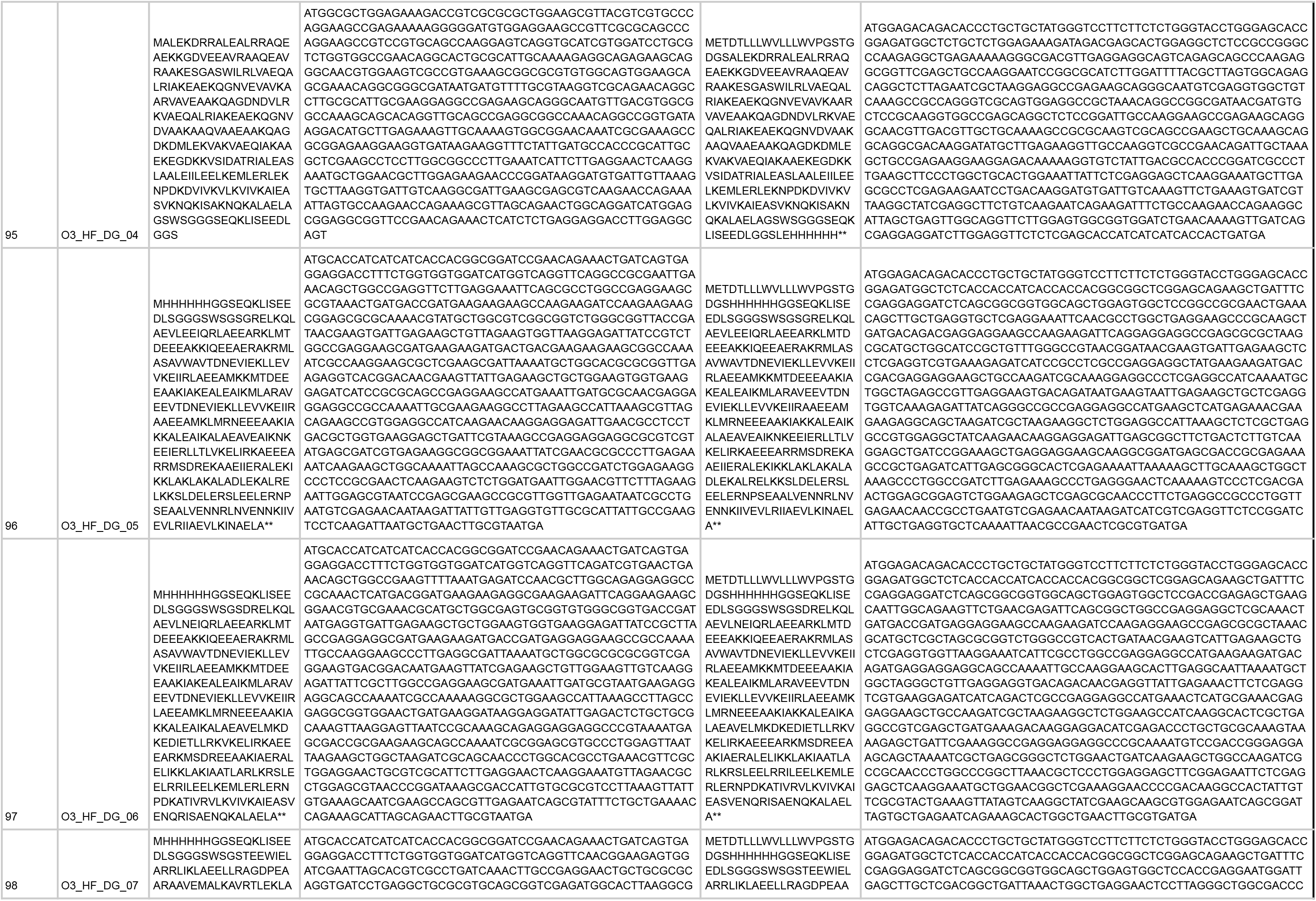
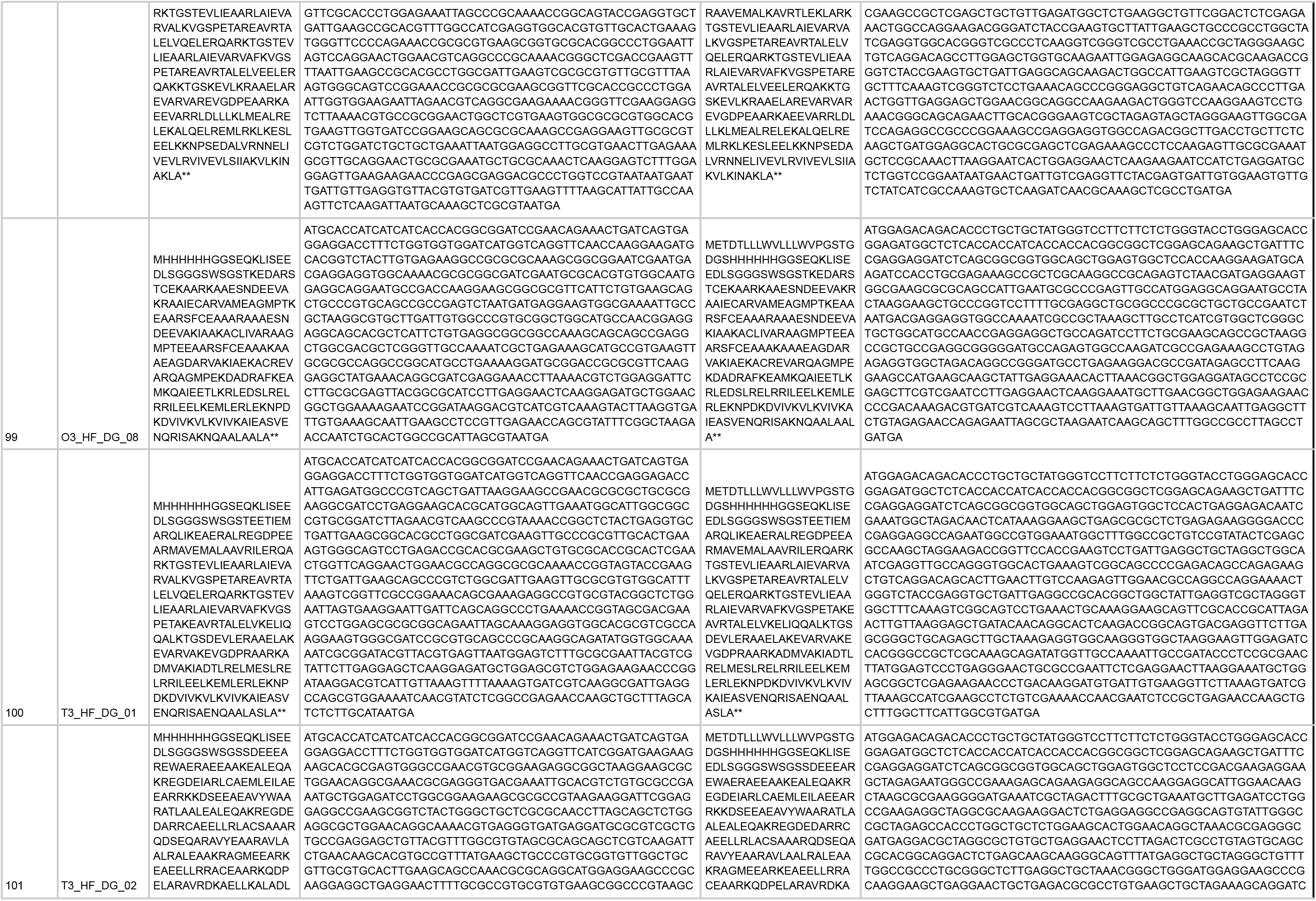
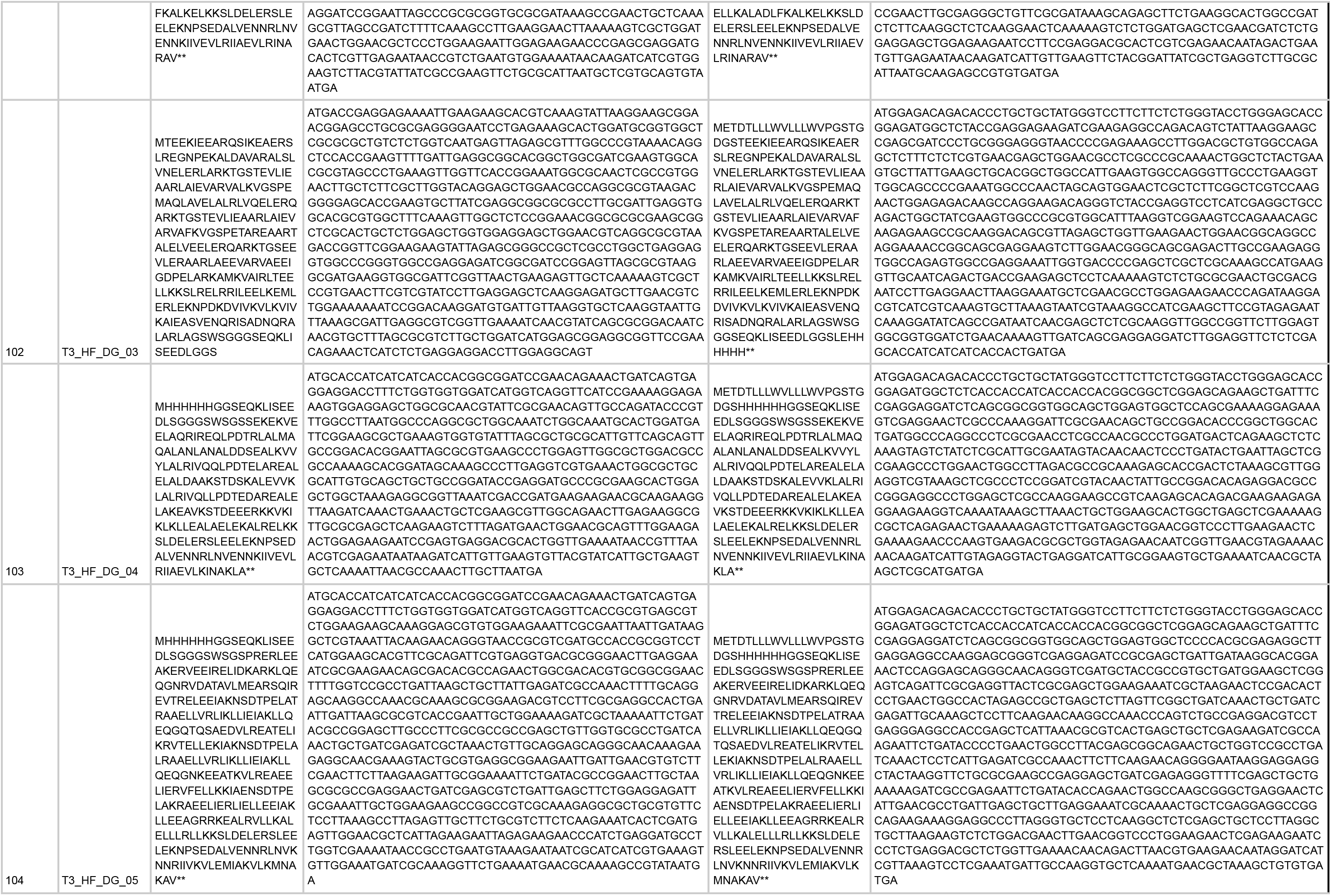
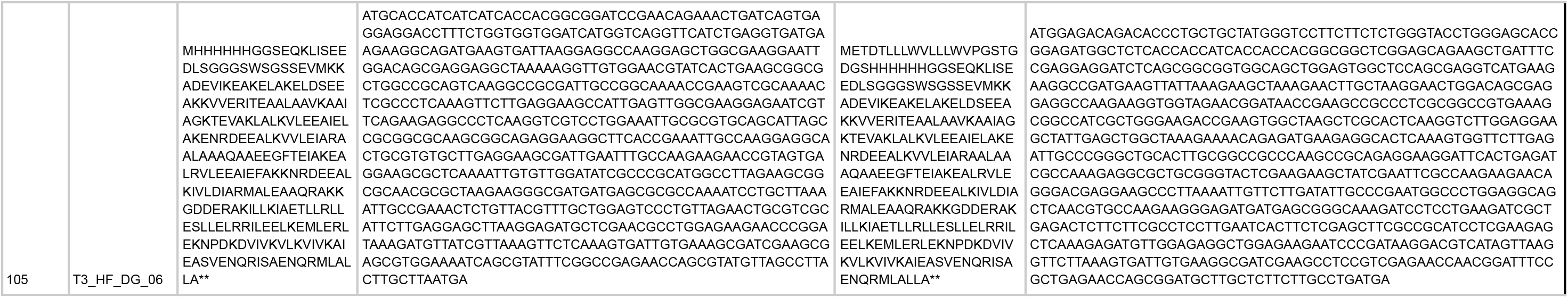
Full protein and DNA sequences for mammalian and bacterial protein expression of KWOCAs using three different protocol variations: canonical design (OG), canonical design followed by filtering based on dG_ins,pred_ (ND), and with inline degreaser (DG).

**Supplementary Table S3.** Crystallographic data collection and refinement statistics.

|  | KWOCA_39 | KWOCA_60 | KWOCA_65 | KWOCA_73 | KWOCA_102 |
| --- | --- | --- | --- | --- | --- |
| **Data collection** |  |  |  |  |  |
| Space group | H 3 | H 3 | P 63 2 2 | P 2<sub>1</sub> | P 4<sub>3</sub> 2<sub>1</sub> 2 |
| Cell dimensions |  |  |  |  |  |
| *a*, *b*, *c* (Å) | 115.66,<br>115.66,<br>72.08 | 90.53, 90.53,<br>125.52 | 70.97, 70.97,<br>200.56 | 72.00,<br>170.81,<br>72.15 | 89.40, 89.40,<br>273.03 |
| $\alpha$, $\beta$, $\gamma$ (°) | 90, 90, 120 | 90, 90, 120 | 90, 90, 120 | 90, 117.83,<br>90 | 90, 90, 90 |
| Resolution (Å) | 58.51 - 3.61<br>(3.96 - 3.61)* | 48.99 - 3.25<br>(3.51 - 3.25)* | 61.27 - 3.15<br>(3.37 - 3.15)* | 42.70 - 3.04<br>(3.11 - 3.04)* | 50.0 - 3.80<br>(3.87 - 3.8)* |
| *R*<sub>merge</sub> | 0.212 (0.663) | 0.119 (261) | 0.266 (2.286) | 0.151 (1.631) | 0.086 (0.803) |
| *I* / $\sigma$ *I* | 4.2 (1.8) | 10.2 (5.5) | 6.0 (1.1) | 4.8 (0.8) | 22.6 (2.7) |
| Completeness (%) | 100.0 (100.0) | 100.0 (100.0) | 100.0 (100.0) | 99.2 (99.9) | 100.0 (100.0) |
| Unique reflections | 4127 (983) | 52176<br>(11095) | 65007 (12073) | 29424 (4770) | 11536 (2668) |
| Redundancy | 5.4 (5.6) | 8.6 (8.6) | 11.3 (12.1) | 3.7 (3.8) | 16.5 (15.6) |
| *CC*1/2 | 0.978 (0.665) | 0.998 (0.975) | 0.995 (0.648) | 0.988 (0.293) | 0.998 (0.981) |
| **Refinement** |  |  |  |  |  |
| Resolution (Å) | 58.51 - 3.61<br>(3.97 - 3.61) | 41.63 - 3.25<br>(4.09 - 3.25) | 61.27 - 3.15<br>(3.47 - 3.15)* | 42.70 - 3.04<br>(3.11 - 3.04) | 46.42 - 3.76<br>(3.85 - 3.76) |
| No. reflections | 4126 (1031) | 5938 (2981) | 5660 (1360) | 29378 (2075) | 10455 (1275) |
| *R*<sub>work</sub> / *R*<sub>free</sub> | 0.23/0.27<br>(0.36/0.38) | 0.24/0.28<br>(0.24/0.30) | 0.27/0.30<br>(0.31/0.38) | 0.26/0.30<br>(0.39/0.43) | 0.25/0.30<br>(0.32/0.40) |
| No. atoms | 2141 | 2386 | 1710 | 10153 | 5952 |
| Protein | 2141 | 2386 | 1710 | 10153 | 5952 |
| *B*-factors (Å<sup>2</sup>) | 183 | 29 | 96 | 109 | 78 |
| Protein | 183 | 29 | 96 | 109 | 78 |
| Bond lengths (Å) | 0.001 | 0.002 | 0.002 | 0.002 | 0.008 |
| Bond angles (°) | 0.326 | 1.336 | 0.347 | 0.371 | 1.309 |

\*Data collected from a single crystal. \*Values in parentheses are for the highest-resolution shell.

**Supplementary Table S4.** CryoEM data collection information.

|  | KWOCA_4 | KWOCA_51 |
| --- | --- | --- |
| Microscope | Talos Arctica | FEI Titan Krios |
| Voltage (kV) | 200 | 300 |
| Detector | Gatan K2 Summit | Gatan K3 Summit |
| Recording mode | Counting | Counting |
| Magnification | 36, 000 X | 105,000 X |
| Movie micrograph pixel size (Å) | 1.15 | 0.84 |
| Dose rate (e<sup>-</sup>/Å<sup>2</sup>/s) | 5.11 | 21.25 |
| No. of frames per movie micrograph | 49 | 75 |
| Frame exposure time (ms) | 200 | 40 |
| Movie micrograph exposure time (s) | 9.8 | 3 |
| Total dose (e<sup>-</sup>/Å<sup>2</sup>) | 50.1 | 65.47 |
| Under focus range (µm) | 1.0 - 1.5 | 0.8 - 1.2 |
| Number of movie micrographs | 1341 | 5932 |

## References

Abrescia, Nicola G. A., Dennis H. Bamford, Jonathan M. Grimes, and David I. Stuart. 2012. “Structure Unifies the Viral Universe.” Annual Review of Biochemistry 81 (April): 795–822.

Adams, Paul D., Pavel V. Afonine, Gábor Bunkóczi, Vincent B. Chen, Ian W. Davis, Nathaniel Echols, Jeffrey J. Headd, et al. 2010. “PHENIX: A Comprehensive Python-Based System for Macromolecular Structure Solution.” *Acta Crystallographica. Section D*, Biological Crystallography 66 (Pt 2): 213–21.

Akopian, David, Kuang Shen, Xin Zhang, and Shu-Ou Shan. 2013. “Signal Recognition Particle: An Essential Protein-Targeting Machine.” Annual Review of Biochemistry 82 (February): 693–721.

Baek, Minkyung, and David Baker. 2022. “Deep Learning and Protein Structure Modeling.” Nature Methods 19 (1): 13–14.

Bale, Jacob B., Shane Gonen, Yuxi Liu, William Sheffler, Daniel Ellis, Chantz Thomas, Duilio Cascio, et al. 2016. “Accurate Design of Megadalton-Scale Two-Component Icosahedral Protein Complexes.” Science 353 (6297): 389–94.

Barouch, Dan H., Zhi-Yong Yang, Wing-Pui Kong, Birgit Korioth-Schmitz, Shawn M. Sumida, Diana M. Truitt, Michael G. Kishko, et al. 2005. “A Human T-Cell Leukemia Virus Type 1 Regulatory Element Enhances the Immunogenicity of Human Immunodeficiency Virus Type 1 DNA Vaccines in Mice and Nonhuman Primates.” Journal of Virology 79 (14): 8828–34.

Baycin-Hizal, Deniz, David L. Tabb, Raghothama Chaerkady, Lily Chen, Nathan E. Lewis, Harish Nagarajan, Vishaldeep Sarkaria, et al. 2012. “Proteomic Analysis of Chinese Hamster Ovary Cells.” Journal of Proteome Research 11 (11): 5265–76.

Bobik, Thomas A., Brent P. Lehman, and Todd O. Yeates. 2015. “Bacterial Microcompartments: Widespread Prokaryotic Organelles for Isolation and Optimization of Metabolic Pathways.” Molecular Microbiology 98 (2): 193–207.

Boder, E. T., and K. D. Wittrup. 1997. “Yeast Surface Display for Screening Combinatorial Polypeptide Libraries.” Nature Biotechnology 15 (6): 553–57.

Boone, Morgane, Pathmanaban Ramasamy, Jasper Zuallaert, Robbin Bouwmeester, Berre Van Moer, Davy Maddelein, Demet Turan, et al. 2021. “Massively Parallel Interrogation of Protein Fragment Secretability Using SECRiFY Reveals Features Influencing Secretory System Transit.” Nature Communications 12 (1): 6414.

Boyken, Scott E., Mark A. Benhaim, Florian Busch, Mengxuan Jia, Matthew J. Bick, Heejun Choi, Jason C. Klima, et al. 2019. “De Novo Design of Tunable, pH-Driven Conformational Changes.” Science 364 (6441): 658–64.

Boyoglu-Barnum, Seyhan, Daniel Ellis, Rebecca A. Gillespie, Geoffrey B. Hutchinson, Young-Jun Park, Syed M. Moin, Oliver J. Acton, et al. 2021. “Quadrivalent Influenza Nanoparticle Vaccines Induce Broad Protection.” Nature 592 (7855): 623–28.

Braun, Elisabeth, and Daniel Sauter. 2019. “Furin-Mediated Protein Processing in Infectious Diseases and Cancer.” Clinical & Translational Immunology 8 (8): e1073.

Brunette, T. J., Fabio Parmeggiani, Po-Ssu Huang, Gira Bhabha, Damian C. Ekiert, Susan E. Tsutakawa, Greg L. Hura, John A. Tainer, and David Baker. 2015. “Exploring the Repeat Protein Universe through Computational Protein Design.” Nature 528 (7583): 580–84.

Bruun, Theodora U. J., Anne-Marie C. Andersson, Simon J. Draper, and Mark Howarth. 2018. “Engineering a Rugged Nanoscaffold To Enhance Plug-and-Display Vaccination.” ACS Nano 12 (9): 8855–66.

Burley, Stephen K., Charmi Bhikadiya, Chunxiao Bi, Sebastian Bittrich, Li Chen, Gregg V. Crichlow, Cole H. Christie, et al. 2021. “RCSB Protein Data Bank: Powerful New Tools for Exploring 3D Structures of Biological Macromolecules for Basic and Applied Research and Education in Fundamental Biology, Biomedicine, Biotechnology, Bioengineering and Energy Sciences.” Nucleic Acids Research 49 (D1): D437–51.

Campeotto, Ivan, Adi Goldenzweig, Jack Davey, Lea Barfod, Jennifer M. Marshall, Sarah E. Silk, Katherine E. Wright, Simon J. Draper, Matthew K. Higgins, and Sarel J. Fleishman. 2017. “One-Step Design of a Stable Variant of the Malaria Invasion Protein RH5 for Use as a Vaccine Immunogen.” Proceedings of the National Academy of Sciences of the United States of America 114 (5): 998–1002.

Carragher, B., N. Kisseberth, D. Kriegman, R. A. Milligan, C. S. Potter, J. Pulokas, and A. Reilein. 2000. “Leginon: An Automated System for Acquisition of Images from Vitreous Ice Specimens.” Journal of Structural Biology 132 (1): 33–45.

Chaudhary, Namit, Drew Weissman, and Kathryn A. Whitehead. 2021. “mRNA Vaccines for Infectious Diseases: Principles, Delivery and Clinical Translation.” Nature Reviews. Drug Discovery 20 (11): 817–38.

Classen, Scott, Greg L. Hura, James M. Holton, Robert P. Rambo, Ivan Rodic, Patrick J. McGuire, Kevin Dyer, et al. 2013. “Implementation and Performance of SIBYLS: A Dual Endstation Small-Angle X-Ray Scattering and Macromolecular Crystallography Beamline at the Advanced Light Source.” Journal of Applied Crystallography 46 (Pt 1): 1–13.

Cohen, Alexander A., Priyanthi N. P. Gnanapragasam, Yu E. Lee, Pauline R. Hoffman, Susan Ou, Leesa M. Kakutani, Jennifer R. Keeffe, et al. 2021. “Mosaic Nanoparticles Elicit Cross-Reactive Immune Responses to Zoonotic Coronaviruses in Mice.” Science 371 (6530): 735–41.

Dalvie, Neil C., Joseph R. Brady, Laura E. Crowell, Mary Kate Tracey, Andrew M. Biedermann, Kawaljit Kaur, John M. Hickey, et al. 2021. “Molecular Engineering Improves Antigen Quality and Enables Integrated Manufacturing of a Trivalent Subunit Vaccine Candidate for Rotavirus.” Microbial Cell Factories 20 (1): 94.

Dalvie, Neil C., Sergio A. Rodriguez-Aponte, Brittany L. Hartwell, Lisa H. Tostanoski, Andrew M. Biedermann, Laura E. Crowell, Kawaljit Kaur, et al. 2021. “Engineered SARS-CoV-2 Receptor Binding Domain Improves Manufacturability in Yeast and Immunogenicity in Mice.” Proceedings of the National Academy of Sciences of the United States of America 118 (38). https://doi.org/10.1073/pnas.2106845118.

DiMaio, Frank, Andrew Leaver-Fay, Phil Bradley, David Baker, and Ingemar André. 2011. “Modeling Symmetric Macromolecular Structures in Rosetta3.” PloS One 6 (6): e20450.

DiMaio, Frank, Yifan Song, Xueming Li, Matthias J. Brunner, Chunfu Xu, Vincent Conticello, Edward Egelman, Thomas Marlovits, Yifan Cheng, and David Baker. 2015. “Atomic-Accuracy Models from 4.5-Å Cryo-Electron Microscopy Data with Density-Guided Iterative Local Refinement.” Nature Methods 12 (4): 361–65.

Douglas, Trevor, and Mark Young. 2006. “Viruses: Making Friends with Old Foes.” Science 312 (5775): 873–75.

Dou, Jiayi, Anastassia A. Vorobieva, William Sheffler, Lindsey A. Doyle, Hahnbeom Park, Matthew J. Bick, Binchen Mao, et al. 2018. “De Novo Design of a Fluorescence-Activating β-Barrel.” Nature 561 (7724): 485–91.

Dyer, Kevin N., Michal Hammel, Robert P. Rambo, Susan E. Tsutakawa, Ivan Rodic, Scott Classen, John A. Tainer, and Greg L. Hura. 2014. “High-Throughput SAXS for the Characterization of Biomolecules in Solution: A Practical Approach.” Methods in Molecular Biology 1091: 245–58.

Edwardson, Thomas G. W., Stephan Tetter, and Donald Hilvert. 2020. “Two-Tier Supramolecular Encapsulation of Small Molecules in a Protein Cage.” Nature Communications 11 (1): 5410.

Egea, Pascal F., Robert M. Stroud, and Peter Walter. 2005. “Targeting Proteins to Membranes: Structure of the Signal Recognition Particle.” Current Opinion in Structural Biology 15 (2): 213–20.

Ellis, Daniel, Natalie Brunette, Katharine H. D. Crawford, Alexandra C. Walls, Minh N. Pham, Chengbo Chen, Karla-Luise Herpoldt, et al. 2021. “Stabilization of the SARS-CoV-2 Spike Receptor-Binding Domain Using Deep Mutational Scanning and Structure-Based Design.” Frontiers in Immunology 12 (June): 710263.

Emsley, Paul, and Kevin Cowtan. 2004. “Coot: Model-Building Tools for Molecular Graphics.” Acta Crystallographica. Section D, Biological Crystallography 60 (Pt 12 Pt 1): 2126–32.

Fallas, Jorge A., George Ueda, William Sheffler, Vanessa Nguyen, Dan E. McNamara, Banumathi Sankaran, Jose Henrique Pereira, et al. 2017. “Computational Design of Self-Assembling Cyclic Protein Homo-Oligomers.” Nature Chemistry 9 (4): 353–60.

Farhan, Hesso, and Catherine Rabouille. 2011. “Signalling to and from the Secretory Pathway.” Journal of Cell Science 124 (Pt 2): 171–80.

Goodsell, D. S., and A. J. Olson. 2000. “Structural Symmetry and Protein Function.” Annual Review of Biophysics and Biomolecular Structure 29: 105–53.

Gordon, Sydney R., Elizabeth J. Stanley, Sarah Wolf, Angus Toland, Sean J. Wu, Daniel Hadidi, Jeremy H. Mills, David Baker, Ingrid Swanson Pultz, and Justin B. Siegel. 2012. “Computational Design of an α-Gliadin Peptidase.” Journal of the American Chemical Society 134 (50): 20513–20.

Güler-Gane, Gülin, Sara Kidd, Sudharsan Sridharan, Tristan J. Vaughan, Trevor C. I. Wilkinson, and Natalie J. Tigue. 2016. “Overcoming the Refractory Expression of Secreted Recombinant Proteins in Mammalian Cells through Modification of the Signal Peptide and Adjacent Amino Acids.” PloS One 11 (5): e0155340.

Heijne, G. von. 1990. “The Signal Peptide.” The Journal of Membrane Biology 115 (3): 195–201.

He, Linling, Xiaohe Lin, Ying Wang, Ciril Abraham, Cindy Sou, Timothy Ngo, Yi Zhang, Ian A. Wilson, and Jiang Zhu. 2021. “Single-Component, Self-Assembling, Protein Nanoparticles Presenting the Receptor Binding Domain and Stabilized Spike as SARS-CoV-2 Vaccine Candidates.” Science Advances 7 (12). https://doi.org/10.1126/sciadv.abf1591.

Hessa, Tara, Hyun Kim, Karl Bihlmaier, Carolina Lundin, Jorrit Boekel, Helena Andersson, Ingmarie Nilsson, Stephen H. White, and Gunnar von Heijne. 2005. “Recognition of Transmembrane Helices by the Endoplasmic Reticulum Translocon.” Nature 433 (7024): 377–81.

Hessa, Tara, Nadja M. Meindl-Beinker, Andreas Bernsel, Hyun Kim, Yoko Sato, Mirjam Lerch-Bader, Ingmarie Nilsson, Stephen H. White, and Gunnar von Heijne. 2007. “Molecular Code for Transmembrane-Helix Recognition by the Sec61 Translocon.” Nature 450 (7172): 1026–30.

Hsia, Yang, Jacob B. Bale, Shane Gonen, Dan Shi, William Sheffler, Kimberly K. Fong, Una Nattermann, et al. 2016. “Design of a Hyperstable 60-Subunit Protein Dodecahedron. [corrected].” Nature 535 (7610): 136–39.

Hsia, Yang, Rubul Mout, William Sheffler, Natasha I. Edman, Ivan Vulovic, Young-Jun Park, Rachel L. Redler, et al. 2021. “Design of Multi-Scale Protein Complexes by Hierarchical Building Block Fusion.” Nature Communications 12 (1): 2294.

Hsieh, Ching-Lin, Jory A. Goldsmith, Jeffrey M. Schaub, Andrea M. DiVenere, Hung-Che Kuo, Kamyab Javanmardi, Kevin C. Le, et al. 2020. “Structure-Based Design of Prefusion-Stabilized SARS-CoV-2 Spikes.” Science 369 (6510): 1501–5.

Huang, Po-Ssu, Scott E. Boyken, and David Baker. 2016. “The Coming of Age of de Novo Protein Design.” Nature 537 (7620): 320–27.

Hura, Greg L., Angeli L. Menon, Michal Hammel, Robert P. Rambo, Farris L. Poole 2nd, Susan E. Tsutakawa, Francis E. Jenney Jr, et al. 2009. “Robust, High-Throughput Solution Structural Analyses by Small Angle X-Ray Scattering (SAXS).” Nature Methods 6 (8): 606–12.

Janin, Joël, Ranjit P. Bahadur, and Pinak Chakrabarti. 2008. “Protein-Protein Interaction and Quaternary Structure.” Quarterly Reviews of Biophysics 41 (2): 133–80.

Jardine, Joseph G., Takayuki Ota, Devin Sok, Matthias Pauthner, Daniel W. Kulp, Oleksandr Kalyuzhniy, Patrick D. Skog, et al. 2015. “HIV-1 VACCINES. Priming a Broadly Neutralizing Antibody Response to HIV-1 Using a Germline-Targeting Immunogen.” Science 349 (6244): 156–61.

Jardine, Joseph, Jean-Philippe Julien, Sergey Menis, Takayuki Ota, Oleksandr Kalyuzhniy, Andrew McGuire, Devin Sok, et al. 2013. “Rational HIV Immunogen Design to Target Specific Germline B Cell Receptors.” Science 340 (6133): 711–16.

Kabsch, Wolfgang. 2010. “XDS.” Acta Crystallographica. Section D, Biological Crystallography 66 (Pt 2): 125–32.

Kanekiyo, Masaru, Chih-Jen Wei, Hadi M. Yassine, Patrick M. McTamney, Jeffrey C. Boyington, James R. R. Whittle, Srinivas S. Rao, Wing-Pui Kong, Lingshu Wang, and Gary J. Nabel. 2013. “Self-Assembling Influenza Nanoparticle Vaccines Elicit Broadly Neutralizing H1N1 Antibodies.” Nature 499 (7456): 102–6.

Kelley, Brian. 2009. “Industrialization of mAb Production Technology: The Bioprocessing Industry at a Crossroads.” mAbs 1 (5): 443–52.

Khmelinskaia, Alena, Adam Wargacki, and Neil P. King. 2021. “Structure-Based Design of Novel Polyhedral Protein Nanomaterials.” Current Opinion in Microbiology 61 (June): 51–57.

King, Neil P., Jacob B. Bale, William Sheffler, Dan E. McNamara, Shane Gonen, Tamir Gonen, Todd O. Yeates, and David Baker. 2014. “Accurate Design of Co-Assembling Multi-Component Protein Nanomaterials.” Nature 510 (7503): 103–8.

King, Neil P., William Sheffler, Michael R. Sawaya, Breanna S. Vollmar, John P. Sumida, Ingemar André, Tamir Gonen, Todd O. Yeates, and David Baker. 2012. “Computational Design of Self-Assembling Protein Nanomaterials with Atomic Level Accuracy.” Science 336 (6085): 1171–74.

Klima, Jason C., Lindsey A. Doyle, Justin Daho Lee, Michael Rappleye, Lauren A. Gagnon, Min Yen Lee, Emilia P. Barros, et al. 2021. “Incorporation of Sensing Modalities into de Novo Designed Fluorescence-Activating Proteins.” Nature Communications 12 (1): 856.

Kober, Lars, Christoph Zehe, and Juergen Bode. 2013. “Optimized Signal Peptides for the Development of High Expressing CHO Cell Lines.” Biotechnology and Bioengineering 110 (4): 1164–73.

Konrath, Kylie M., Kevin Liaw, Yuanhan Wu, Xizhou Zhu, Susanne N. Walker, Ziyang Xu, Katherine Schultheis, et al. 2022. “Nucleic Acid Delivery of Immune-Focused SARS-CoV-2 Nanoparticles Drives Rapid and Potent Immunogenicity Capable of Single-Dose Protection.” Cell Reports, January, 110318.

Kuo, Chih-Chung, Austin Wt Chiang, Isaac Shamie, Mojtaba Samoudi, Jahir M. Gutierrez, and Nathan E. Lewis. 2018. “The Emerging Role of Systems Biology for Engineering Protein Production in CHO Cells.” Current Opinion in Biotechnology 51 (June): 64–69.

Lawrence, M. C., and P. M. Colman. 1993. “Shape Complementarity at Protein/protein Interfaces.” Journal of Molecular Biology 234 (4): 946–50.

Leaver-Fay, Andrew, Michael Tyka, Steven M. Lewis, Oliver F. Lange, James Thompson, Ron Jacak, Kristian Kaufman, et al. 2011. “ROSETTA3: An Object-Oriented Software Suite for the Simulation and Design of Macromolecules.” Methods in Enzymology 487: 545–74.

Le Fourn, Valérie, Pierre-Alain Girod, Montse Buceta, Alexandre Regamey, and Nicolas Mermod. 2014. “CHO Cell Engineering to Prevent Polypeptide Aggregation and Improve Therapeutic Protein Secretion.” Metabolic Engineering 21 (January): 91–102.

Leman, Julia Koehler, Brian D. Weitzner, Steven M. Lewis, Jared Adolf-Bryfogle, Nawsad Alam, Rebecca F. Alford, Melanie Aprahamian, et al. 2020. “Macromolecular Modeling and Design in Rosetta: Recent Methods and Frameworks.” Nature Methods 17 (7): 665–80.

Liu, Yan, Anton Nguyen, Robert L. Wolfert, and Shaoqiu Zhuo. 2009. “Enhancing the Secretion of Recombinant Proteins by Engineering N-Glycosylation Sites.” Biotechnology Progress 25 (5): 1468–75.

Li, Xueming, Paul Mooney, Shawn Zheng, Christopher R. Booth, Michael B. Braunfeld, Sander Gubbens, David A. Agard, and Yifan Cheng. 2013. “Electron Counting and Beam-Induced Motion Correction Enable near-Atomic-Resolution Single-Particle Cryo-EM.” Nature Methods 10 (6): 584–90.

López-Sagaseta, Jacinto, Enrico Malito, Rino Rappuoli, and Matthew J. Bottomley. 2016. “Self-Assembling Protein Nanoparticles in the Design of Vaccines.” Computational and Structural Biotechnology Journal 14: 58–68.

Malladi, Sameer Kumar, David Schreiber, Ishika Pramanick, Malavika Abhineshababu Sridevi, Adi Goldenzweig, Somnath Dutta, Sarel Jacob Fleishman, and Raghavan Varadarajan. 2020. “One-Step Sequence and Structure-Guided Optimization of HIV-1 Envelope gp140.” Current Research in Structural Biology 2 (April): 45–55.

Marcandalli, Jessica, Brooke Fiala, Sebastian Ols, Michela Perotti, Willem de van der Schueren, Joost Snijder, Edgar Hodge, et al. 2019. “Induction of Potent Neutralizing Antibody Responses by a Designed Protein Nanoparticle Vaccine for Respiratory Syncytial Virus.” Cell 176 (6): 1420–31.e17.

McCoy, Airlie J., Ralf W. Grosse-Kunstleve, Paul D. Adams, Martyn D. Winn, Laurent C. Storoni, and Randy J. Read. 2007. “Phaser Crystallographic Software.” Journal of Applied Crystallography 40 (Pt 4): 658–74.

Mu, Zekun, Kevin Wiehe, Kevin O. Saunders, Rory Henderson, Derek W. Cain, Robert Parks, Diana Martik, et al. 2021. “Ability of Nucleoside-Modified mRNA to Encode HIV-1 Envelope Trimer Nanoparticles.” *bioRxiv : The Preprint Server for Biology*, August. https://doi.org/10.1101/2021.08.09.455714.

Ohtsubo, Kazuaki, and Jamey D. Marth. 2006. “Glycosylation in Cellular Mechanisms of Health and Disease.” Cell 126 (5): 855–67.

Padilla, J. E., C. Colovos, and T. O. Yeates. 2001. “Nanohedra: Using Symmetry to Design Self Assembling Protein Cages, Layers, Crystals, and Filaments.” Proceedings of the National Academy of Sciences of the United States of America 98 (5): 2217–21.

Pallesen, Jesper, Nianshuang Wang, Kizzmekia S. Corbett, Daniel Wrapp, Robert N. Kirchdoerfer, Hannah L. Turner, Christopher A. Cottrell, et al. 2017. “Immunogenicity and Structures of a Rationally Designed Prefusion MERS-CoV Spike Antigen.” Proceedings of the National Academy of Sciences of the United States of America 114 (35): E7348–57.

Palombarini, Federica, Elisa Di Fabio, Alberto Boffi, Alberto Macone, and Alessandra Bonamore. 2020. “Ferritin Nanocages for Protein Delivery to Tumor Cells.” Molecules 25 (4). https://doi.org/10.3390/molecules25040825.

Peleg, Yoav, Renaud Vincentelli, Brett M. Collins, Kai-En Chen, Emma K. Livingstone, Saroja Weeratunga, Natalya Leneva, et al. 2021. “Community-Wide Experimental Evaluation of the PROSS Stability-Design Method.” Journal of Molecular Biology 433 (13): 166964.

Pettersen, Eric F., Thomas D. Goddard, Conrad C. Huang, Gregory S. Couch, Daniel M. Greenblatt, Elaine C. Meng, and Thomas E. Ferrin. 2004. “UCSF Chimera--a Visualization System for Exploratory Research and Analysis.” Journal of Computational Chemistry 25 (13): 1605–12.

Pettersen, Eric F., Thomas D. Goddard, Conrad C. Huang, Elaine C. Meng, Gregory S. Couch, Tristan I. Croll, John H. Morris, and Thomas E. Ferrin. 2021. “UCSF ChimeraX: Structure Visualization for Researchers, Educators, and Developers.” Protein Science: A Publication of the Protein Society 30 (1): 70–82.

Punjani, Ali, John L. Rubinstein, David J. Fleet, and Marcus A. Brubaker. 2017. “cryoSPARC: Algorithms for Rapid Unsupervised Cryo-EM Structure Determination.” Nature Methods 14 (3): 290–96.

Putnam, Christopher D., Michal Hammel, Greg L. Hura, and John A. Tainer. 2007. “X-Ray Solution Scattering (SAXS) Combined with Crystallography and Computation: Defining Accurate Macromolecular Structures, Conformations and Assemblies in Solution.” Quarterly Reviews of Biophysics 40 (3): 191–285.

Rajoo, Sasikumar, Pascal Vallotton, Evgeny Onischenko, and Karsten Weis. 2018. “Stoichiometry and Compositional Plasticity of the Yeast Nuclear Pore Complex Revealed by Quantitative Fluorescence Microscopy.” Proceedings of the National Academy of Sciences of the United States of America 115 (17): E3969–77.

Rakestraw, J. Andy, Stephen L. Sazinsky, Andrea Piatesi, Eugene Antipov, and K. Dane Wittrup. 2009. “Directed Evolution of a Secretory Leader for the Improved Expression of Heterologous Proteins and Full-Length Antibodies in Saccharomyces Cerevisiae.” Biotechnology and Bioengineering 103 (6): 1192–1201.

Rapoport, Tom A. 2007. “Protein Translocation across the Eukaryotic Endoplasmic Reticulum and Bacterial Plasma Membranes.” Nature 450 (7170): 663–69.

Rawi, Reda, Lucy Rutten, Yen-Ting Lai, Adam S. Olia, Sven Blokland, Jarek Juraszek, Chen-Hsiang Shen, et al. 2020. “Automated Design by Structure-Based Stabilization and Consensus Repair to Achieve Prefusion-Closed Envelope Trimers in a Wide Variety of HIV Strains.” Cell Reports 33 (8): 108432.

Rojas, Gertrudis, Tania Carmenate, Julio Felipe Santo-Tomás, Pedro A. Valiente, Marlies Becker, Annia Pérez-Riverón, Yaima Tundidor, et al. 2019. “Directed Evolution of Super-Secreted Variants from Phage-Displayed Human Interleukin-2.” Scientific Reports 9 (1): 800.

Ruiter, M. V. de, R. M. van der Hee, A. J. M. Driessen, E. D. Keurhorst, M. Hamid, and J. J. L. M. Cornelissen. 2019. “Polymorphic Assembly of Virus-Capsid Proteins around DNA and the Cellular Uptake of the Resulting Particles.” Journal of Controlled Release: Official Journal of the Controlled Release Society 307 (August): 342–54.

Rupp, Oliver, Madolyn L. MacDonald, Shangzhong Li, Heena Dhiman, Shawn Polson, Sven Griep, Kelley Heffner, et al. 2018. “A Reference Genome of the Chinese Hamster Based on a Hybrid Assembly Strategy.” Biotechnology and Bioengineering 115 (8): 2087–2100.

Sagt, C. M., B. Kleizen, R. Verwaal, M. D. de Jong, W. H. Müller, A. Smits, C. Visser, J. Boonstra, A. J. Verkleij, and C. T. Verrips. 2000. “Introduction of an N-Glycosylation Site Increases Secretion of Heterologous Proteins in Yeasts.” Applied and Environmental Microbiology 66 (11): 4940–44.

Sahasrabuddhe, Aniruddha, Yang Hsia, Florian Busch, William Sheffler, Neil P. King, David Baker, and Vicki H. Wysocki. 2018. “Confirmation of Intersubunit Connectivity and Topology of Designed Protein Complexes by Native MS.” Proceedings of the National Academy of Sciences of the United States of America 115 (6): 1268–73.

Sahtoe, Danny D., Florian Praetorius, Alexis Courbet, Yang Hsia, Basile I. M. Wicky, Natasha I. Edman, Lauren M. Miller, et al. 2022. “Reconfigurable Asymmetric Protein Assemblies through Implicit Negative Design.” Science 375 (6578): eabj7662.

Schneidman-Duhovny, Dina, Michal Hammel, John A. Tainer, and Andrej Sali. 2013. “Accurate SAXS Profile Computation and Its Assessment by Contrast Variation Experiments.” Biophysical Journal 105 (4): 962–74.

Schneidman-Duhovny, Dina, Michal Hammel, John A. Tainer, and Andrej Sali. 2013. “Accurate SAXS Profile. 2016. “FoXS, FoXSDock and MultiFoXS: Single-State and Multi-State Structural Modeling of Proteins and Their Complexes Based on SAXS Profiles.” Nucleic Acids Research 44 (W1): W424–29.

Silva, Daniel-Adriano, Shawn Yu, Umut Y. Ulge, Jamie B. Spangler, Kevin M. Jude, Carlos Labão-Almeida, Lestat R. Ali, et al. 2019. “De Novo Design of Potent and Selective Mimics of IL-2 and IL-15.” Nature 565 (7738): 186–91.

Song, Joon Young, Won Suk Choi, Jung Yeon Heo, Jin Soo Lee, Dong Sik Jung, Shin-Woo Kim, Kyung-Hwa Park, et al. 2022. “Safety and Immunogenicity of a SARS-CoV-2 Recombinant Protein Nanoparticle Vaccine (GBP510) Adjuvanted with AS03: A Randomised, Placebo-Controlled, Observer-Blinded Phase 1/2 Trial.” EClinicalMedicine 51 (September): 101569.

Suloway, Christian, James Pulokas, Denis Fellmann, Anchi Cheng, Francisco Guerra, Joel Quispe, Scott Stagg, Clinton S. Potter, and Bridget Carragher. 2005. “Automated Molecular Microscopy: The New Leginon System.” Journal of Structural Biology 151 (1): 41–60.

Tetter, Stephan, Naohiro Terasaka, Angela Steinauer, Richard J. Bingham, Sam Clark, Andrew J. P. Scott, Nikesh Patel, et al. 2021. “Evolution of a Virus-like Architecture and Packaging Mechanism in a Repurposed Bacterial Protein.” Science 372 (6547): 1220–24.

Ueda, George, Aleksandar Antanasijevic, Jorge A. Fallas, William Sheffler, Jeffrey Copps, Daniel Ellis, Geoffrey B. Hutchinson, et al. 2020a. “Tailored Design of Protein Nanoparticle Scaffolds for Multivalent Presentation of Viral Glycoprotein Antigens.” eLife 9 (August). https://doi.org/10.7554/eLife.57659.

Ueda, George, Aleksandar Antanasijevic, Jorge A. Fallas, William Sheffler, Jeffrey Copps, Daniel Ellis, Geoffrey B. Hutchinson, et al. 2020b. “Tailored Design of Protein Nanoparticle Scaffolds for Multivalent Presentation of Viral Glycoprotein Antigens.” eLife 9 (August). https://doi.org/10.7554/eLife.57659.

Uhlén, Mathias, Max J. Karlsson, Andreas Hober, Anne-Sophie Svensson, Julia Scheffel, David Kotol, Wen Zhong, et al. 2019. “The Human Secretome.” Science Signaling 12 (609): eaaz0274.

UniProt Consortium. 2021. “UniProt: The Universal Protein Knowledgebase in 2021.” Nucleic Acids Research 49 (D1): D480–89.

Veesler, David, Karolina Cupelli, Markus Burger, Peter Gräber, Thilo Stehle, and John E. Johnson. 2014. “Single-Particle EM Reveals Plasticity of Interactions between the Adenovirus Penton Base and Integrin αVβ3.” Proceedings of the National Academy of Sciences of the United States of America 111 (24): 8815–19.

Vieira Gomes, Antonio Milton, Talita Souza Carmo, Lucas Silva Carvalho, Frederico Mendonça Bahia, and Nádia Skorupa Parachin. 2018. “Comparison of Yeasts as Hosts for Recombinant Protein Production.” Microorganisms 6 (2). https://doi.org/10.3390/microorganisms6020038.

Walls, Alexandra C., Brooke Fiala, Alexandra Schäfer, Samuel Wrenn, Minh N. Pham, Michael Murphy, Longping V. Tse, et al. 2020. “Elicitation of Potent Neutralizing Antibody Responses by Designed Protein Nanoparticle Vaccines for SARS-CoV-2.” Cell 183 (5): 1367–82.e17.

Walsh, Gary. 2018. “Biopharmaceutical Benchmarks 2018.” Nature Biotechnology 36 (12): 1136–45.

Wang, Dan, Phillip W. L. Tai, and Guangping Gao. 2019. “Adeno-Associated Virus Vector as a Platform for Gene Therapy Delivery.” Nature Reviews. Drug Discovery 18 (5): 358–78.

Wang, Ray Yu-Ruei, Yifan Song, Benjamin A. Barad, Yifan Cheng, James S. Fraser, and Frank DiMaio. 2016. “Automated Structure Refinement of Macromolecular Assemblies from Cryo-EM Maps Using Rosetta.” eLife 5 (September). https://doi.org/10.7554/eLife.17219.

Williams, Christopher J., Jeffrey J. Headd, Nigel W. Moriarty, Michael G. Prisant, Lizbeth L. Videau, Lindsay N. Deis, Vishal Verma, et al. 2018. “MolProbity: More and Better Reference Data for Improved All-Atom Structure Validation.” Protein Science: A Publication of the Protein Society 27 (1): 293–315.

Winn, Martyn D., Charles C. Ballard, Kevin D. Cowtan, Eleanor J. Dodson, Paul Emsley, Phil R. Evans, Ronan M. Keegan, et al. 2011. “Overview of the CCP4 Suite and Current Developments.” Acta Crystallographica. Section D, Biological Crystallography 67 (Pt 4): 235–42.

Wittrup, K. D. 1995. “Disulfide Bond Formation and Eukaryotic Secretory Productivity.” Current Opinion in Biotechnology 6 (2): 203–8.

Xiao, Su, Joseph Shiloach, and Michael J. Betenbaugh. 2014. “Engineering Cells to Improve Protein Expression.” Current Opinion in Structural Biology 26 (June): 32–38.

Xu, Xun, Harish Nagarajan, Nathan E. Lewis, Shengkai Pan, Zhiming Cai, Xin Liu, Wenbin Chen, et al. 2011. “The Genomic Sequence of the Chinese Hamster Ovary (CHO)-K1 Cell Line.” Nature Biotechnology 29 (8): 735–41.

Xu, Ziyang, Megan C. Wise, Neethu Chokkalingam, Susanne Walker, Edgar Tello-Ruiz, Sarah T. C. Elliott, Alfredo Perales-Puchalt, et al. 2020. “In Vivo Assembly of Nanoparticles Achieved through Synergy of Structure-Based Protein Engineering and Synthetic DNA Generates Enhanced Adaptive Immunity.” Advancement of Science 7 (8): 1902802.

Zhang, Kai. 2016. “Gctf: Real-Time CTF Determination and Correction.” Journal of Structural Biology 193 (1): 1–12.

Zhang, Shuguang. 2003. “Fabrication of Novel Biomaterials through Molecular Self-Assembly.” Nature Biotechnology 21 (10): 1171–78.

Zheng, Shawn Q., Eugene Palovcak, Jean-Paul Armache, Kliment A. Verba, Yifan Cheng, and David A. Agard. 2017. “MotionCor2: Anisotropic Correction of Beam-Induced Motion for Improved Cryo-Electron Microscopy.” Nature Methods 14 (4): 331–32.

Zivanov, Jasenko, Takanori Nakane, Björn O. Forsberg, Dari Kimanius, Wim Jh Hagen, Erik Lindahl, and Sjors Hw Scheres. 2018. “New Tools for Automated High-Resolution Cryo-EM Structure Determination in RELION-3.” eLife 7 (November). https://doi.org/10.7554/eLife.42166.

