## Supplementary material for "Improving the secretion of designed protein assemblies through negative design of cryptic transmembrane domains": design script

#############
cage_design.sh
#############

```
 #if input pdb doesn't already exist, make it from the original dock.pickle file
if [[ ! -e input/scaffolds/${dock}.pdb ]]; then
	PYTHONPATH="/path/to/your/rpxdock" /path/to//your/.conda/envs/rpxdock/bin/python
input/extract_scaffold.py ${dock} ${rpx_pickle_path}
	sleep 3
fi

file_input="input/scaffolds/${dock}.pdb"
mkdir -p output/${sym}/${comp1}/${dock}
outpath="output/${sym}/${comp1}/${dock}"

symfile="/path/to/your/symdefiles//${sym}.sym"
if [[ ${sym} == "T3" ]]; then symdof1="JAC1"
elif [[ ${sym} == "O3" ]]; then symdof1="JTP1"
elif [[ ${sym} == "I3" ]]; then symdof1="JCT00"
else echo "not a valid  sym"; exit ; fi
symdof2="${symdof1}"

#run Rosetta
if [[ ! -e ${outpath}/score.sc ]]; then
      /path/to/your/rosetta/main/source/bin/rosetta_scripts.static.linuxgccrelease \
      -dunbrack_prob_buried 0.8 -dunbrack_prob_nonburied 0.8 -dunbrack_prob_buried_semi 0.8
-dunbrack_prob_nonburied_semi 0.8 \
            -out::file::pdb_comments \
            -parser:protocol xml/cage_design.xml \
            -s      ${file_input} \
            -native ${file_input} \
            -beta \
            -nstruct 20 \
            -parser:script_vars sym="${symfile}" symdof1="${symdof1}" symdof2="${symdof2}"
outpath="${outpath}" \
            -overwrite 1 \
            -unmute all \
            -out:chtimestamp 1 \
            -out:suffix "" \
            -out:level 300 \
            -out::path::all ${outpath}/ \
      -output_only_asymmetric_unit true \
      -failed_job_exception False \
      -mute core.select.residue_selector.SecondaryStructureSelector \
            > ${outpath}/${sym}_${dock}_${comp1}_${comp2}.log
else
      echo "${outpath}/score.sc exists!"
fi
```

##############
cage_design.xml
##############

```
<ROSETTASCRIPTS>
	<SCOREFXNS>
		<ScoreFunction name="sfxn_hard" weights="beta" symmetric="1" >
	<Reweight scoretype="approximate_buried_unsat_penalty" weight="5.0" />
	<Set approximate_buried_unsat_penalty_hbond_energy_threshold="-0.25" />
	<Set approximate_buried_unsat_penalty_burial_atomic_depth="4.0" />
	<Set approximate_buried_unsat_penalty_assume_const_backbone="true" />
	<Reweight scoretype="lk_ball" weight="0" />
	<Reweight scoretype="lk_ball_iso" weight="0" />
	<Reweight scoretype="lk_ball_wtd" weight="0" />
	<Reweight scoretype="lk_ball_bridge" weight="0" />
	<Reweight scoretype="lk_ball_bridge_uncpl" weight="0" />
		</ScoreFunction>

		<ScoreFunction name="sfxn_soft" weights="beta_soft" symmetric="1" >
	<Reweight scoretype="approximate_buried_unsat_penalty" weight="5.0" />
	<Set approximate_buried_unsat_penalty_hbond_energy_threshold="-0.25" />
	<Set approximate_buried_unsat_penalty_burial_atomic_depth="4.0" />
	<Set approximate_buried_unsat_penalty_assume_const_backbone="true" />
	<Reweight scoretype="lk_ball" weight="0" />
	<Reweight scoretype="lk_ball_iso" weight="0" />
	<Reweight scoretype="lk_ball_wtd" weight="0" />
	<Reweight scoretype="lk_ball_bridge" weight="0" />
	<Reweight scoretype="lk_ball_bridge_uncpl" weight="0" />
		</ScoreFunction>

		<ScoreFunction name="sfxn_up_ele" weights="beta" symmetric="1" >
	<Reweight scoretype="approximate_buried_unsat_penalty" weight="5.0" />
	<Set approximate_buried_unsat_penalty_hbond_energy_threshold="-0.25" />
	<Set approximate_buried_unsat_penalty_burial_atomic_depth="4.0" />
	<Set approximate_buried_unsat_penalty_assume_const_backbone="true" />
	<Reweight scoretype="lk_ball" weight="0" />
	<Reweight scoretype="lk_ball_iso" weight="0" />
	<Reweight scoretype="lk_ball_wtd" weight="0" />
	<Reweight scoretype="lk_ball_bridge" weight="0" />
	<Reweight scoretype="lk_ball_bridge_uncpl" weight="0" />
			<Reweight scoretype="fa_elec" weight="1.4" />
			<Reweight scoretype="hbond_sc" weight="2.0" />
		</ScoreFunction>

		<ScoreFunction name="sfxn_clean" weights="beta" symmetric="1" />
	</SCOREFXNS>

	<TASKOPERATIONS>
		//detects residues between building blocks
```

```
		<BuildingBlockInterface name="design_bbi_1comp" multicomp="0"
sym_dof_names="%%symdof1%%" nsub_bblock="%%nsub_bb%%" fa_rep_cut="3.0" contact_dist="10"
bblock_dist="5" />
		//selects surface and boundary residues of building block
		<SelectBySASA name="bb_surf" mode="sc" state="monomer" core_asa="0" surface_asa="0"
core="0" boundary="1" surface="1" verbose="0" />
		//detects if any residues are not the same as native input
		<RestrictNativeResidues name="nonnative" prevent_repacking="1" invert="0" />
	</TASKOPERATIONS>

	<RESIDUE_SELECTORS>
		//selects CPG residues
		<ResidueName name="CPG" residue_name3="CYS,PRO,GLY" />
			<Not name="not_CPG" selector="CPG" />

		//interface selection
		<Task name="design_bbi_selector" designable="true" task_operations="design_bbi_1comp" />
		<Task name="bb_surf_selector" designable="true" task_operations="bb_surf" />
			<And name="design_int_no_nonnative"
selectors="design_bbi_selector,bb_surf_selector,not_CPG" />
		<Task name="nonnative_selector" designable="true" task_operations="nonnative" />

		//set design_resis
		<Or name="design_resis" selectors="design_int_no_nonnative,nonnative_selector" /> //selects
full interface, including preserved residues
		<Neighborhood name="pack_resis" selector="design_resis" distance="5"
include_focus_in_subset="false" />
		<Or name="design_and_pack_resis" selectors="design_resis,pack_resis" />
			<Not name="lock_resis" selector="design_and_pack_resis" />

		//layer design
		<Layer name="core" use_sidechain_neighbors="true" core_cutoff="4.9" surface_cutoff="2.7"
sc_neighbor_dist_exponent="0.7" select_core="true" />
		<Layer name="bdry" use_sidechain_neighbors="true" core_cutoff="4.9" surface_cutoff="2.7"
sc_neighbor_dist_exponent="0.7" select_boundary="true" />
		<Layer name="surf" use_sidechain_neighbors="true" core_cutoff="4.9" surface_cutoff="2.7"
sc_neighbor_dist_exponent="0.7" select_surface="true" />
		<SecondaryStructure name="helix" ss="H" overlap="0" minH="3" minE="3" use_dssp="true"
include_terminal_loops="false" />
		<SecondaryStructure name="sheet" ss="E" overlap="0" minH="3" minE="3" use_dssp="true"
include_terminal_loops="false" />
		<SecondaryStructure name="loop"	ss="L" overlap="0" minH="3" minE="3" use_dssp="true"
include_terminal_loops="true" />
			<And name="design_surf" selectors="surf,design_resis" />
			<And name="design_bdry" selectors="bdry,design_resis" />
			<And name="design_core" selectors="core,design_resis" />

		//for revert
```

```
			<Neighborhood name="revert_pack_resis" selector="nonnative_selector" distance="5"
include_focus_in_subset="false" />
			<Or name="revert_design_and_pack_resis" selectors="nonnative_selector,revert_pack_resis"
/>
				<Not name="revert_lock_resis" selector="revert_design_and_pack_resis" />

			//for filters
			<Or name="design_core_bdry" selectors="design_core,design_bdry" />

	//for simple_metrics
	<Chain name="chainA" chains="A" />
	<Chain name="chainB" chains="B" />
	<And name="int_resis_chainA" selectors="design_resis,chainA" />
	<And name="int_resis_chainB" selectors="design_resis,chainB" />

	//for contact_mol_surf
	<SymmetricalResidue name="int_resis_chainB_sym" selector="int_resis_chainB" />

	//for ala_mut
	<ResidueName name="very_polar_resis" residue_name3="ASP,GLU,ASN,GLN,HIS,LYS,ARG" />
	<And name="polar_design_resis" selectors="design_resis,very_polar_resis" />
	  <Not name="not_polar_design_resis" selector="polar_design_resis" />
	  </RESIDUE_SELECTORS>

	  <TASKOPERATIONS>
			<IncludeCurrent name="ic" /> //includes input pdb's rotamers
			<LimitAromaChi2 name="limitaro" chi2max="110" chi2min="70" /> //disallow extreme aromatic
rotamers
			<RestrictToRepacking name="repack_only" /> //for minimize/repack
			<ExtraRotamersGeneric name="ex1_ex2" ex1="1" ex2="0" ex2aro="1" /> //use ex1 and ex2
rotamers

			//layer design
			<DesignRestrictions name="layer_design" >
				<Action selector_logic="surf AND ( helix OR sheet )" aas="DEKRNQST" />
				<Action selector_logic="surf AND loop" aas="DEKRNQSTGP" />
				<Action selector_logic="bdry AND helix" aas="ILVAFMDEKRNQSTWY" />
				<Action selector_logic="bdry AND sheet" aas="ILVAFDEKRNQSTWY" />
				<Action selector_logic="bdry AND loop" aas="ILVAFMDEKRNQSTGPWY" />
				<Action selector_logic="core AND helix" aas="ILVAFM" />
				<Action selector_logic="core AND sheet" aas="ILVAF" />
				<Action selector_logic="core AND loop" aas="ILVAFMGP" />
			</DesignRestrictions>

			//setup tasks
			<OperateOnResidueSubset name="design_task" selector="design_resis" >
				<RestrictAbsentCanonicalAASRLT aas="FWYHCPGDENQKRMILVSTA" />
</OperateOnResidueSubset>
			<OperateOnResidueSubset name="pack_task" selector="pack_resis" >
```

```
			<RestrictToRepackingRLT/> </OperateOnResidueSubset>
		<OperateOnResidueSubset name="lock_task" selector="lock_resis" >
			<PreventRepackingRLT/> </OperateOnResidueSubset>

		//for surf_design
		<OperateOnResidueSubset name="pack_core_task" selector="core" >
			<RestrictToRepackingRLT/> </OperateOnResidueSubset>

		//for revert
		<JointSequence name="revert" use_current="true" use_native="true" use_natro="true" />
		<OperateOnResidueSubset name="revert_pack_task" selector="revert_pack_resis" >
			<RestrictToRepackingRLT/> </OperateOnResidueSubset>
		<OperateOnResidueSubset name="revert_lock_task" selector="revert_lock_resis" >
			<PreventRepackingRLT/> </OperateOnResidueSubset>

		//setup for ala_mut
	<OperateOnResidueSubset name="polar_design_resis_task" selector="polar_design_resis" >
	  <RestrictAbsentCanonicalAASRLT aas="A" /> </OperateOnResidueSubset>
	<OperateOnResidueSubset name="lock_polar_design_resis_task" selector="not_polar_design_resis" >
	  <PreventRepackingRLT/> </OperateOnResidueSubset>
	  </TASKOPERATIONS>

	  <MOVERS>
		//setting up the degreaser
		<SecretionOptimizationMover name="degreaseA" max_mutations="1" chain_to_degrease="A"
dG_ins_threshold="3.5" dump_singles="false" dump_multis="false" score_tolerance="15"
scorefxn="sfxn_clean" aas_allowed="DEKRQNSTY" />

		//regen sym and sample without a/r; 1-component
		<SymDofMover name="gen_docked_config_no_transforms_1comp_mover"
symm_file="%%sym%%" sym_dof_names="%%symdof1%%"
			flip_input_about_axes="0,0" />

		<SymPackRotamersMover name="design_hard" scorefxn="sfxn_hard"
task_operations="layer_design,pack_task,lock_task,limitaro,ic,ex1_ex2" />
		<SymPackRotamersMover name="design_surf" scorefxn="sfxn_up_ele"
task_operations="layer_design,pack_core_task,pack_task,lock_task,limitaro,ic,ex1_ex2" />

		<TaskAwareSymMinMover name="min_clean" scorefxn="sfxn_clean" bb="0" chi="1" rb="1"
task_operations="pack_task,lock_task" />
		<SymPackRotamersMover name="repack_clean" scorefxn="sfxn_clean"
task_operations="pack_task,lock_task,limitaro,ic,ex1_ex2,repack_only" />

		<ParsedProtocol name="min_repack_min" >
			<Add mover="min_clean" />
			<Add mover="repack_clean" />
			<Add mover="min_clean" />
		</ParsedProtocol>
```

```
			<TaskAwareSymMinMover name="min_clean_no_rb" scorefxn="sfxn_clean" bb="0" chi="1"
rb="0" task_operations="pack_task,lock_task,repack_only" /> //this is for ddG
			<AddResidueLabel name="label_design_resis" residue_selector="design_resis"
label="design_resis" />

	//mutate polar residues at interface to Ala before calculating some filters
	<SymPackRotamersMover name="ala_mut" scorefxn="sfxn_clean"
task_operations="polar_design_resis_task,lock_polar_design_resis_task" />
		</MOVERS>

		<FILTERS>
			<ClashCheck name="clash_check_1comp" nsub_bblock="3" verbose="1" write2pdb="1"
cutoff="2" confidence="1" />

			<ShapeComplementarity name="sc1_1comp" multicomp="0" verbose="1" min_sc="0.5"
sym_dof_name="%%symdof1%%" write_int_area="1" write_median_dist="1" confidence="1" />
			<ShapeComplementarity name="sc2_1comp" multicomp="0" verbose="1" min_sc="0.5"
sym_dof_name="%%symdof2%%" write_int_area="1" write_median_dist="1" confidence="1" />

			<ShapeComplementarity name="sc1_1comp_hpc" multicomp="0" verbose="1" min_sc="0.5"
sym_dof_name="%%symdof1%%" write_int_area="1" write_median_dist="1" confidence="1" />
			<ShapeComplementarity name="sc2_1comp_hpc" multicomp="0" verbose="1" min_sc="0.5"
sym_dof_name="%%symdof2%%" write_int_area="1" write_median_dist="1" confidence="1" />
	<MoveBeforeFilter name="sc1_hpc" mover="ala_mut" filter="sc1_1comp_hpc" confidence="1" />
	<MoveBeforeFilter name="sc2_hpc" mover="ala_mut" filter="sc2_1comp_hpc" confidence="1" />

			<Sasa name="sasa_1comp" threshold="300" upper_threshold="750" jump="1" hydrophobic="0"
polar="0" confidence="1" />
			<Sasa name="sasa_1comp_hpc" threshold="300" upper_threshold="750" jump="1"
hydrophobic="0" polar="0" confidence="1" />
		<MoveBeforeFilter name="sasa_hpc" mover="ala_mut" filter="sasa_1comp_hpc" confidence="1" />

	<ContactMolecularSurface name="contact_mol_surf" min_interface="0" distance_weight="1.0"
verbose="1" quick="0" target_selector="int_resis_chainA" binder_selector="int_resis_chainB_sym"
confidence="0" />

	<Ddg name="ddG" repeats="1" extreme_value_removal="0" translate_by="1000" scorefxn="sfxn_clean"
task_operations="pack_task,lock_task,repack_only,ic,ex1_ex2" repack="1" threshold="0"
relax_mover="min_clean_no_rb" repack_bound="0" relax_bound="0" repack_unbound="1" relax_unbound="1"
confidence="0" />
	<MoveBeforeFilter name="ddG_hpc" mover="ala_mut" filter="ddG" confidence="0" />

			<SecondaryStructureCount name="ss_count_1comp" filter_helix_sheet="1"
num_helix_sheet="2" min_helix_length="4" min_sheet_length="3" min_loop_length="1" return_total="1"
confidence="1" residue_selector="design_core_bdry" min_element_resis="3" />

			<OligomericAverageDegree name="avg_deg_1comp" threshold="0" multicomp="0"
distance_threshold="10" sym_dof_names="%%symdof1%%" task_operations="pack_task,lock_task"
confidence="0" />
```

```
			<Mutations name="mutations" rate_threshold="0.0" mutation_threshold="60"
report_mutations="1" verbose="1" write2pdb="1" task_operations="nonnative" confidence="0" />
			<ResidueCount name="AlaCount" residue_types="ALA"
residue_selector="design_resis" confidence="0" />
			<ResidueCount name="MetCount" residue_types="MET"
residue_selector="design_resis" confidence="0" />
			<ResidueCount name="HPcCount" residue_types="VAL,LEU,ILE"
residue_selector="design_resis" confidence="0" />
			<ResidueCount name="AroCount" residue_types="TRP,PHE,TYR,HIS"
residue_selector="design_resis" confidence="0" />

			//cage regeneration terms
			<GetRBDOFValues name="%%symdof1%%_1_disp"  sym_dof_name="%%symdof1%%"
get_disp="1"  get_init_value="1" />
			<GetRBDOFValues name="%%symdof1%%_1_angle" sym_dof_name="%%symdof1%%"
get_angle="1" get_init_value="1" />
			<GetRBDOFValues name="%%symdof2%%_2_disp"  sym_dof_name="%%symdof2%%"
get_disp="1"  get_init_value="1" />
			<GetRBDOFValues name="%%symdof2%%_2_angle" sym_dof_name="%%symdof2%%"
get_angle="1" get_init_value="1" />
		</FILTERS>

  //FOR SIMPLE METRICS
  <SIMPLE_METRICS>
    <SelectedResidueCountMetric name="chainA_len" residue_selector="chainA" custom_type="chainA_" />
    <SelectedResidueCountMetric name="chainB_len" residue_selector="chainB" custom_type="chainB_" />
    <SelectedResidueCountMetric name="chnA_int_resis" residue_selector="int_resis_chainA"
custom_type="int_resi_chnA_" />
    <SelectedResidueCountMetric name="chnB_int_resis" residue_selector="int_resis_chainB"
custom_type="int_resi_chnB_" />
		</SIMPLE_METRICS>

  <MOVERS>
    <RunSimpleMetrics name="run_metrics" metrics="chainA_len,chainB_len" />
  </MOVERS>

		<PROTOCOLS>
			//generate docked configuration
			<Add mover_name="gen_docked_config_no_transforms_1comp_mover" />

			//pre-design filters
			<Add filter_name="clash_check_1comp" />
			<Add filter_name="sasa_1comp" />
			<Add filter_name="ss_count_1comp" />

			//layer design
			<Add mover_name="label_design_resis" />
			<Add mover_name="design_hard" />
```

```
		<Add mover_name="min_repack_min" />

		//boundary/surface design with up_ele
		<Add mover_name="design_surf" />
		<Add mover_name="min_repack_min" />

		//call the degreaser
		<Add mover_name="degreaseA" />

		//filters
		<Add filter_name="clash_check_1comp" />
		<Add filter_name="sc1_1comp" />
		<Add filter_name="sc2_1comp" />
		<Add filter_name="sc1_hpc" />
		<Add filter_name="sc2_hpc" />
		<Add filter_name="sasa_1comp" />
		<Add filter_name="sasa_hpc" />
		<Add filter_name="contact_mol_surf" />
		<Add filter_name="ddG" />
		<Add filter_name="ddG_hpc" />
		<Add filter_name="ss_count_1comp" />
		<Add filter_name="avg_deg_1comp" />
		<Add filter_name="mutations" />
		<Add filter_name="AlaCount" />
		<Add filter_name="MetCount" />
		<Add filter_name="HPcCount" />
		<Add filter_name="AroCount" />

		<Add mover_name="run_metrics"/>

		//recalculate angle and displacement
		<Add filter_name="%%symdof1%%_1_disp" />
		<Add filter_name="%%symdof1%%_1_angle" />
		<Add filter_name="%%symdof2%%_2_disp" />
		<Add filter_name="%%symdof2%%_2_angle" />
	</PROTOCOLS>

	<OUTPUT scorefxn="sfxn_clean" />
</ROSETTASCRIPTS>
```
